# Nervous system-wide single-cell morphology atlas of excitatory and inhibitory neurons in larval zebrafish

**DOI:** 10.1101/2025.06.06.658008

**Authors:** Xu-Fei Du, Zhi-Feng Yue, Ming-Quan Chen, Wan-Lan Li, Tian-Lun Chen, Han-Yang Hu, Hong-Li Wan, Ting-Ting Zhao, Yong-Wei Zhong, Xin-Yu Ning, Xiu-Dan Zheng, Hong-An Ren, Rui-Qi Wang, Rong Zhao, Xiao-Lan Peng, Zhi-Ming Jia, Li-Jun Chen, Chen-Xi Jin, Jia-Wen Huang, Wei Deng, Liu-Qin Qian, Li Wang, Mei-Yu Zheng, Wei Zhang, Xiao-Yu Shen, Xiu-Lian Shen, Xiao-Ying Qiu, Qi-Meng Zhao, Dan-Yang Li, Si-Jia Wang, Yu-Chen Gong, Yan Li, Yu Mu, Peng Ji, Jie He, Yu-Fan Wang, Jiu-Lin Du

**Author notes:** These authors contributed equally (Xu-Fei Du, Zhi-Feng Yue). These authors jointly supervised this work: Xu-Fei Du, Jiu-Lin Du.

## Abstract

Single-neuron morphology mapping is fundamental for deciphering brain architecture and function, yet in vertebrates it remains limited in spatial coverage and cell-type information. Here we present a nervous system-wide single-neuron morphology atlas of larval zebrafish with annotated excitatory/inhibitory identity and dendrite-axon polarity. The atlas comprises >20,000 reconstructed neurons from >13,400 animals, spanning the central and peripheral nervous system and representing ∼24.5% of the total excitatory and inhibitory neurons in the brain, all integrated into a common physical space with neuroanatomical parcellations. We define >500 excitatory and inhibitory neuronal morphotypes and construct putative inter-cell-type and interregional connectomes. Projection and morphotype analyses reveal distinct laterality and directionality biases and modular innervation patterns in excitatory and inhibitory neurons, while network analyses uncover fundamental connectivity motifs, hubs, and sensorimotor pathways. Accessible through custom-built interactive platform ZExplorer, this atlas provides a foundational resource for multimodal data integration, circuit analysis, computational modeling, and hypothesis-driven research.

## INTRODUCTION

The nervous system, the most complex system in the body, controls how animals sense, represent, and respond to both external and internal worlds through its intrinsic organization and connections with other body parts. This control depends on a myriad of elements and their interactions in this complex system. Since Santiago Ramón y Cajal first identified the neuron as the fundamental unit of the nervous system and proposed the concept of unidirectional information flow through neural circuits,^1^ it has remained a challenge to understand how the nervous system is organized, both structurally and functionally, and particularly in terms of the relationship between structure and function.^2,3^ To tackle this challenge, a critical step is to elucidate the connectome at meso- and microscopic levels, i.e., the comprehensive map of neuron types and their connections.

Over the past decade, advances in light microscopy (LM) and electron microscopy (EM) techniques have enabled extensive efforts to map the connectome of the nervous system by acquiring large-scale structural datasets at cellular or synaptic resolution across various model organisms. In invertebrates, whole-body EM connectomes have been reconstructed for adult nematode *Caenorhabditis elegans*.^4–6^ For *Drosophila melanogaster*, LM-based reconstructions have captured the single-cell morphologies of ∼16,000 neurons,^7^ and full synaptic-resolution EM reconstructions of all neurons in the central brain have been achieved.^8–11^ Given the distinct organization of vertebrate and invertebrate nervous systems, connectome mapping in vertebrates is required for advancing our understanding of vertebrate brains. In recent years, substantial progress has been made in reconstructing single-neuron morphologies in the mouse brain: reconstruction time has been reduced from days to hours per neuron, and the number of reconstructed neurons has increased from dozens to twenty thousand.^12–16^ Nevertheless, due to the large number of neurons (∼70 million in the mouse brain) and their morphological complexity, current efforts primarily focus on mapping brain-wide projections of neurons located in a limited number of specific brain regions. Achieving a comprehensive dataset that captures ‘whole-brain neurons with whole-brain projections’ in higher vertebrates remains a significant challenge.^17–21^

Among vertebrate models, the larval zebrafish (*Danio rerio*) possesses a compact brain, making it ideal for whole-brain connectome mapping. With a scale of one hundred thousand neurons, its brain preserves conserved neurotransmitter types and neuroanatomical regions found in higher vertebrates.^22^ Its optical transparency and a wealth of available genetic tools^23^ make it uniquely suited for live whole-brain structural and functional imaging at single-cell resolution by LM.^24–26^ Its small brain volume (<0.1 mm^3^) also facilitates intact-tissue three-dimensional (3D) *in situ* RNA sequencing^27,28^ and whole-brain serial-section EM.^29–33^ Integrating LM and EM through cross-modality registration^34,35^ opens the door to linking functional neuronal activity with structural synaptic connectivity. Based on this understanding, the larval zebrafish is regarded as a promising model for uncovering nervous system organization in vertebrates.

Recent advances in LM- and EM-based reconstructions have paved the way for constructing the zebrafish connectome, yet existing datasets remain limited in scope and completeness. A pioneering LM-based zebrafish dataset captured around 2,000 brain neurons but lacked type identities.^36^ In parallel, EM studies have generated increasingly large-scale datasets,^29–33^ moving toward the complete reconstruction of neuronal morphology and synaptic connectivity. However, interpreting these microscale connectomes requires neuronal type information, which is absent^29–32^ or limited^33^ in current EM datasets. A feasible strategy is to assign the neuronal type identity from LM-based single-neuron morphology atlases to EM-reconstructed neurons via morphology-based matching.^33,37^ Therefore, LM-based reconstructions with neuronal type information are essential for annotating and interpreting emerging complete EM connectomes.

In the present study, we built a single-cell morphology atlas of excitatory (E) and inhibitory (I) neurons across the nervous system of larval zebrafish at 6 days post-fertilization (dpf). First, we established a body-wide common physical space (CPSv1) for multimodal data integration, upon which we parcellated the entire brain, anterior spinal cord, and peripheral ganglia in 3D. We then mapped the distribution of E and I neurons (i.e., cytoarchitecture) and reconstructed the morphologies of 20,011 individual neurons across the central and peripheral nervous systems (CNS and PNS) from 13,429 individual animals. The reconstructed morphology dataset covered ∼24.5% of the counted total E and I neurons in the brain. At the mesoscale, we annotated neurons with dendrite-axon polarity and classified them into over 500 morphotypes, uncovered projection design rules, and constructed a morphotype-based putative connectome for circuit mining. At the macroscale, we constructed directed and weighted E and I interregional connectomes based on neuron projectomes, and identified connectivity motifs, network hubs, and sensorimotor pathways across the nervous system. All reference transgenic patterns, region parcellations, cytoarchitectures, and single-neuron morphology reconstructions are available via a custom-built online platform, ZExplorer (https://zebrafish.cn/LM-Atlas/EI). It enables interactive 3D visualization of selected individual elements across datasets within the CPSv1 and supports neuron querying by soma location, projection target, and morphotype, together with analyses of regional soma distribution, axonal projection, and connectivity (Figure S1).

## RESULTS

### Generating a larval zebrafish body-wide CPSv1 with the nervous system parcellated

To build single-neuron morphology atlases, a cellular-resolution 3D reference space is essential for integrating data originating from individual animals. Therefore, we first constructed a larval zebrafish common physical space (CPSv1) via generating a two-channel volumetric average template from 16 live image scans of double transgenic zebrafish *Tg(elavl3:H2B-GCaMP6s);TgBAC(vglut2a:LOXP-DsRed-LOXP-GFP)* at 6 dpf, by using the symmetric groupwise normalization (SyGN) algorithm in the advanced normalization tools (ANTs) (see Methods).^38^ The SyGN method permits concurrent and unbiased generation of an average image for each channel from the corresponding input population through iteration (Figures S2A and S2B). The CPSv1 contains two original reference templates: one with all neurons expressing nuclear GCaMP6s, driven by the pan-neuronal promoter *elavl3*, and another with the majority of excitatory glutamatergic neurons expressing cytosolic DsRed, driven by the glutamatergic neuronal promoter *vglut2a*, thus providing two options for subsequent data integration (Figure 1A). The CPSv1 covers the entire brain, spinal cord segments 1–11, peripheral ganglia, and body structures inferred from autofluorescence signals on the skin, jaw, eye, ear, and fin. Its dimensions are approximately 636 (left to right) × 1,860 (rostral to caudal) × 360 (dorsal to ventral) voxels, with the nervous system occupying ∼64.5 million voxels and the brain ∼41.9 million voxels.

**Figure 1.**
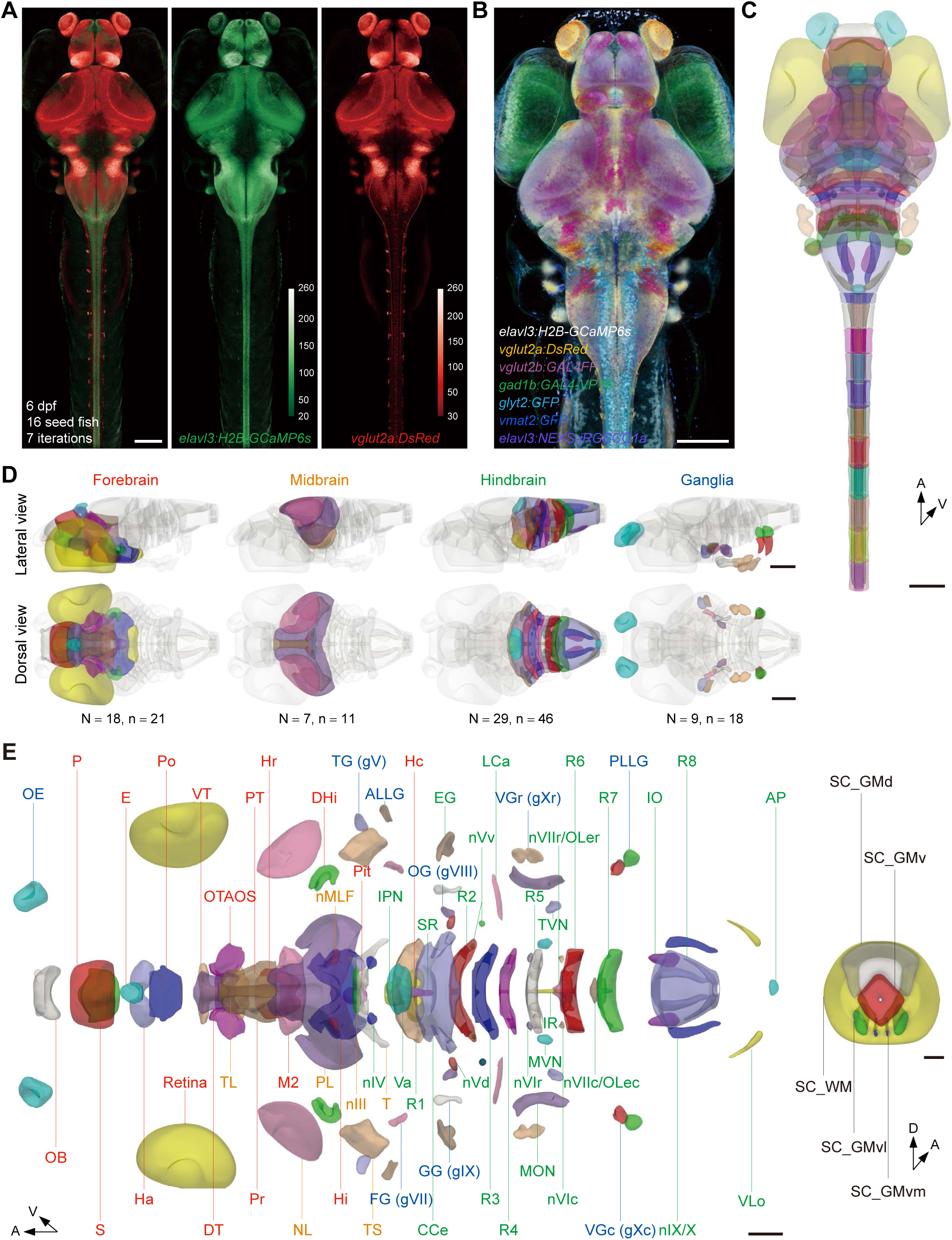
The common physical space (CPSv1) of the larval zebrafish nervous system. (A) Volume rendering of the larval zebrafish CPSv1 with two reference templates, derived from two original gene expression patterns (*elavl3:H2B-GCaMP6s* and *vglut2a:DsRed*) at 6 dpf (dorsal view). 16 seed fish images were iterated 7 times. Color-coded scale bars: image intensity encoded as 8-bit values. (B) Composite volume rendering of 7 transgenic reference templates aligned to the CPSv1 (dorsal view). A partial spinal cord is shown. (C) Dorsal view of 173 non-overlapping, gap-free parcellations (n in D and E) of the nervous system in the CPSv1, color-coded by anatomical regions (N in D and E). (D) Lateral and dorsal views of brain regions in the forebrain, midbrain, hindbrain, and peripheral ganglia. N, number of anatomical regions; n, number of parcellations (bilateral instances counted separately). (E) Dorsal view of the radially disassembled brain and peripheral regions (left) and transverse view of spinal cord regions (right), showing a detailed representation of regions with annotations. Scale bars: 50 μm (E, the transverse view of the spinal cord), 100 μm (A–D and E left). A, anterior; D, dorsal; V, ventral. See also Figures S1 and S2, Video S1, and Table S2.

To establish standard neuroanatomical parcellations within the CPSv1, we registered 702 reference expression patterns (Table S1) to the CPSv1 to delineate brain regions and nuclei. We first generated reference templates for excitatory, inhibitory, and neuromodulatory neuronal populations, which provide important reference information for anatomical annotation. These included an additional excitatory population labeled by *TgBAC(vglut2b:GAL4FF)*, two inhibitory populations labeled by *TgBAC(gad1b:GAL4-VP16)* and *Tg(glyt2:GFP)*, and the monoaminergic population labeled by *Tg(vmat2:GFP)*. We also generated a cytosolic pan-neuronal template using *Tg(elavl3:NEXS-jRGECO1a)* (Figures 1B, S2C and S2D, and Video S1; see Methods). To further enrich neuroanatomical information, we incorporated atlas data from Z-Brain (n = 24),^39^ ZBB (n = 245),^23,40^ and mapzebrain (n = 428),^36,41^ which together comprise a total of 697 expression patterns covering various data types, including enhancer trap, transgene, immunostaining, *in situ* hybridization, and spinal dye backfill (Table S1; https://zebrafish.cn/LM-Atlas/EI). We optimized a unified set of ATNs registration parameters that enabled robust alignment between live scans (our ZExplorer and ZBB), fixed scans (Z-Brain and mapzebrain), and between live and fixed scans (see Methods). Based on mean landmark distances (MLDs) derived from six expression patterns, the inter-atlas registration precision between our ZExplorer and other atlases was 8.40 ± 1.18 μm (mean ± SEM; Figures S2E–S2K; see Methods), close to the average neuronal soma diameter in larval zebrafish (5.80 ± 0.001 μm; see below).

Guided by the integrated expression pattern atlases, we manually parcellated the nervous system into non-overlapping, gap-free, 3D anatomical segments with smooth boundaries (see Methods). Referring to the brain ontologies in Mueller and Wullimann’s early zebrafish brain atlas^22^ and in the Z-Brain,^39^ we annotated 68 hierarchically organized neuroanatomical regions, including 54 brain structures, 5 regions for each segment in the spinal cord, and 9 peripheral ganglia (Figures 1C–1E, and Video S1). Accounting for bilateral pairs of certain regions across hemispheres and for multiple spinal cord segments, we obtained a total of 173 regions, including 78 central brain regions and 77 spinal regions (within 11 vertebrae) in the CNS, as well as 18 peripheral ganglia in the PNS. Relevant regional information and the reference templates used for delineation are summarized in Table S2. These parcellations can be further refined by using multimodal criteria such as neuronal morphology and functional properties in the future.

Taken together, these efforts yield a body-wide larval zebrafish CPSv1 with 3D annotations of the nervous system’s anatomical structures, forming a foundational spatial framework for single-neuron morphology mapping.

### Generating cytoarchitecture atlases of E and I neurons

We then mapped the cytoarchitecture^42^ of E and I neurons, i.e., their spatial distribution and cell counts across the nervous system. At 6 dpf, we imaged four larvae for each relevant transgenic line, in which glutamatergic (*TgBAC(vglut2a:LOXP-DsRed-LOXP-GFP)* and *TgBAC(vglut2b:GAL4FF);Tg(4×nrUAS:GFP)*), GABAergic (*TgBAC(gad1b:GAL4-VP16);Tg(4×nrUAS:GFP))*, glycinergic (*Tg(glyt2:GFP)*), or all neurons (*Tg(elavl3:H2B-GCaMP6s)*) were fluorescently labeled. Labeled neuronal somata in the brain, spinal cord segments 1–11, and peripheral ganglia were manually annotated, and the resulting digitized soma positions were registered to the CPSv1, yielding the cytoarchitecture atlases (Figure 2A and Video S2; see Methods).

**Figure 2.**
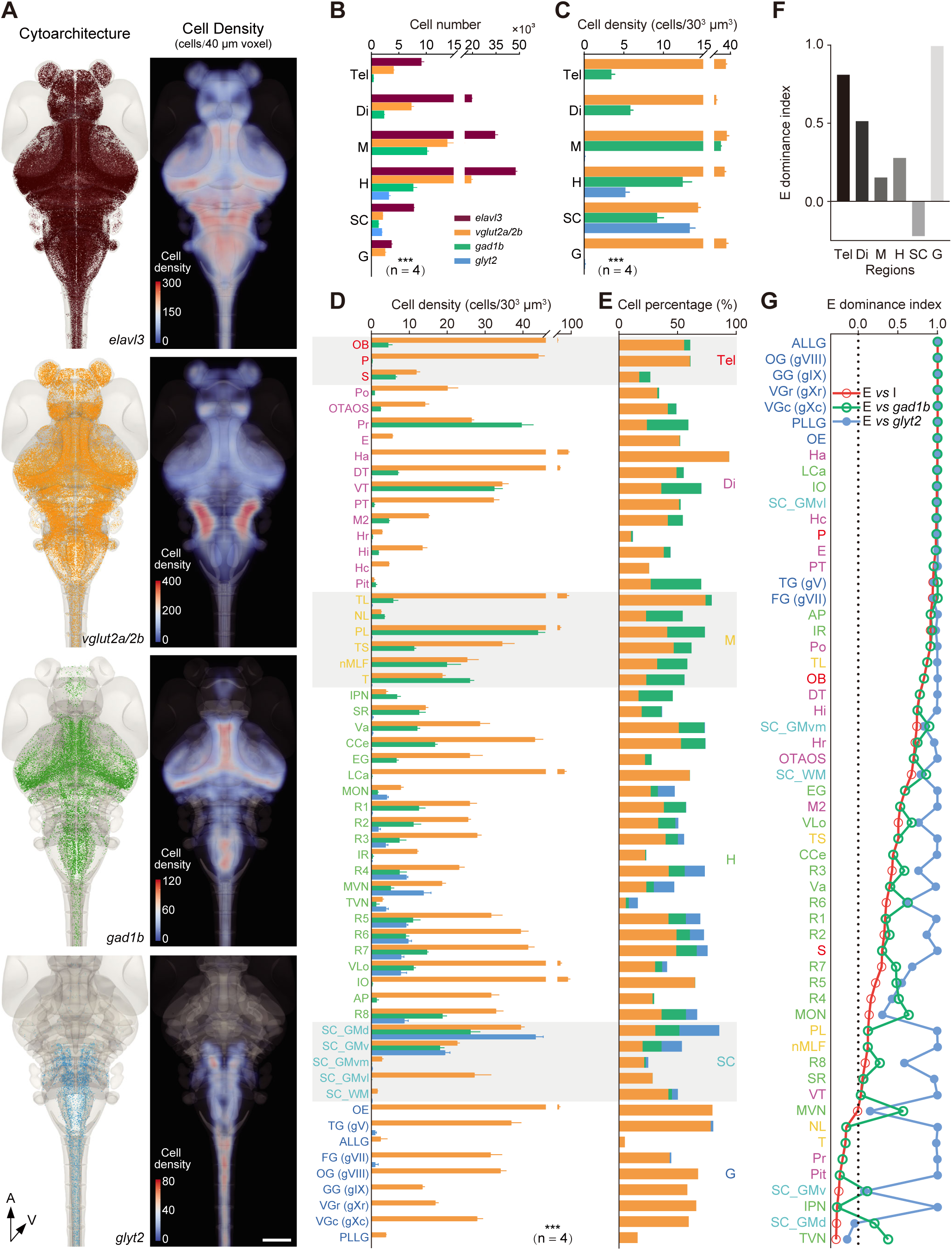
Cytoarchitecture atlases of E and I neurons. (A) Volume rendering of cell somata (left) and cell density (right; number of cells per 40 μm voxel rendered in 3D, shown in dorsal view) for total (*elavl3*), glutamatergic (*vglut2a/2b*), GABAergic (*gad1b*), and glycinergic (*glyt2*) neurons in 6-dpf larvae. Each larva was aligned to the CPSv1. Only a partial spinal cord is shown. (B) Cell number distributions across primary brain subdivisions (Tel, Di, M, and H), spinal cord (SC) segments 1–11, and peripheral ganglia (G), showing global significant differences both among regions and among glutamatergic, GABAergic, and glycinergic neurons. (C and D) Density (per 30^3^ μm^3^) of E and I neurons across 6 high-level (C) and 57 low-level regions (D), showing global significant differences both among regions and among glutamatergic, GABAergic, and glycinergic neurons. (E) Percentage composition of E and I neurons (relative to total *elav3* neurons) across 57 anatomical regions. The cranial motor regions are merged into higher order regions, and the retina is not included. (F and G) E neuron dominance index, calculated as (N_E_−N_I_)/(N_E_+N_I_), across 6 high-level (F) and 57 low-level regions (G). N_E_, number of excitatory neurons; N_I_, number of inhibitory (GABAergic + glycinergic, F), or GABAergic and glycinergic neurons (G). In (G), the indices are ranked in descending order of their values. Red: E *vs*. I; Green, E *vs*. GABAergic; Blue, E *vs*. glycinergic. Scale bar: 100 μm (A). A, anterior; V, ventral. Tel, telencephalon; Di, diencephalon; M, midbrain; H, hindbrain. Region labels on the *y*-axis are color-coded by the attribution of primary subdivisions (D, E, and G). Data are averaged from 4 animals per group. \*\*\**P* < 0.001 (Two-way ANOVA with Tukey’s post hoc multiple comparison test). Error bars represent mean ± SEM. See also Video S2 and Table S2.

Cell counting revealed 122,139 ± 4,171 total neurons (*elavl3*+), ∼49,000 E neurons (*vglut2a*+: 39,495 ± 2,755; *vglut2b*+: 9,646 ± 194), and ∼27,000 I neurons (*gad1b*+: 21,956 ± 625; *glyt2*+: 5,227 ± 398), corresponding to an overall E:I ratio of ∼1.8:1. When mapped onto anatomical parcellations in the CPSv1, *vglut2a/2b*+ excitatory neurons were distributed throughout both the CNS (telencephalon, diencephalon, midbrain, hindbrain, and spinal cord) and PNS (peripheral ganglia), while *gad1b*+ GABAergic neurons were CNS-restricted and *glyt2*+ glycinergic neurons were enriched in the hindbrain and spinal cord (Figure 2B). Restricting the analysis to the brain yielded 110,661 ± 3,937 total neurons (*elavl3*+), 44,517 ± 1,174 excitatory neurons (*vglut2a+* and *vglut2b*+), and 23,832 ± 265.3 inhibitory neurons (*gad1b*+ and *glyt2*+). Notably, all populations increased progressively along the anteroposterior axis from the telencephalon to the hindbrain (Figure 2B).

We next examined neuron densities (per 30^3^ μm^3^) to further characterize spatial variation in neuronal distribution across regions. Distinct patterns emerged across primary nervous system subdivisions: *vglut2a/2b*+ neurons were denser in the telencephalon, midbrain, hindbrain, and peripheral ganglia; *gad1b*+ neuron density peaked in the midbrain; and *glyt2*+ neurons were most abundant in the spinal cord (Figure 2C). Brain region-level analyses revealed significant variability in neuron density both within and between E and I neuronal populations (Figure 2D).

To characterize E-I balance across the nervous system, we first quantified E or I cell percentage relative to the total neuronal population (*elavl3*+). E and I neurons accounted for ∼63% of all neurons, reflecting the presence of other neuronal populations and/or incomplete coverage by transgenic labeling, particularly when using the GAL4/UAS binary reporter system (see Methods), where UAS-transgenes are subject to position effects.^43^ Notably, lower E and I percentages were observed in the preoptic area (Po), superior/inferior raphe (SR and IR), and rostral/caudal hypothalamus (Hr and Hc) (Figure 2E), consistent with the enrichment of neuromodulatory, neuropeptidergic, or neuroendocrine neuronal populations in these areas. We next quantified the E versus I dominance, defined as (N_E_-N_I_)/(N_E_+N_I_), where N denotes cell number. At the subdivision level (Figure 2F), E neurons dominated the brain and peripheral ganglia, with a decreasing gradient of their dominance from the telencephalon to midbrain, while I neurons showed dominance in the spinal cord. At the region level (Figure 2G), sensorimotor-related areas such as the tectal periventricular cell layer (PL) and the nucleus of the medial longitudinal fasciculus (nMLF) displayed balanced E-I compositions, while I dominance emerged in the tectal neuropil layers (NL), pretectum (Pr), tegmentum (T), and interpeduncular nucleus (IPN).

Together, these results show that the numbers of E and I neurons gradually increase along the brain’s anteroposterior axis, likely reflecting the relative sizes of primary subdivisions, while each neuronal population exhibits distinct density patterns across subdivisions and regions, indicating regional specialization in the cellular composition. A prominent spatial trend is the gradual shift from E dominance in the forebrain to I dominance in the spinal cord. This large-scale gradient mirrors vertebrate CNS organization: forebrain regions preferentially integrate and relay excitatory information; midbrain and hindbrain circuits rely on balanced E-I interactions for sensorimotor transformation; and spinal circuits depend heavily on inhibitory neurons to coordinate locomotion.

### Atlasing single-cell morphology of nervous system-wide E and I neurons

To systematically map the morphology of single E and I neurons across the nervous system, we utilized the inherent mosaicism of transient transgenesis for stochastic sparse labeling. A mix of a GAL4 driver (*vglut2a:GAL4FF*, *vglut2b:GAL4FF*, *BACgad1b:GAL4-VP16* or *glyt2:GAL4FF*), a membrane-targeted *UAS* reporter (*14*×*UAS-E1b:tdTomato-CAAX*), and *Tol2* transposase mRNA was injected into fertilized embryos, producing labeling in only one or a few neurons in each GAL4 expression domain. To increase coverage of specific neuronal populations, we additionally used *isl2b* and *atoh7* to label retinal ganglion cells (excitatory), *cbln12* to label cerebellar granule cells (excitatory), and *cpce* to label cerebellar Purkinje cells (inhibitory) (see Methods). Injected embryos also carried the *elavl3:H2B-GCAMP6s* transgene, which provided a reference channel (green fluorescence) for registration. Larvae with suitable sparse labeling at 6 dpf were live-imaged, and the green reference channel was used to register to the CPSv1 via a two-step rigid-affine and nonlinear ANTs transformation. We used the red morphology channel for semi-manual neuron reconstruction and aligned individual digitized neuronal skeletons to the CPSv1 using the two-step transformation matrices (Figure 3A and Video S3; see Methods).

**Figure 3.**
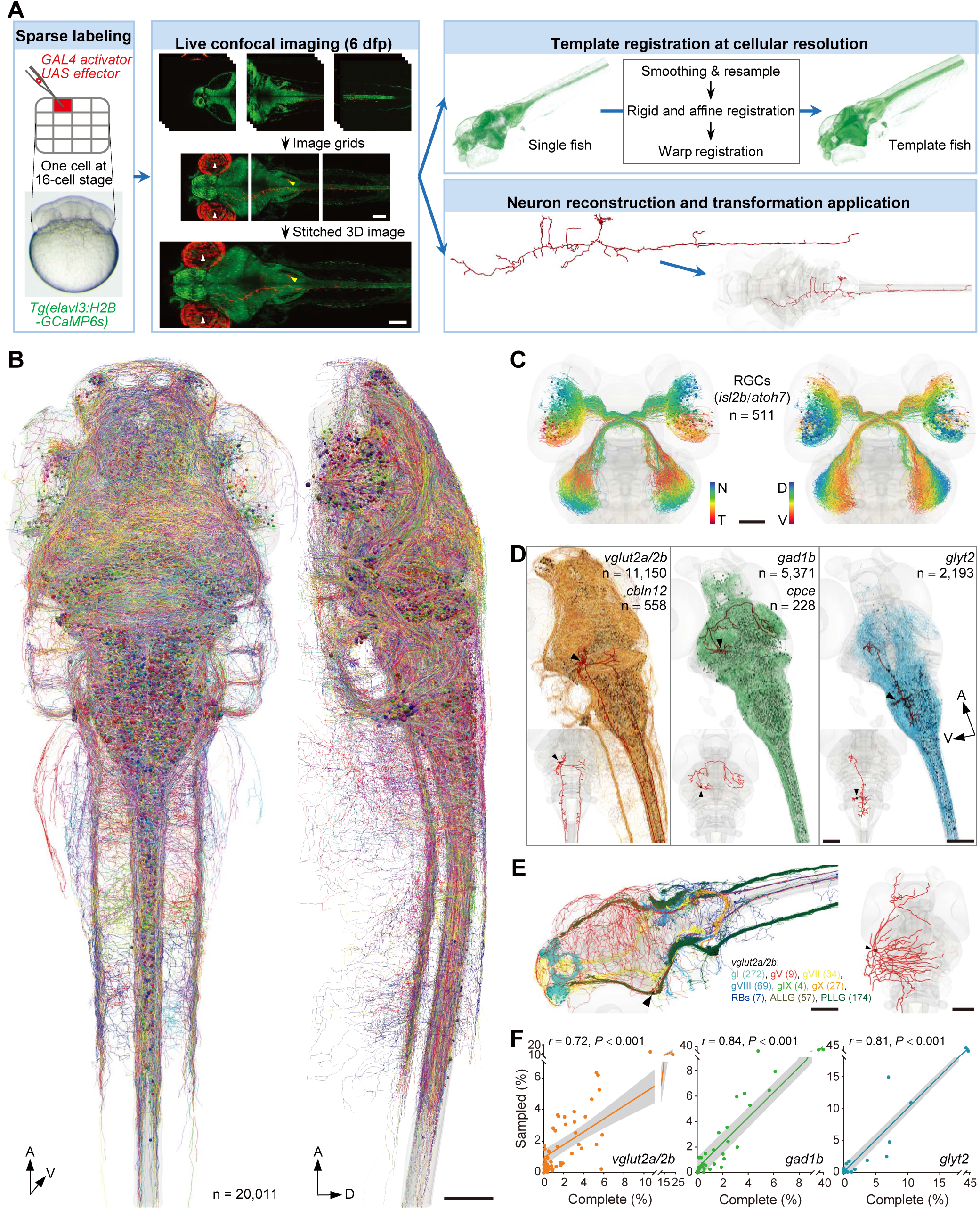
Nervous system-wide single-neuron morphology atlases of E and I neurons. (A) Pipeline of building single-neuron morphology atlases, including sparse labeling of neurons, 3D live imaging, neuron reconstruction, and alignment to the CPSv1. White arrowheads: autofluorescence of the eyeballs. (B) 3D visualization (left, dorsal view; right, lateral view) of 20,011 reconstructed E and I neurons from 13,429 fish, aligned to the CPSv1. Neurons are randomly color-coded. (C) RGCs (*isl2b*, n = 470; *atoh7*, n = 41) color-coded by soma position along the nasal-temporal (left) and dorsal-ventral (right) axes, displaying spatial topography. (D) Single-neuron atlases of excitatory (left: *vglut2a/2b*, n = 11,150; *cbln12*, n = 558), GABAergic (middle: *gad1b*, n = 5,371; *cpce*, n = 228), and glycinergic (right: *glyt2*, n = 2,193) inhibitory neurons. Example neuron tracings are shown in red in the main panels and in the insets, with black arrowheads indicating the soma. (E) Map of sensory neurons including color-coded cranial sensory (including gI, gV, gVII, gVIII, gIX, and gX), Rohon-Beard (RB), and lateral line ganglion neurons (including ALLG and PLLG). Right, example neuron tracing. All sensory neurons are marked by *vglut2a/2b*. The numbers in the parentheses indicate the neuron number. gI, gV, gVII, gVIII, gIX, gX: ganglion of the cranial nerve I, V, VII, VIII, IX, or X; ALLG, PLLG: anterior or posterior lateral line ganglion. (F) Correlation of relative regional distributions between sampled and complete neurons across 52 anatomical regions (sub-regions in the spinal cord and vagal ganglia were merged). Left, glutamatergic; Middle, GABAergic; Right, glycinergic. Each dot represents the data for a single anatomical region. Pearson correlation and linear fitting revealed a significant correlation. Sampled and complete neurons refer to the reconstructed neurons (D) and counted neurons in cytoarchitectures (see Figures 2A–2C), respectively. Scale bars: 50 μm (C), 100 μm (A, B, D, and E). n, the number of neurons shown. A, anterior; D, dorsal; N, nasal; T, temporal; V, ventral. See also Figure S3 and Video S3.

Using this standardized workflow involving sparse neuron labeling, live imaging, neuron reconstruction, and image registration, we built a single-neuron morphology atlas comprising 20,011 E and I neurons (n_E_ = 12,219, n_I_ = 7,792; excitatory: n*_vglut2a/2b_*= 11,150, n*_cbln12_* = 558, n*_isl2b/atoh7_* = 511; inhibitory: n*_gad1b_* = 5,371, n*_cpce_* = 228, n*_glyt2_* = 2,193; n_brain_ = 16,741, n_spinal cord_ = 1,600, n_peripheral ganglia_ = 1,159, n_retina_ = 511), distributed comparably between hemispheres (n_left_ = 9,962, n_right_ = 10,049) and covering the brain, rostral spinal cord, peripheral ganglia, and retina (Figures 3B–3E and Video S3). This dataset covers over 25% of the total E and I neurons counted (∼76,000; excluding the retina), and ∼24.5% of those in the brain (∼68,349) (see cytoarchitecture data above). For subsequent analyses, retinal (*isl2b*/*atoh7*) and granule (*cbln12*) excitatory neurons were grouped with *vglut2a/2b*, and Purkinje neurons (*cpce*) with *gad1b* to form unified E and I categories.

To assess the registration precision of the reconstructed morphology atlas, we first measured alignment accuracy in 88 fish using the landmark-based approach (see Methods),^44^ yielding a MLD of 3.28 ± 0.06 μm (Figures S3A–S3F), which is below the average neuronal soma diameter (5.80 ± 0.001 μm, measured from all reconstructed neurons). Because landmarks were identified at high-contrast anatomical boundaries that were easier to align, we further evaluated registration precision in homogeneous neuronal regions. We quantified the 3D soma-to-soma distances of individual Mauthner cells identified by spinal dye backfills^45^ (Figure S3G; see Methods) and single-neuron reconstructions (Figure S3H), yielding a mean distance of 6.06 ± 0.32 μm (Figure S3I). These results indicate that our registration achieves near-cellular resolution.

We next assessed sampling unbiasedness by comparing regional distributions of reconstructed neurons with those in the cytoarchitecture atlas. All E and I neuronal populations showed strong correlations (Figure 3F and S3J–S3O; *vglut2a/2b*: *r* = 0.72; *gad1b*: *r* = 0.84; *glyt2*: *r* = 0.81; all *P* < 0.001), suggesting unbiased sampling. To further test robustness, we performed repeated subsampling at varying sizes. Even with only a few hundred neurons, the resulting correlations remained comparable to those obtained from larger sample sizes (Figure S3P), further supporting the highly stochastic and unbiased sampling yielded by transient transgenic labeling.

Taken together, we established a single-neuron morphology atlas annotated with E and I neurotransmitter identities, spanning the retina, brain, spinal cord, and peripheral ganglia. In the atlas, the body-wide template allows the alignment of neurites from cranial sensory and motor neurons that extend beyond the brain and spinal cord. Therefore, this comprehensive atlas enables the mining of neural circuits from sensory inputs to motor outputs at a whole-body scale.

### Distinct morphological and projection features of E and I neurons

We next characterized the morphology of E and I neuronal populations. From *vglut2a/2b* to *gad1b* to *glyt2* neurons, the number of branch points decreased, while the maximum branch order increased slightly, implying different branching patterns among these three types of neurons (Figures 4A–4C). The *vglut2a/2b* neurons were larger in size than both *gad1b* and *glyt2* neuronal populations, as revealed by the total cable length and maximum Euclidian distance from the soma (Figures 4A, 4D, and 4E). Among I populations, *glyt2* neurons were larger than *gad1b* neurons (Figures 4D and 4E), consistent with their role in connecting hindbrain and spinal cord structures to upper brain regions (see Figure 3D, typical case). Within each E and I neuronal population, the total length positively correlated with the number of branch points, suggesting an interrelation between neuronal size and structural complexity (Figure 4F). In accordance, the larger the size of the neurites (*vglut2a/2b* > *glyt2* > *gad1b*), the more regions they innervated (Figure 4G).

**Figure 4.**
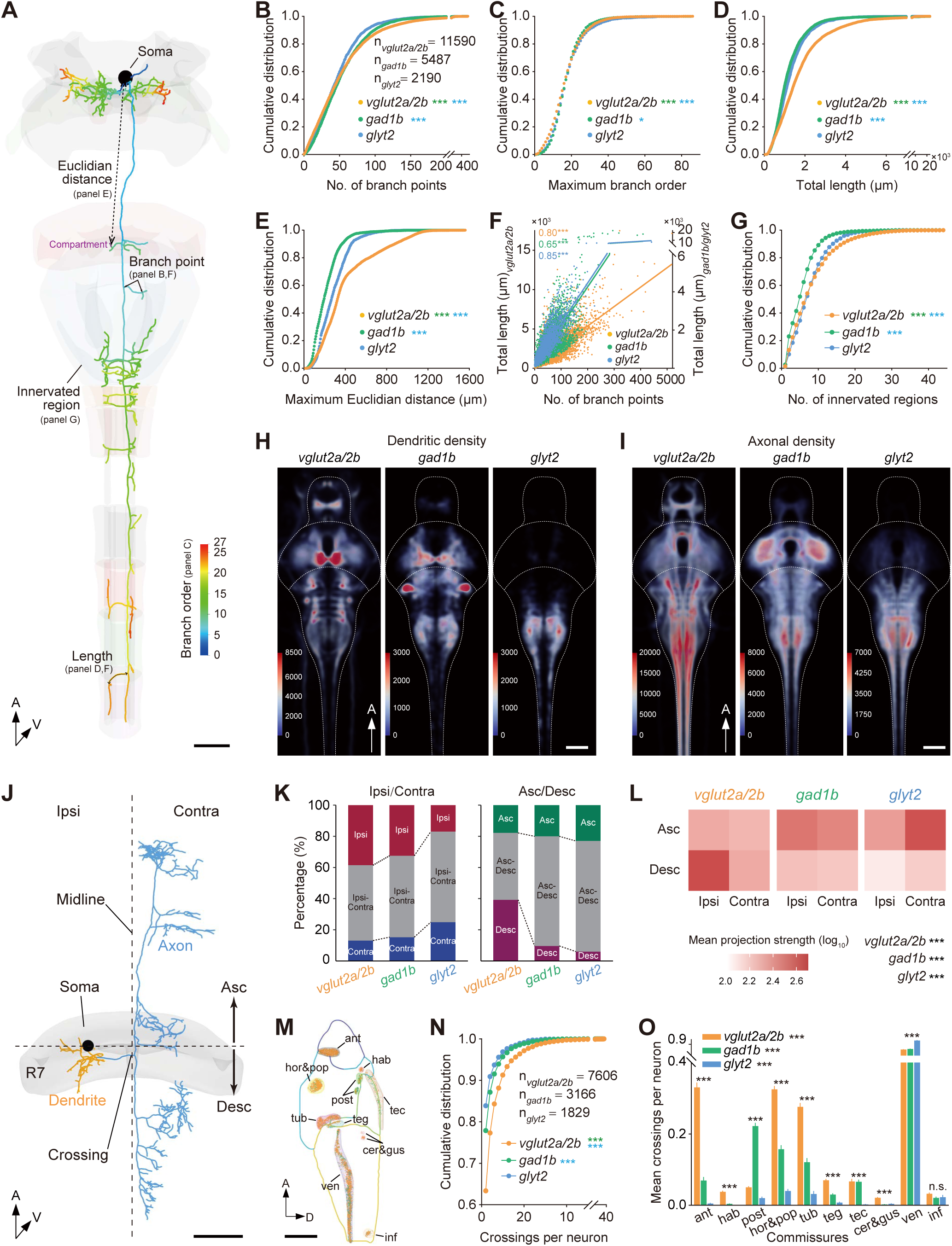
Distinct morphological features, dendrite-axon distribution, and axonal projection patterns of E and I neurons. (A) Volume rendering of a single reconstructed *vglut2a*+ neuron located in the nucleus of the medial longitudinal fasciculus (nMLF) and its target brain regions. Neurites are color-coded by branch order. The meanings of morphological measurements shown in (B–F) are indicated, where the Euclidian distance represents the distance between the soma and the closer endpoint of each branch segment relative to the soma. (B–E) Cumulative distribution of four morphological parameters in E (*vglut2a/2b*) and I (*gad1b* and *glyt2*) neurons, showing differences in structural complexity (B and C) and size (D and E). Color asterisks represent corresponding inter-cell population comparison (Kolmogorov-Smirnov test). Mean number of branch points in (B): *vglut2a/2b* (54.2 ± 0.4) > *gad1b* (53.2 ± 0.5) > *glyt2* (48.2 ± 0.7); Mean maximum branch order in (C): *glyt2* (18.5 ± 0.2) > *gad1b* (17.9 ± 0.1) > *vglut2a/2b* (17.4 ± 0.1); Mean total length in (D): *vglut2a/2b* (1764 ± 13 μm) > *glyt2* (1296 ± 18 μm) > *gad1b* (1236 ± 15 μm); Mean maximum Euclidian distance in (E): *vglut2a/2b* (410.8 ± 2.6 μm) > *glyt2* (289.6 ± 3.1 μm) > *gad1b* (218.2 ± 1.7 μm). Neurons with truncated axons innervating the spinal cord were excluded. (F) Scatterplot showing the correlation between the number of branch points and the total branch length for the three neuronal populations. Correlation was assessed using the Pearson correlation and linear fitting (*r_vglut2a/2b_* = 0.80, *r_gad1b_*= 0.65, *r_glyt2_* = 0.85). (G) Cumulative distribution of the number of innervated regions. Pairwise comparison among the three neuronal populations was performed using Kolmogorov-Smirnov test, and *P* values are represented by color asterisks. (H and I) Volume rendering of the density maps of dendritic (H) and axonal (I) branches. Colors indicate point density within a 5 μm-radius sphere. The maximum number of calibration bars indicates the density value at which the heat map saturates. (J) Volume rendering of a single reconstructed *vglut2a* cranial relay neuron with the soma located in the R7,^51^ illustrating the definition of ipsilateral (“Ipsi”)/contralateral (“Contra”) and ascending (“Asc”)/descending (“Desc”) axons (light blue). (K) Proportions of neurons with Ipsi, Contra, or bilateral (Ipsi-Contra) axonal projections (left), or with Asc, Desc, or bidirectional (Asc-Desc) axonal projections (right). (L) Heatmap of mean projection strengths across four axonal trajectory quadrants. Asterisks represent comparison among the four quadrants (nonparametric Friedman test with Dunn’s post hoc multiple comparisons). (M) Sagittal view of 10 commissural regions with midline-crossing points of contralateral and ipsilateral-contralateral projections. (N) Cumulative distributions of the number of crossings per neuron. Color asterisks represent corresponding inter-cell population comparison (Kolmogorov-Smirnov test). (O) Distribution of the mean number of crossings through different commissures at single cell level. Asterisks indicate inter-commissure difference within each cell population (shown beside of the name of neuronal populations) and inter-cell-population differences within each commissure (shown above bars) (Two-way ANOVA with Tukey’s post hoc multiple comparison test). Scale bars: 100 μm (A, H–J, and M). The n represents the number of neurons examined. A, anterior; D, dorsal; V, ventral. Spatially adjacent commissures were treated as a single commissural region for quantification, including the hor and pop, and the cer and gus. ant, anterior commissure; cer&gus, cerebellar and secondary gustatory commissures; hab, habenular commissure; hor&pop, horizontal and postoptic commissures; inf, commissure infima of Haller; post, posterior commissure; tec, intertectal commissure; teg, ventral tegmental commissure; tub, posterior tuberculum commissure; ven, ventral rhombencephalic commissure. n.s., not significant; \**P* < 0.05; \*\*\**P* < 0.001. See also Figure S4.

To analyze axonal innervation patterns and define input-output domains of neurites, we annotated polarized dendritic and axonal compartments of reconstructed neurons. We generated a bidirectional UAS reporter plasmid *Synaptophysin (Sypb)-EGFP-tdKatushka2-CAAX* to simultaneously label the whole morphology and presynaptic sites of individual E and I neurons and identified axonal compartments based on Sypb localization (Figures S4A-S4F).We found three recurring structural configurations that distinguish dendritic and axonal compartments in projection neurons: (1) dendrites and axons emerge from the soma as separate primary branches (bipolar or multipolar); (2) a single primary fiber bifurcates near the soma into dendritic and axonal arbors (pseudounipolar); or (3) a single fiber undergoes multiple proximal bifurcations before producing a relatively long daughter branch that forms the axonal stem. Leveraging these features, we developed a machine-learning algorithm to automatically classify the dendrite-axon compartments (see Methods). We evaluated its performance using 12 morphotypes with known polarity (Figures S4G–S4I; see below in Figures 5, S5, and S6), achieving a 97% ± 0.01% accuracy. To ensure reliability, we excluded most local neurons from polarity annotation and manually reviewed dendrite-axon assignments across annotated morphotypes (see Figure S5). In total, we validated and annotated 18,084 neurons with dendrite-axon polarity.

**Figure 5.**
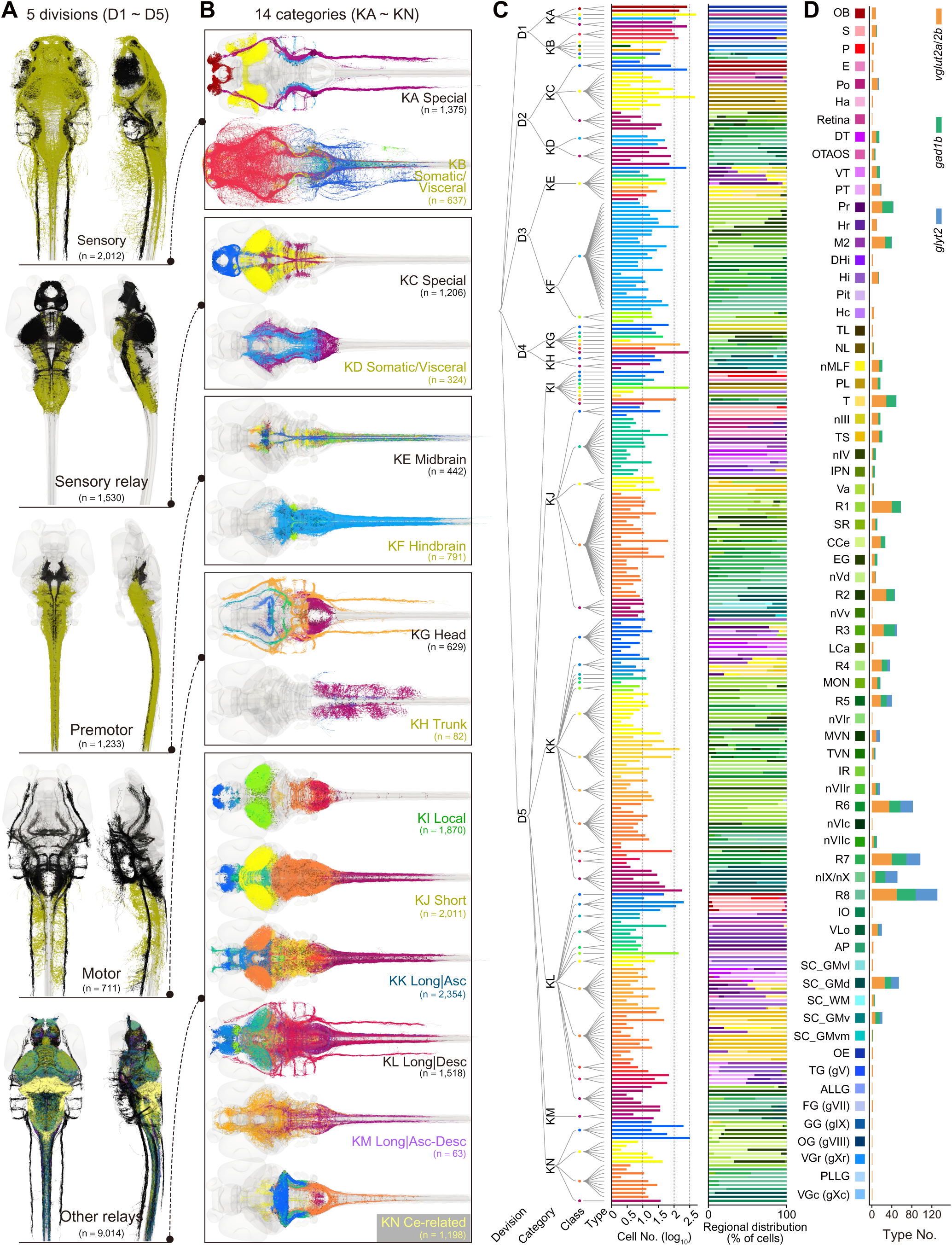
Morphological classification of E and I neurons. (A and B) Volume rendering of all E and I morphotypes grouped into 5 divisions (A, dorsal and lateral views) and 14 categories (B, KA to KM, dorsal view). Neuronal tracing colors in (A) match the color of category names in (B), and neuronal tracing colors in (B) match the color of class symbols in (C). Cells in the midbrain category are color-coded by type, as their class and category are the same (C) Dendrogram of E neurons comprising 5 divisions, 14 categories, 71 classes, and 305 types, with bar plots showing cell number and regional distribution. The color codes for brain regions are shown in (D). (D) Bar plot showing the number of E and I morphotypes per region, ordered by the anteroposterior centroid within primary brain subdivisions, spinal cord, and peripheral ganglia. The n represents the number of cells. See also Figures S5 and S6, and Table S2.

With this dendrite-axon annotated single-neuron-morphology dataset, we first obtained a global view of dendrite-axon (input-output) distribution patterns across the E and I neuronal populations. We extracted all node points along dendritic and axonal branches and transformed the points into density map (Figures 4H and 4I). The results showed distinct distributions of dendrites and axons among E and I neuronal populations, while also highlighting shared areas of high density, including the tectal neuropil, nMLF, ventral anterior hindbrain, lateral middle hindbrain, and caudal hindbrain. These input-output hotspots are linked to the sensorimotor integration and transformation areas in zebrafish,^46–50^ and are consistent with the hub regions revealed by the graph theoretical analysis of the interregional connectivity networks (see below in Figure 8).

We next investigated the projection features of E and I neurons at the single-cell level by analyzing their projection strength to specific brain regions. Here, we defined the projection strength of each neuron to a target region as the total length of coherent axonal arbors with terminals in that region (see Methods). We verified that this parameter is comparable to the number of presynaptic sites for evaluating projection strength by using reconstructed neurons with presynaptic site labeling (Figures S4J and 4K; Pearson’s *r* = 0.784, *P* < 0.001), suggesting that the length of terminal-bearing axonal fibers is an alternative parameter for assessing projection strength.

As the architectures of ipsilateral/contralateral and ascending/descending axonal projections (Figure 4J) shape interhemispheric and top-down/bottom-up information flow, we qualitatively categorized the neurons into three groups according to ipsilateral/contralateral (Figure 4K, left) or ascending/descending projections (Figure 4K, right; see Methods). Across all the three E and I neuronal populations, 53% of neurons exhibited bilateral (ipsilateral-contralateral) projections, and 61% exhibited bidirectional (ascending-descending) projections. Notably, from *vglut2a/2b* to *gad1b* to *glyt2* neurons, the percentage of contralateral-only projecting neurons increased, while the percentage of descending-only projecting neurons decreased. Quantitatively, we calculated the axonal projection strengths across four quadrants: ascending-ipsilateral, ascending-contralateral, descending-ipsilateral, and descending-contralateral. The *vglut2a/2b*, *gad1b*, and *glyt2* neurons displayed significantly stronger projections in the ipsilateral-descending, ipsilateral-ascending, and contralateral-ascending directions, respectively (Figure 4L), revealing distinct projection biases across E and I neuronal populations.

To further understand how contralateral communication is implemented anatomically, we examined how often axons of contralateral-projecting neurons cross the midline and which major brain commissures they travers (Figure 4M), noting that midline-crossing dendrites are rare. On average, ∼75% of contralateral and ipsilateral-contralateral projecting neurons from all three E and I populations crossed hemispheres only once (Figure 4N), with around 69% of crossings occurring within 20□μm of the soma along the anteroposterior axis. Multiple crossings existed, with up to dozens in some neurons, contributing to a significant discrepancy in the total number of crossings between the E and I neuronal populations (Figure 4N). These multiple crossings arose either from the main axonal path traversing the midline repeatedly via different commissures and/or from collateral branches extending contralaterally. We further analyzed the distribution of crossings through ten different commissural regions for all three E and I neuronal populations at the single cell level and found that the ventral rhombencephalic commissure (“ven”) is the most frequently used by all three populations (Figure 4O). In terms of differences among the three neuronal populations, the number of crossings was higher for E neurons compared with I neurons for most commissural regions, especially the anterior (“ant”), horizontal & postoptic (“hor&pop”), and posterior tuberculum (“tub”) commissures, while the *gad1b* neurons showed a greater number of crossings in the posterior commissure (“post”) (Figure 4O).

Taken together, E neurons exhibit greater structural complexity, larger size, and broader innervation patterns compared to I neurons. Furthermore, E neurons display preferential ipsilateral-descending projections, whereas I neurons show contralateral-ascending projections. These biases in projection laterality and directionality, together with distinct interhemispheric crossing patterns, provide a comprehensive overview of the distinct projection frameworks of E and I neuronal populations throughout the brain.

### Morphological classification of E and I neurons

To analyze the morphotypes of E and I neurons, we developed a methodology termed MBLAST (named by referring to NBLAST^52^) to assign cell types to large-scale neuron reconstruction datasets based on morphological similarity (see Methods). Based on MBLAST, we classified *vglut2a/2b*, *gad1b*, and *glyt2* neurons separately into morphotypes (Figure S5; 14,301 cells in total, including n*_vglut2a/2b_* = 9,774, n*_gad1b_*= 3,411, n*_glyt2_* = 1,116), reflecting differences in the structural morphology of dendrites and axons.

To organize all neuronal morphotypes within a framework reflecting macroscale functional and structural organization, we established a five-level nested hierarchy, comprising divisions (D), categories (K), classes, types, and subtypes. At the two highest levels (divisions and categories), both excitatory (*vglut2a/2b*) and inhibitory (*gad1b*, *glyt2*) neurons were assigned to the same organizational scheme, including 5 divisions (D1 to D5) and 12 categories (KA to KM) (Figures 5A and 5B). The highest-level division was defined based on potential roles of neurons in sensorimotor information processing, containing the divisions of sensory, sensory relay, premotor, motor, and other relays (Figure 5A). The first four divisions each contain two categories (Figure 5B). The sensory and sensory relay divisions were categorized according to the modality of sensations, distinguishing between special (including olfaction, vision, audition, and vestibular) and somatic/visceral components.

The premotor and motor divisions were categorized based on the location of cells—the premotor division was divided into midbrain and hindbrain categories, while the motor division was divided into head and trunk categories. The fifth division, ‘other relays’, contained all remaining morphotypes that lie between the sensory and motor parts (Figure 5B). It was divided into six categories based mainly on the range and direction of axonal projections: local (projections within one specific brain region), short (short-range projections across adjacent brain regions within the same primary subdivision), long|asc (long-range ascending projections across different primary subdivisions), long|desc (long-range descending projections across different primary subdivisions), long|asc-desc (long-range bidirectional projections across different primary subdivisions), and cerebellum-related (morphotypes in the cerebellum and those of cerebellar efferents and afferents).

At the level of class, neurons were organized separately within each neurotransmitter population (glutamatergic, GABAergic, or glycinergic), with types and subtypes forming parallel hierarchies for distinct E and I neuronal populations (Figures 5C, S6A, and S6B). The classification of neurons was also largely determined by their potential functional roles in information processing. For instance, within the special categories in sensory and sensory relay divisions, the classes included specialized morphotypes involved in sensing or relaying information for smell, vision, hearing, and vestibular. Reticulospinal neurons (RSNs) were assigned as a class in the hindbrain category in the premotor division. Different motor branches of cranial nerves belong to different classes in the head category of the motor division. Classes in the ‘other relay’ division were mainly defined based on their major distribution and projection regions.

In total, we classified 71 classes, 305 types, and 348 subtypes for E neurons (*vglut2a/2b*), 36 classes, 155 types, and 164 subtypes for GABAergic I neurons (*gad1b*), and 19 classes, 84 types, and 88 subtypes for glycinergic I neurons (*glyt2*). The full morphotypes at the level of subtypes were displayed in Figure S5, and each subtype was assigned with an ID (up to 11 characters) encoding its neurotransmitter identity (Glu, GABA, or Gly), division (one-digit number), category (A–N), class (two-digit numbers), type (two-digit numbers), and subtype (a lowercase letter). To assess cross-dataset compatibility, we further incorporated previously published neuronal reconstructions from the mapzebrain atlas, in which pan-neuronal markers were primarily used to label individual neurons.^36^ The majority of these reconstructed neurons (1,326 out of 2,098 neurons) were matched to morphotypes in our inventory, corresponding to 122 *vglut2a/2b* types, 31 *gad1b* types, and 9 *glyt2* types. The remaining unmatched neurons likely represent morphotypes that are underrepresented or belong to neurotransmitter types beyond the E and I populations, highlighting future expansion of the taxonomy as additional data become available.

To further evaluate the sampling coverage of our datasets (see Figure S3P), we assessed cell type distributions within the sampled population. We repeatedly (1,000 rounds) selected subsets of cells (incrementing by 10 cells per step) from the classified cell pool and compared the cell type composition between the subset and the overall population. The cell type composition began to show no significant dissimilarity when sampling reached ∼17% (on average) of the corresponding E and I classified neuronal populations (Figures S6C–S6E, start point of the black-only region of the gray/black dot trace), though the overall cell type coverage remained below 50%. To determine the minimal sampling size after which cell type coverage stabilizes, we applied a structural breakpoint detection method^53,54^ to the cell type coverage curve and found that coverage stabilized at ∼22% (on average) of the corresponding neuronal population (Figure S6C–S6E, vertical dotted color lines), resulting in an average cell-type coverage rate of nearly 65%.

Next, we characterized the distribution of neurons of each type across regions (right in Figures 5C, S6A, and S6B) and examined the type diversity within individual regions (Figure 5D). This bidirectional relationship between morphotypes and brain regions provides information on the regionalization of cell types and the complexity of connections among different regions. The constructed graphs revealed two prominent features: (1) with the exception of some types in the sensory and motor divisions, most types are not constrained to a single region; however, they are typically distributed within one primary subdivision, as indicated by their shared color tone representing the specific subdivision (right in Figures 5C, S6A, and S6B); (2) the hindbrain shows a higher degree in both the number of regions occupied by a single type (2.3 ± 1.4) and the number of E and I morphotypes within a single region (29.2 ± 6.9). Although these analyses may be sensitive to the granularity of cell clustering, these results indicate that neurons in different primary subdivisions have distinct projection patterns. In particular, the findings suggest that the hindbrain contains more computing elements for sending premotor output to the spinal cord and efference copy/corollary discharge to upper brain regions.^55^

Overall, based on large-scale single-neuron morphological reconstructions, we established a system-wide taxonomy of E and I neurons across the zebrafish nervous system. By linking morphotypes with potential functional roles and regional organization, this framework provides a structural basis for identifying neuronal types and dissecting circuit architecture.

### Cabling design of E and I morphotypes

To examine the innervation characteristics of the E and I morphotypes, we constructed type-to-region innervation matrices (Figures S7A–S7C; see Methods). The CPSv1 contains 68 nervous system-wide anatomical regions at the lowest level of the ontology tree (see Figures 1C–1E and Table S2). We generated the matrices separately for dendrites and axons and distinguished ipsilateral and contralateral projections across the 68 regions.

Within individual classes, different neuronal types often diverged in either their dendritic arborization or axonal projection directionality (Figures 6A–6E). For example, two types in the nMLF possessed comparable axonal morphologies characterized by multiple collaterals innervating the contralateral spinal cord, yet their dendritic configurations showed partial overlap alongside marked differences (Figures 6A–6C; Glu-3E0104 *vs*. Glu-3E0107), indicating divergent sources of dendritic input. Conversely, mitral cells in the olfactory bulb displayed similar dendritic arborizations, but could be classified into two types based on distinct axonal projection patterns (Figures 6A, 6D, and 6E; Glu-2C0102 *vs*. Glu-2C0103). These results indicate that the diversification of neuronal types can arise from variation in either dendritic architecture or axonal organization.

**Figure 6.**
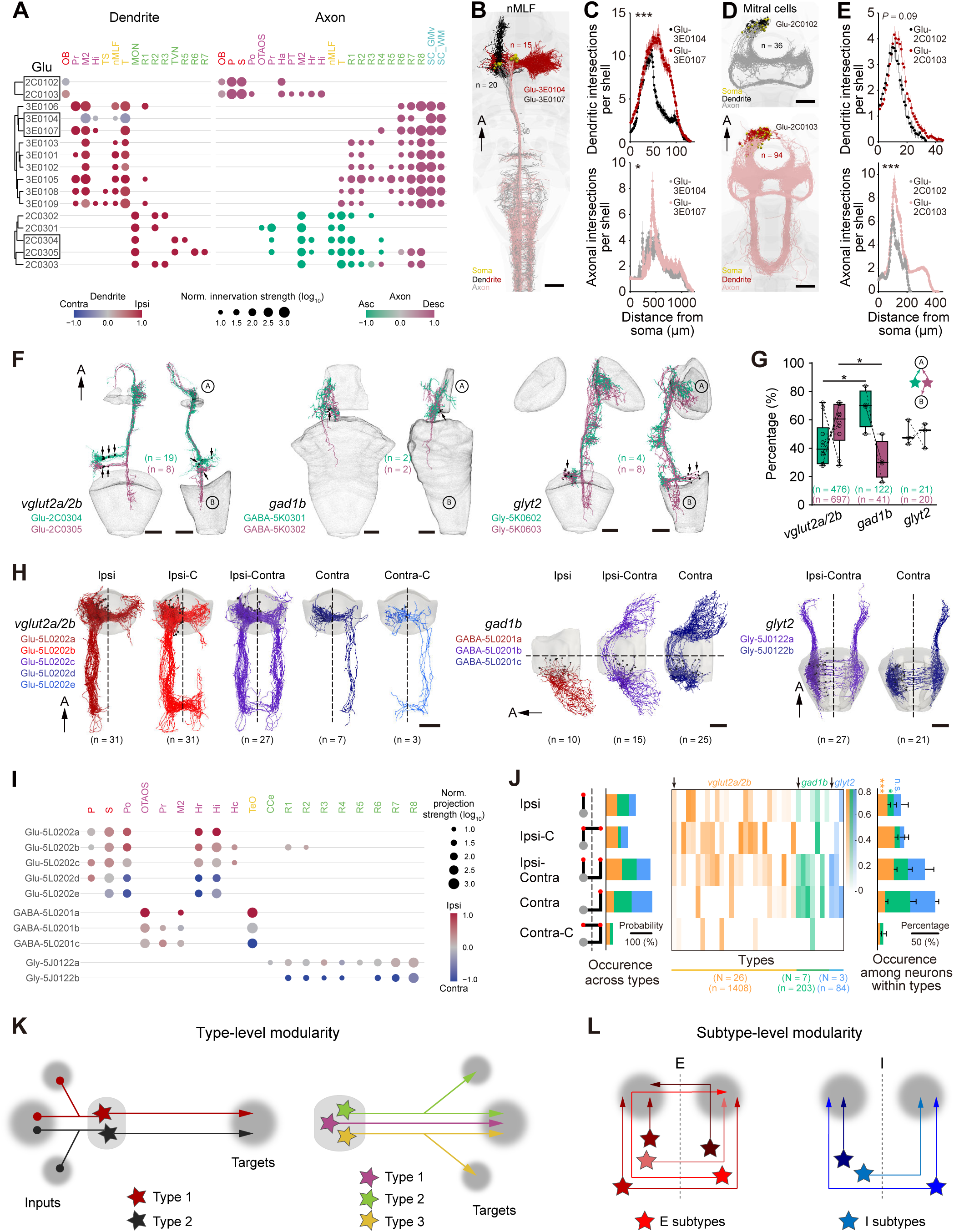
Projection design of E and I morphotypes. (A) Dot plot showing dendritic (left) and axonal (right) innervation strength (IS, dot size) and preference (dot color) for ipsilateral/contralateral (dendrite, (IS_ipsi_−IS_contra_)/(IS_ipsi_+IS_contra_)) or descending/ascending (axon, (IS_desc_−IS_asc_)/(IS_desc_+IS_asc_)) innervations of neuronal types from three classes across regions. Black boxes indicate example types shown in (B, D, and F). (B and D) Examples of two *vglut2a/2b* neuronal types within one class showing similar axonal but different dendritic innervations (B, MeL (black) *vs*. MeM (red) neurons in nMLF^56^), or similar dendritic but different axonal innervations (D, mitral cells in the OB). (C and E) 3D Sholl analysis (branch intersections with concentric spheres centered on the soma) of dendrites (top) and axons (bottom) from types shown in (B and D). The radius increment is 1 μm. The two tailed Mann-Whitney test was used. (F) Volume rendering of two types (seagreen and marron) within one class in each E and I neuronal population, featured by their common ascending projection pathways. Somata (arrows) in black. A and B represent two innervated regions. (G) Boxplot showing the proportions of ascending-only (seagreen) and ascending-descending (marron) projection types (two types sum to 100%) across the three E and I neuronal populations. Inset: schematic illustrating two types sharing an ascending projection trajectory. The central mark, bottom, and top of the boxplot indicate the median, first, and third quartiles, respectively. Two-way ANOVA with Tukey’s post hoc multiple comparison test was performed. (H) Examples showing ipsilateral/contralateral projection subtypes within one type with targeted bilateral projections in *vglut2a/2b* (left), *gad1b* (middle), and *glyt2* (right) neurons. Dendrites in darker color. Midline indicated by a dotted line. (I) Dot plot showing projection strength (PS, dot size) and preference (dot dolor) for ipsilateral/contralateral projections ((PS_ipsi_−PS_contra_)/(PS_ipsi_+PS_contra_)) of subtypes in (H). (J) Heatmap (middle) of cell proportions of five ipsilateral/contralateral projection modes (subtypes) in each corresponding type (column, N = type count; see Figure S5) from E and I neuron populations, and summary of mode occurrence probability across types (left) and among individual neurons within types (right). Arrows indicate the examples shown in (H). Asterisks represent intra-neuronal population comparisons (nonparametric Friedman test with Dunn’s post hoc multiple comparisons). (K and L) Schematic illustrating the type-level (K) and subtype-level (L) modularity underlying hierarchical classification from class to type (K) and from type to subtype (L), respectively. Scale bars: 50 μm (B, D, F, and H). Regions in (A and I) are ordered by their centroid positions along the anteroposterior axis within primary subdivisions indicated by different colors. IDs of corresponding morphotypes are shown. Numbers in parentheses indicate the number of neurons shown or examined. A, anterior. Ipsi, ipsilateral; Ipsi-C, ipsilateral-collateral; Ipsi-Contra, ipsilateral-contralateral; Contra, contralateral; Contra-C, contralateral-collateral. n.s., not significant; \**P* < 0.05; \*\*\**P* < 0.001. See also Figures S7A–S7C, and Table S2.

Similar to mitral cells, we found many neuronal types within a class shared a conserved primary projection trajectory but diverged through additional paths targeting distinct regions. This organization was frequently represented by two types sharing an ascending trajectory, with one type additionally extending a descending projection (Figure 6F). For instance, two types (Glu-2C0304 and Glu-2C0305) in the class of vestibular relay neurons^57^ (see Figure 5B, KC, marron, *vglut2a/2b*) in rhombomeres 5/6 projected to cranial motor neurons in nIII/nIV through a shared ascending trajectory, while one type (Glu-2C0305) also extended a descending path to rhombomeres 7/8 (i.e., the caudal hindbrain; Figure 6F, left). The relative proportions of these ascending-only and ascending-descending types varied among the three E and I neuronal populations (Figure 6G), consistent with the stronger descending projection bias observed in E neurons compared with I neurons (see Figure 4L).

Within individual types, different neuronal subtypes differed primarily in projection laterality. For a single type targeting mirror-symmetric regions in both hemispheres, bilateral innervation is not solely mediated by a single ipsilateral-contralateral mode but instead achieved through a combination of distinct projection configurations, including ipsilateral (Ipsi), ipsilateral-collateral (Ipsi-C), ipsilateral-contralateral (Ipsi-Contra), contralateral (Contra), and contralateral-collateral (Contra-C) (Figures 6H and 6I). These configurations comprised main axonal paths with or without midline-crossing collaterals and were observed in both descending and ascending projections. Neurons adopting different laterality patterns were classified as subtypes. Quantitative analysis revealed marked differences between E and I neurons in the utilization of these configurations: *vglut2a/2b* E neurons preferentially utilized ipsilateral-related modes, whereas *gad1b* and *glyt2* I neurons predominantly relied on ipsilateral-contralateral and contralateral projections, rarely employing collaterals (Figure 6J). These findings indicate that E and I neurons employ distinct strategies to achieve bilateral innervation.

Taken together, these analyses reveal two fundamental organizational principles governing projection architecture. First, type-level modularity arises when related types share a conserved core projection trajectory but diverge in either dendritic architecture or axonal projection directionality, enabling the diversification of input integration or the coordinated innervation of multiple brain regions within individual classes (Figure 6K). Second, subtype-level modularity emerges through variations in projection laterality, yielding ipsilateral, contralateral, or bilateral subtypes that collectively ensure bilateral coverage of target regions by individual types (Figure 6L).

### Putative inter-cell-type connectome and neural circuit mining

To investigate the nervous system-wide connectivity of E and I neurons, we constructed a neural network based on type-level morphotypes with dendrite-axon annotations. GABAergic and glycinergic neurons with similar morphologies were grouped into a single type (see Figure S5B), resulting in 197 I types. Together with 305 E types, this yielded a total of 502 types.

Spatial proximity between dendrites and axons serves as a reliable proxy for synaptic contacts and can be used to infer putative synaptic connectivity between neuron pairs.^58–62^ We therefore constructed inter-cell-type putative connectivity matrices by quantifying spatial proximity between axonal and dendritic branch segments, measured as the Euclidean distance between segment centers, across the 502 E and I types (see Methods). Correlation analysis of these matrices across different distance thresholds revealed that pairwise similarity peaked at 2 µm (within a 0.5–4 µm range; Figures S7D–S7F), implying that this threshold provides a stable and representative estimate of the underlying connectivity. Considering that smaller thresholds (0.5 and 1 µm) may yield false negatives while larger ones (3 and 4 µm) risk false positives, we selected 2 µm as the optimal cutoff to generate a directed, weighted inter-cell-type putative mesoscopic connectome (Figure S7G).

We characterized this type-level connectivity using three parameters: (1) neuronal participation, defined as the fraction of neurons within a type that connect to a partner type. Across 32,803 type pairs, over 60% of types showed ≥50% participation, indicating robust inter-cell-type connectivity (Figure 7A); (2) divergence/convergence (D&C) index, reflecting how broadly neurons of a given type distribute outputs or integrate inputs across partner types. This index followed a single-phase decay distribution (R² = 0.98), suggesting a structured organization of D&C within the connectome (Figure 7B); and (3) connection strength, representing the number of putative contacts between types. This parameter followed a Gaussian distribution (R^2^ = 0.95) with a mean of 16.62 ± 0.04, reflecting the typical range of putative connectivity across the network (Figure 7C).

**Figure 7.**
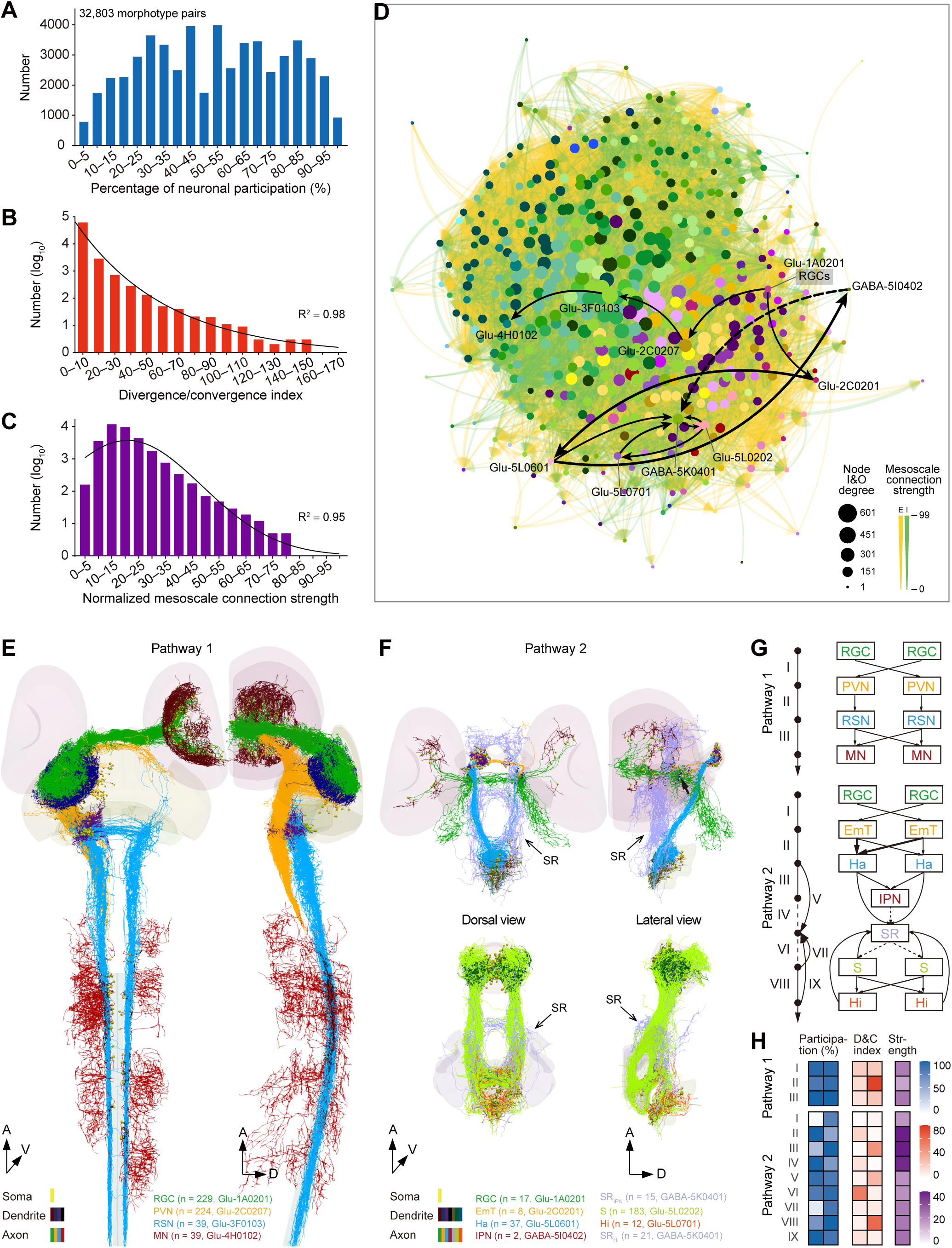
Directed and weighted inter-cell-type putative connectome supporting neural circuit mining. (A–C) Distributions of the percentage of neuronal participation (A), divergence/convergence index (B), and normalized mesoscale connection strength (C), which are calculated based on 32,803 connected morphotype pairs within 475 morphotypes (see Figure S7D). For each connected morphotype pair, two values were calculated from the input and output morphotypes for the metrics in (A) and (B). Distributions of (B) and (C) were fitted respectively with a one-phase decay (R^2^ = 0.98) and Gaussian (R^2^ = 0.95), with the *y*-axis plotted in log scale. (D) Graph visualization (the NetworkX package was used) of the inter-cell-type E and I network with 475 nodes and 32,803 edges. Node size represents the total degree (total number of input and output connections), and node color indicates its primary distributed region, following the coloring scheme shown in Figure 5D. The thickness of edges with arrows represents normalized directed connection strength, color-coded by E (orange) or I (green) attribute. Black edges highlight two visual pathways shown in (E and F). (E and F) 3D visualization of two visual pathways starting from RGCs, rendered by morphologies of participating E and I types. Dendrites and axons are color-coded differently, and only connected neurons are shown, with distributed regions serving as the background. In (E), only the right retina-involved pathway is shown. In (F), the pathway is split into two parts (top and bottom), linked via a morphotype in the superior raphe (SR, open arrow). Black arrow (F) points to the arborization fields 3 and 4 (AF3 and AF4). Involved morphotypes with their cell numbers and IDs are annotated at the bottom. (G) Wiring diagrams of the two pathways in E and F, with each morphotype node named by attributed neuronal class or distributed brain region. The two-hemisphere layout displays ipsilateral, contralateral, or bilateral projections. The coloring scheme matches the axon colors shown in (E and F), and solid and dashed edges indicate E and I connections, respectively. Notably, EmT-to-Ha connections show left-side preference. (H) Connection metrics for the two pathways in (E and F), including the percentage of neuronal participation (“Participation (%)”) for input (left) and output (right) types, divergence/convergence index (“D&C index”) for input (left) and output (right) types, and normalized connection strength (“Strength”) between input and output types. Scale bars: 50 μm (E and F). A, anterior; D, dorsal; V, ventral. EmT, eminentia thalami; Ha, habenula; Hi, intermediate hypothalamus; IPN, interpeduncular nucleus; MN, motor neuron; PVN, periventricular neuron; RGC, retinal ganglion cell; RSN, reticulospinal neuron; S, subpallium; SR, superior raphe. See also Figure S7D and Video S4.

We then illustrated the utility of the type-level connectome for circuit mining by identifying two visual pathways exhibiting relatively strong and weak connectivity to RGCs, respectively (Figures 7D–7H and Video S4). Pathway 1, characterized by strong connections with RGCs, represents a classical visuomotor circuit (Figures 7E, 7G, and Video S4). After receiving RGC innervation, it begins with tectal projection neurons (i.e., periventricular neurons, PVNs) in the optic tectum (TeO), a central hub for visual information processing. Among the connected PVN types, the most numerous are tectobulbar neurons with ipsilaterally and laterally positioned projections (iTB-L), which are involved in approach movements of zebrafish.^63^ From the TeO, the pathway continues to hindbrain RSNs, which act as premotor neurons before reaching the motor neurons (MNs) in the spinal cord. Analysis revealed that most of the 28 RSN types (see Figure S5, Glu-3F0101–Glu-3F0128) form connections with iTB-L neurons. We highlighted here a RSN type located in the rhombomere 1 (R1) with bilateral projections. It exhibits strong connectivity to motor neurons, marked by the highest convergence index (22.3) in comparison with other RSNs (Figures 7E and 7H).

Pathway 2 (Figures 7F, 7G, and Video S4), characterized by relatively weak connections with RGCs, aligns with and corroborates previous research on light-preference behavior in zebrafish.^64,65^ After receiving RGC innervation, this pathway begins with eminentia thalami (EmT) neurons, whose dendrites form connections with RGCs at the arborization fields 3 and 4 (AF3 and AF4) (Figure 7F, arrow in top-right image). The single-cell morphology of EmT neurons supports the preferred projection from bilateral EmT to the left dorsal habenula (Ha), consistent with previous functional and dye-tracing studies^64^ (Figures 7F and 7G). Accordingly, the downstream neurons in the habenula are mainly distributed on the left dorsal side. These neurons connect either directly to the primary inhibitory neurons in the superior raphe (SR) or indirectly to the SR inhibitory neurons via local inhibitory neurons in the IPN. The feedforward inhibition motif, involving Ha, IPN, and SR, may underlie the bimodal role of raphe serotonergic neurons in light-preference behavior.^65^ Notably, the strongest downstream connection from the superior raphe leads to neurons in the subpallium (S), a telencephalon region previously implicated in light/dark choice behavior in adult fish.^66^ In addition, we identified a feedback loop from the subpallium to superior raphe, either directly or indirectly via neuronal types in the intermediate hypothalamus (Hi), further highlighting the dynamic regulatory processes within these circuits (Figures 7F and 7G).

Taken together, we established a putative inter-cell-type mesoscopic connectome and demonstrated its utility in unraveling specific neural circuits, thereby providing a framework to guide future functional investigations.

### E and I neuron-based interregional connectome and information flow framework

While the putative inter-cell-type connectome provides intricate details about connectivity between neuronal types, it lacks information about brain regions. As brain regions serve as integral units for processing and transmitting information, we further constructed a directed and weighted interregional macroscopic connectome based on the axonal projection of E and I neurons. We defined the macroscale connection strength (CS) from a source to a target region as follows: first, we calculated the total length of coherent axonal arbors with terminals in the target region and normalized this value by the number of sampled neurons in the source region. This normalized value was then scaled by the complete cell count of the source region (see Methods). The retina was not included due to the lack of cell count. This approach produced a quantitative macroscopic connectome for the zebrafish nervous system.

We conducted the analysis on 53 anatomical regions, in which the cranial motor nuclei were merged into higher-level brain regions, and the rostral and caudal islands of the same anatomical structures were combined. The wiring diagrams were generated separately for E and I neurons, presenting both ipsilateral/contralateral and ascending/descending connections (Figure 8A). These diagrams revealed several prominent features. First, the macroscale connection strength in both E and I networks vary by a factor of 10^6^ across the nervous system and follow a one-component lognormal distribution (Figure 8B). Second, although a weak negative correlation was observed between connection strength and distance for both E (*P* = 0.04) and I (*P* < 0.001) interregional connections, the effect sizes are minimal (R^2^ = 0.003 for E, and R^2^ = 0.02 for I), indicating negligible associations (Figure 8C). This suggests that long-distance connections are targeted rather than randomly formed. Third, the E dominance, calculated as (CS_E_ − CS_I_)/(CS_E_ + CS_I_), is common for both ipsilateral and contralateral connections in both descending and ascending directions, and applies to both inter-brain region (B-B) connections and mutual brain-spinal cord (B-S) connections (Figures 8A and 8D). For B-B connections in particular, the degree of E dominance is greater for descending than for ascending connections. Fourth, both E and I descending connections exhibit a stronger ipsilateral preference, calculated as (CS_Ipsi_ − CS_Contra_)/(CS_Ipsi_ + CS_Contra_), than ascending ones, and B-B connections prefer ipsilateral connections more than B-S connections (Figures 8A and 8E).

**Figure 8.**
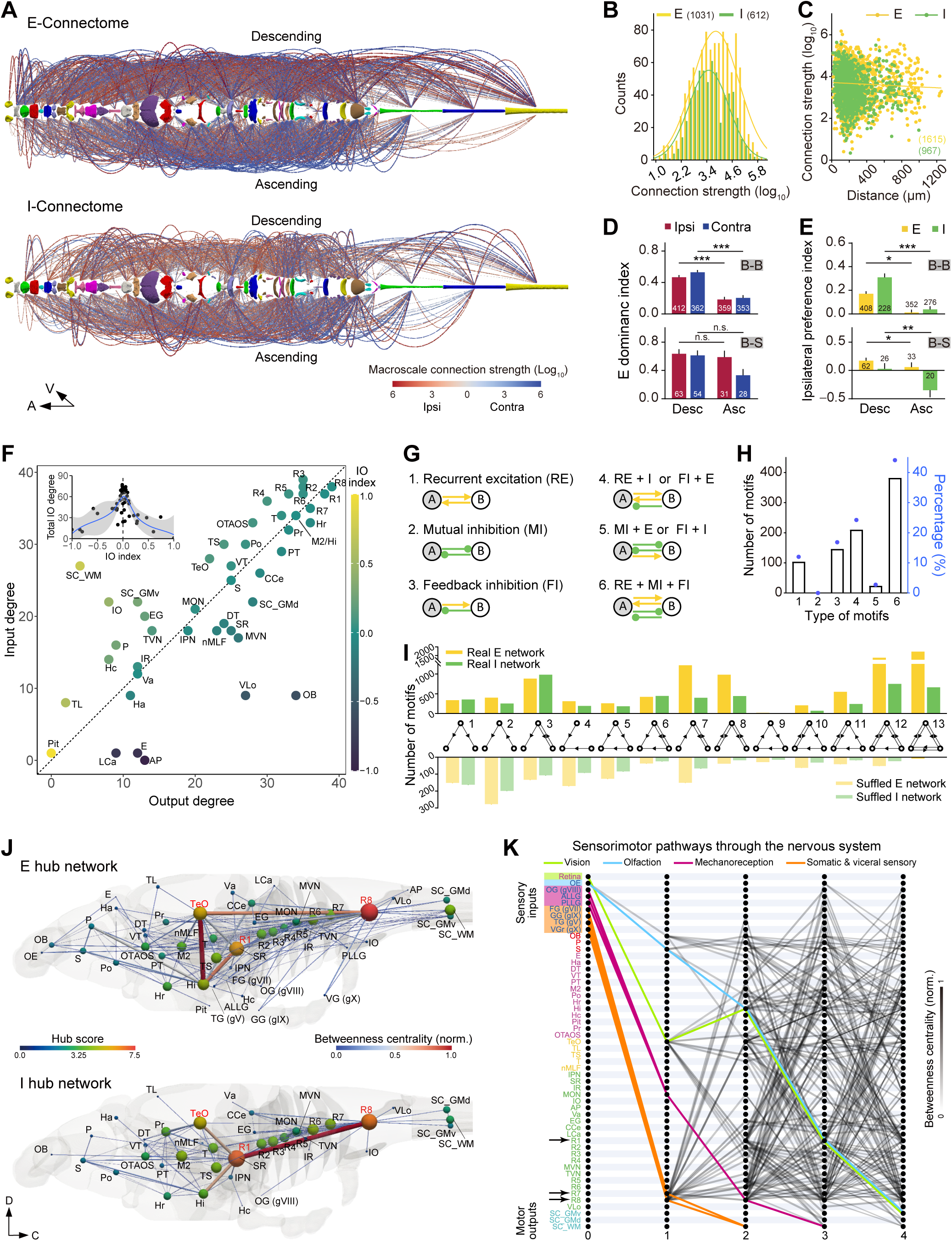
E and I neuron-based interregional connectomes supporting analyses of network motif, hub, and information flow. (A) Wiring diagram across 53 anatomical regions, based on the morphology and cytoarchitecture of E or I neurons (see Methods). Parabolas represent directed ipsilateral (red) and contralateral (blue) connection between regions, which are arranged linearly based on their anteroposterior centroid coordinates. Upper parabolas indicate descending connections, while lower parabolas indicate ascending connections. The parabola’s height and color tone both represent macroscale connection strength (CS). (B) Histogram showing distribution of log-scaled interregional connection strength for E (yellow) or I (green) neurons, each fitted by a Gaussian curve (solid lines; E neurons: R^2^ = 0.91; I neurons: R^2^ = 0.89). (C) Correlation between interregional distance and directed connection strength. E neurons: *r* = −0.05, *P* = 0.04; I neurons: *r* = −0.13, *P* < 0.001. (D) Summary of E dominance index, calculated as (CS_E_−CS_I_)/(CS_E_+CS_I_), in ipsilateral (red) or contralateral (blue) descending and ascending interregional connections. B-B, inter-brain regions; B-S, mutual brain-spinal cord. (E) Summary of ipsilateral preference index, calculated as (CS_Ipsi_−CS_Contra_)/(CS_Ipsi_+CS_Contra_), in E (yellow) or I (green) descending and ascending interregional connections. (F) Relationship between the input (I) degree and output (O) degree for 45 brain and spinal cord regions (excluding the ganglia and retina). The IO index, calculated as (D_I_−D_O_)/(D_I_+D_O_), is color coded. Positive IO index indicates input dominance (receiver), and negative one indicates output dominance (sender). Points falling to the left and right of the 45° diagonal indicate predominantly receiving and sending roles, respectively. Inset: scatterplot of IO index *vs*. total IO degree, fitted by a Gaussian distribution (R^2^ = 0.77). (G) Schematic of six two-node motifs with ≥2 directed connections with E and I identities. Yellow, E connection; Green, I connection. A and B represent any two regions. (H) Numbers (bar) and percentages (dot) of the two-node motifs in (G) in the interregional network. (I) Counts of 13 three-node motifs in the E (yellow) or I (green) interregional networks (top) and corresponding shuffled networks with the same IO degree distribution (bottom). Motif distributions in real networks significantly differ from those in shuffled networks, where expected counts were derived from the mean across 1,000 shuffles and scaled to match the total motif count of the real network. Real *vs*. shuffled E: *Χ*^2^ = 15,564, *P* < 0.001; Real *vs*. shuffled I: *Χ*^2^ = 11,479, *P* < 0.001. (J) Constructed E and I hub networks. Each node (sphere) represents one of 53 regions (retina excluded), with its color and size indicating its hub score. R8, R1, and TeO are the top three hub regions for both E and I hub network. The color-coded edge width (normalized betweenness centrality) reflects its importance in the information flow among regions. (K) Flat connectivity map (based on the E hub network in J) showing sensorimotor pathways from the retina and peripheral sensory ganglia to the spinal cord within 4 steps. The shortest pathway from the starter sensory region to the spinal cord is highlighted with colors. The retina-to-TeO connection is knowledge-based. Arrows point to the premotor hubs R1, R7, or R8. A, anterior; C, caudal; D, dorsal; V, ventral. Numbers in parentheses (B and C) or on bars (D and E) represent the number of source-target pairs. n.s., not significant; \**P* < 0.05; \*\**P* < 0.01; \*\*\**P* < 0.001 (nonparametric Kruskal-Wallis test with Dunn’s post hoc multiple comparisons in D and E). See also Figure S8 and Video S5.

To further explore the features of this interregional connectome, we applied graph-theoretical methods^67^ to analyze the topological properties of the directed and weighted interregional networks (45 regions, excluding peripheral ganglia due to the lack of input node information). First, we evaluated the node degree, a fundamental network measurement. Given a directed network, we calculated the input degree (D_I_), output degree (D_O_), and the IO index ((D_I_ − D_O_)/(D_I_ + D_O_)), which reflects node polarity (positive for receivers and convergence, negative for senders and divergence) (Figure 8F). We found a roughly balanced distribution between receivers and senders. Specifically, 22 nodes had an IO index > 0 (i.e., positioned to the left of the 45-degree diagonal), 4 nodes had an IO index = 0, and 19 nodes had an IO index < 0 (i.e., positioned to the right of the 45-degree diagonal) (Figure 8F). In addition, nodes with higher total IO degrees exhibited a more balanced input-output ratio (IO index around zero) (Figure 8F, inset), indicating a shift from polarized to non-polarized nodes as the number of connections increases.

We next examined two-region motifs (E and I interconnections between nodes) by applying the concept of circuit motifs involving E and I neurons.^68^ We identified six distinct motifs involving at least two directed connections (Figure 8G). Quantitative analysis (Figure 8H) revealed several prominent features: (1) the pure recurrent excitation motif (RE, #1) is present but relatively rare (∼12%); (2) the pure mutual inhibition motif (MI, #2) is absent, and even the motif combining MI with a single excitatory connection (MI+E, #5) is uncommon (∼2%); (3) motifs involving feedback inhibition (FI, #3–6) are predominant (collectively ∼88%), indicating that excitatory connections are frequently accompanied by feedback inhibition; and (4) the most prevalent motif (#6), accounting for ∼44% of cases, consists of bidirectional connections for both E and I.

We then examined three-region motifs (three-node connected subgraphs)^69^ to further investigate the structural design. Specifically, we employed an algorithm to calculate the number of all 13 distinct motifs for both E and I networks (Figure 8I). We found that the proportions of motifs in the E and I networks are correlated (Figure 8I, top; R^2^ = 0.6, *P* < 0.01), reinforcing the coordinated organization between E and I connections. Moreover, motifs featuring reciprocal connections (#3, 6–8, and 10–13) account for a substantial proportion (E, 85%; I, 80%). To assess the significance of these motifs, we compared their occurrences in the real networks against shuffled networks that preserve the same IO degree distribution (Figure 8I, bottom). The real networks exhibited significantly different motif occurrence patterns compared with the shuffled ones (Figure 8I; Real *vs*. shuffled E: Χ^2^ = 15,564, *P* < 0.001; Real *vs*. shuffled I: Χ^2^ = 11,479, *P* < 0.001). Notably, the shuffled networks exhibited a markedly lower proportion of motifs containing reciprocal connections (shuffled E, 40%; shuffled I, 33%) compared with the real networks.

To further characterize information flow across the nervous system-wide network, we quantified the hub distribution by calculating nine centrality metrics for each node, including input strength, output strength, input degree, output degree, weighted betweenness centrality, weighted closeness, PageRank, clustering, and vitality (Figure S8A; see Methods). We defined a region as a hub if it scored above the mean across all metrics, and identified hubs for the E, I, and two separate I (GABAergic and glycinergic) networks. t-SNE visualization^70^ of nine network metrics revealed top hubs in the zebrafish nervous system, including the TeO, R1, R6, R7, and R8 (Figure S8B), which are highly connected with both distant and proximal regions (Figure S8C). We then constructed a hub network, in which nodes represent anatomical regions weighted by hub scores, and edges denote directed information flow between regions weighted by betweenness centrality (Figures 8J, S8D, S8E, and Video S5). As the betweenness centrality measures the proportion of shortest paths between all node pairs that pass through an edge, this network provides the most efficient and likely pathways for information transfer from any source to any target region.

Using the E hub network as a framework, we examined sensorimotor information flow from sensory regions to the spinal cord, the motor output terminal (Figure 8K). We observed a convergence of multimodal sensory information, including vision, olfaction, mechanoreception (audition and water flow), and somatic and visceral senses, onto the premotor hubs R1, R7, or R8 in the hindbrain, from which signals are relayed to the spinal cord. The processing depth ranges from 2 to 4 steps: somatic and visceral sensory pathways are processed most rapidly (2 steps), followed by mechanoreception (3 steps), and then vision and olfaction (4 steps) (Figure 8K).

Collectively, the directed, weighted interregional networks reveal key organizing principles of large-scale circuit architecture. E connections dominate the macroscale connectivity landscape, with descending connections showing a stronger ipsilateral bias than ascending ones. At the network topology level, both E and I connections assemble into specific subgraph motifs characterized by feedback inhibition and reciprocal connectivity, indicating coordinated E and I connectivity. From a systems-level perspective, central hubs such as the TeO, R1, and R8 are shared between the E and I networks. Furthermore, multimodal sensorimotor processing occurs in different shortest steps, with R1, R7, and R8 acting as integrative centers that channel diverse sensory information into motor outputs, underscoring a parallel and hierarchical organization of sensorimotor transformations.

## Discussion

The larval zebrafish serves as a unique model organism in neuroscience research, enabling all-optical structural and functional imaging of the entire brain at cellular resolution in awake animals^71^. To decode how this vertebrate brain works, it is important to develop a comprehensive digital zebrafish holographic atlas integrating multimodal information. In the present study, we generate key datasets in a common coordinate framework, comprising: (1) a whole body-wide CPSv1 of 6-dpf larval zebrafish with two original transgenic reference templates and 3D neuroanatomical parcellations (173 regions); (2) a gene-expression-pattern image atlas by incorporating relevant reference templates (702 patterns) into the CPSv1; (3) a nervous system-wide cytoarchitecture atlas of E and I neurons; (4) a single-cell morphology atlas of E and I neurons (20,011 cells), characterized by the nervous system-wide coverage of cell distribution, whole body-wide coverage of neuronal projection, neurotransmitter identity annotation, dendrite-axon annotation, and morphological classification (600 subtypes); and (5) putative mesoscale (inter-cell-type) and macroscale (interregional) connectomes. These datasets, publicly available at https://zebrafish.cn/LM-Atlas/EI, offer a platform for integrating, exploring, and interpreting data at a nervous system-wide scale.

Previous brain templates were usually generated based on tissue autofluorescence^72^ or single gene expression pattern.^36,39,40^ In this study, we employed the multichannel functionality in ANTs to create dual-channel templates. This approach generated two cellular-resolution reference templates within a common coordinate framework, providing more flexible references and bridges for image registration. Furthermore, we expanded the reference template set by employing shape-based average templates directly aligned to the CPSv1 via a single registration step. This approach avoids the conventional practice of averaging multiple individually registered scans and enhances both accuracy and efficiency. We also improved the registration process by decoupling the rigid-affine and warp steps, ensuring well-converged rigid-affine alignment before proceeding to the computationally intensive warp step. Furthermore, we optimized registration parameters, enhancing accuracy for both live-to-live and live-to-fixed samples. Using gold-standard landmarks and individually identifiable neurons within dense neuronal regions as reference points, we achieved near cellular-resolution precision for inter-atlas (∼8 μm, see Figure S2K) and single-cell morphology atlas registrations (∼6 μm, see Figures S3F–S3I). Beyond well-resolved nervous system structures, autofluorescence from the fish body enables a complete spatial layout of the organism. Future developments, including increased imaging depth and incorporation of visceral structures, will extend CPSv1 into a more comprehensive whole-body framework. This will support the building of projectome and connectome atlases spanning the central, peripheral (somatic and autonomic), and even enteric nervous systems.^73^ Our single-neuron morphology mapping follows a genetically targeted strategy, focusing on the largest and most fundamental neuronal populations: E and I neurons. We selected four key gene markers—*vglut2a*, *vglut2b*, *gad1b*, and *glyt2*—which collectively represent the majority of E and I neurons.^25,74–76^ Future studies could further refine subsets of E and I neurons using intersectional genetic tools^23^ informed by single-cell RNA-seq.^77^ As an immediately accessible next step, the single-neuron morphology atlas could be expanded to include neuromodulatory, neuropeptidergic, and hormone-secreting neurons.

Our study establishes a standardized pipeline for constructing atlases of genetically defined neuronal populations — from reference template generation to cell counting to morphology reconstruction, facilitating systematic characterization of neuronal distribution and morphology. Co-transmission has emerged as a fundamental neural signaling mechanism,^78^ with evidence suggesting widespread co-release of GABA or glutamate with catecholaminergic substances.^74^ Consistently, we identified homologous cell types between published single-cell morphologies of neuromodulatory/neuropeptidergic neurons and our morphotypes. Examples include posterior tuberculum/hypothalamus dopaminergic neurons,^79^ hypothalamus neuropeptide QRFP neurons,^80^ and preoptic oxytocinergic neurons^81,82^ within our *vglut2a/2b* excitatory morphotypes, as well as locus coeruleus norepinephrinergic neurons^79^ and superior raphe serotonergic neurons^83,84^ within *gad1b* inhibitory morphotypes. Overall, our single E and I neuron atlas provides a valuable repository for mining potential co-transmission neuronal types.

To analyze neuron projection features, build wiring diagrams, and infer information flow across the nervous system, we classified the dendrite-axon processes of reconstructed E and I neurons by combining bio-labeling and machine-learning. Utilizing data with co-labeled presynaptic sites and morphology, we summarized three major neurite origin and branching strategies in zebrafish neurons. Notably, a pseudounipolar-like strategy, in which primary axons frequently originate from dendritic branches rather than directly from the soma, is prominent, as exemplified by tectal projection neurons. The trained classifier performed robustly for neuron types with clear polarity, particularly long-range projection neurons. Further refinement of the classification algorithm in a cell-type-specific manner will enhance accuracy.

Neuronal morphology, including dendritic and axonal configurations, is a fundamental criterion for cell type classification, as it underlies connectivity specificity and functional diversity in neural circuits. Morphology-based cell-type classification provides insights into organizational principles governing neuronal projection architecture. One principle is subtype-level modularity, in which bilaterally innervating neuronal types can achieve overall symmetry through a combination of distinct ipsilateral and contralateral projection modes, with E and I neurons exhibiting different preferences for these modes (see Figures 6H–6J and 6L). This organization likely reflects the interplay between genetic programs and environmental influences during neuronal morphogenesis and may contribute to brain lateralization.^85,86^ Variations in the balance of ipsilateral and contralateral projections across individuals and species could shape structural correlates of hemispheric asymmetry and individuality. Because E and I neuronal populations participate in distinct circuit motifs and large-scale architectures,^68^ differences in projection laterality may further contribute to hemispheric specialization of neural circuits. Investigating these projection patterns within functional circuits will be of great interest. A second principle is neuronal type-level modularity, in which related neuronal types share a conserved core projection trajectory while diverge in their dendritic organization or additional axonal projections that extend to distinct targets (see Figures 6A–6G and 6K). Neuronal types with partial overlap yet distinct projections may operate cooperatively within circuits, forming modular projection groups that coordinately innervate multiple brain regions. Structurally, this modularity suggests that new types may emerge through incremental modifications of existing projection patterns, providing a potential developmental mechanism by which neuronal diversity evolves from conserved morphological frameworks. In addition, neuronal morphology is closely linked to spatial distribution, with distinct neuronal types localized to specific brain subdivisions (see Figures 5D and S6). This regional specialization aligns with the idea that neuronal types are structurally and functionally tailored to specific brain areas, underscoring the role of regional organization of the brain.

Our polarity-annotated neuron datasets enable the construction of directed, weighted putative connectome at both inter-cell-type (mesoscale) and interregional (macroscale) levels (see Figures 7 and 8). The inter-cell-type connectome, derived from neurotransmitter-annotated neuronal morphologies under a reliable dendrite-axon proximity threshold, provides insights into circuit mechanisms. As exemplified in Figure 7, we identified a classical visuomotor circuit as well as an understudied circuit underlying phototactic behaviors. These highlight the value of the directed, weighted mesoscopic connectome in mining novel circuits and bridging structural connectivity with potential functions. In terms of the interregional connectome, two features of E and I connectivity are prominent: (1) E connections dominate interregional communication, and (2) E and I connections are co-organized, forming similar subgraph motifs and featuring feedback inhibition across networks. Together, these findings provide a fundamental framework for understanding E and I network architectures.

Beyond documenting connectivity features, we transformed the interregional connectivity network into a hub network by assessing both regional and connection centrality. Separate hub networks were constructed for E and I neurons, highlighting their roles in information flow in the process of sensorimotor transformation (see Figures 8J, 8K, S8D, and S8E). Key findings include: (1) a strong correlation between hub regions and dendrite/axon distribution hotspots (see Figures 4H, 4I, and S8B); (2) shared top-ranking hubs in both E and I networks, notably the TeO, R1, and R8; (3) high hub scores across all hindbrain rhombomeres (see Figure S8B), supporting the notion of the hindbrain as a continuous premotor-coordinating structure; (4) convergent sensorimotor transformation areas, specifically R1, R7, and R8 (see Figure 8K), aligning with those found in brain-wide neural activity maps related to perception and behavior^39,48,49^. In addition, we mapped the shortest sensory processing pathways leading to motor outputs in the spinal cord. Notably, visual and olfactory modalities involve more intermediate steps before motor execution compared with somatic/visceral and mechanoreceptive modalities, a finding that aligns with the hierarchy of sensory experience.^87^

Our study identified hundreds of E and I morphotypes in zebrafish. Integrating transcriptomic datasets with morphologically characterized individual neurons will refine this classification by incorporating genetic determinants.^88^ Application of targeted trans-monosynaptic viral tracing^89–92^ will amass data on precise neuronal connectivity within specific circuits. Through our nervous system-wide CPSv1, comprehensive multimodal mesoscopic atlases can be achieved by integrating molecular, structural (single-neuron/circuit tracing), chemical (neurotransmitter activity), and behavior-related neuronal activity maps.^37^ As increasingly EM datasets emerge,^32,33^ mesoscopic atlases will play a key role in annotating neuronal identities within these datasets.^37^ In the coming future, aligning multimodal mesoscopic atlases with zebrafish EM connectome data will enable the creation of a molecule-structure-function integrated connectome with precise cell-type identity and neuronal connectivity, providing an important framework for zebrafish-based neuroscience research.

### Limitations of the study

While this study provides a nervous system-wide single-neuron morphology atlas of E and I neurons, along with putative inter-cell-type and interregional connectomes for larval zebrafish, several limitations should be noted. First, cytoarchitecture analyses relied on genetically encoded fluorescent markers for cell identification and manual counting. The accuracy was constrained by transgene expression properties, including variability in promoter activity, variegated UAS expression, and limited resolution for defining individual cell boundaries with non-nuclear labeling. Second, current understanding of zebrafish neuroanatomy remains incomplete, and the brain region parcellation employed here was relatively coarse. As neuroanatomical definitions can be refined in the future, our projection maps and connectivity analyses will gain greater functional interpretability. Finally, inter-type connectivity was inferred computationally based on the spatial proximity between annotated dendritic and axonal branch segments. This approach depended on the accuracy of registration, neurite reconstruction, dendrite-axon polarity assignment, and the assumption that thresholded dendrite-axon physical proximity represents synaptic contact. Future integration with ultrastructural EM, transcriptomic profiling, intersectional genetic strategies, and functional interrogation will validate and extend this morphology atlas and the associated connectivity framework.

## RESOURCE AVAILABILITY

### Materials Availability

Plasmids and BAC DNA constructs generated in this study will be deposited to Addgene upon publication and are available upon request. Transgenic zebrafish lines generated in this study have been deposited to China Zebrafish Resource Center (CZRC) (accession pending) and are available upon request.

### Data and code availability

All datasets, including reference transgenic line images, neuroanatomical parcellations, cytoarchitectures, and neuron reconstructions, have been deposited in the Brain Science Data Center (CAS) and will be available upon publication (https://zebrafish.cn/LM-Atlas/download). Inter-atlas transformation matrices will be provided at the same link. All source codes and algorithms for neuronal polarity identification, morphology-based neuron classification, network motif and hub analysis are available at https://github.com/soaringdu/Proj-AtlasEI. Data visualization was performed using integrated <u>Br</u>ain-data <u>A</u>nalysis and <u>V</u>isualization <u>E</u>ngine (iBrAVE),^37^ a custom software built on the open-source ParaView and available at https://github.com/soaringdu/Proj-iBrAVE. iBrAVE supports visualization and analysis of neuron spatial distribution, morphological characterization, interhemispheric crossings, neuron-to-region projection strength, neuron clustering, putative neuronal wiring, and interregional connectivity.

## ACKNOWLEDGMENTS

We are grateful to Drs. Shin-ichi Higashijima and Wei Hu for sharing the BAC and plasmid DNA, Misha Ahrens, Joseph Fetcho, Bo Zhang, and Marnie Halpern for sharing fish lines, and Herwig Baier, Harold Burgess, Florian Engert, and Alexander Schier for sharing line atlas data. We would like to thank Florian Engert for feedback on the brain parcellation and Harold Burgess for feedback on the evaluation of registration precision using landmarks. We would like to thank Ji-Wen Bu, Ye Hua, and Shi-Rong Jin for help with fish husbandry and maintenance; Zhi Zeng and Jian He for providing reconstruction assistance; Ying-Ling Chen, Ying-Ying Wu, Chen-Yan Li, Hua-Qing Shi, and Hong-Yu Li for additional reconstruction work; Ying-Jie Ma and Ying Wang for providing imaging and brain parcellation assistance; Jia-Fei Wei for help with data analysis; and Xing Li for help in the early stages of implementing the interactive web portal. This work utilized the high-performance computing capabilities of the Linux cluster at the Center for Data and Computing in Brain Science at CEBSIT. This work was supported by Brain Science and Brain-like Intelligence Technology - National Science and Technology Major Project (2021ZD0204500, 2021ZD0204502, and 2022ZD0209600), the National Natural Science Foundation of China (32321003, 62320106010, and 62276253), the National Key R&D Program of China (2018YFA0801001), Shanghai Municipal Science and Technology Major Project (2025SHZDZX025D03 and 25511102500), and Key Research Program of Frontier Sciences (QYZDYSSW-SMC028) and Strategic Priority Research Program (XDB32010200) of Chinese Academy of Sciences. X.F.D. is also supported by the Youth Innovation Promotion Association, Chinese Academy of Sciences.

## AUTHOR CONTRIBUTIONS

J.L.D. conceptualized the project. X.F.D. and J.L.D. conceived the experiments and analysis and wrote the manuscript with input from P.J. and Y.F.W., X.F.D. managed the project with input from J.L.D and Z.F.Y.. Z.F.Y. and X.F.D. managed the development of the ZExplorer online platform. Z.F.Y. developed the two-channel template generation and high-precision alignment of 3D images required for atlasing transgene, cell-counting, and neuron tracing data. W.L.L. developed the methods for neuron polarity identification, neuron morphological classification, and network analysis with input from X.F.D. and Z.F.Y.. Z.F.Y., T.L.C., and L.J.C. performed the registration precision evaluation. X.F.D. and M.Q.C. conducted the brain parcellation and annotation with input from Z.F.Y., Y.F.W., D.Y.L., L.J.C., S.J.W., Y.C.G., T.L.C., and Q.M.Z.. X.Y.N. conducted the nervous system-wide cell counting. H.Y.H., X.D.Z., and Y.L. developed genetic tools for transient microinjection and transgenic line generation. Z.M.J. and X.D.Z. conducted the microinjection and sample pre-screening with input from X.F.D., X.L.S., X.Y.Q., and W.Z.. H.L.W., H.A.R., R.Q.W., R.Z., X.L.P., L.W., M.Y.Z., T.L.C., and X.Y.S. conducted the sample preparation and acquired the live imaging data with input from W.Z., X.F.D., X.L.S, and Q.M.Z.. J.H. supervised Y.L., X.Y.Q., and X.Y.S.. Y.M. supervised D.Y.L. and S.J.W.. T.T.Z., Y.W.Z., and X.Y.N. conducted the neuron reconstruction with input from X.F.D. and X.L.S.. C.X.J. produced the videos about the experimental pipeline and designed the web interface with input from X.F.D., Y.F.W., and J.L.D.. J.W.H., W.D., and L.Q.Q. conducted the implementation of the interactive web portal with input from Z.F.Y.. X.F.D. conducted the analyses and made the illustrations with input from M.Q.C., W.L.L., T.L.C., P.J., X.Y.N., and Y.W.Z..

## DECLARATION OF INTERESTS

The authors declare no competing interests.

## METHODS

### Zebrafish husbandry and preparation

Adult zebrafish were maintained at 28 ℃ in an automatic fish housing system at the National Zebrafish Resources of China (Shanghai, China). All image data were collected from transgenic larval zebrafish at 6 days post-fertilization (dpf), a stage at which sex has not yet been determined. Experimental larvae were raised in 10% Hank’s solution consisted of (in mM): 140 NaCl, 5.4 KCl, 0.25 Na_2_HPO_4_, 0.44 KH_2_PO_4_, 1.3 CaCl_2_, 1.0 MgSO_4_ and 4.2 NaHCO_3_ (pH 7.2) under a 14-h:10-h light:dark cycle at 28℃. Larvae lacking the *nacre* genetic background were treated with 0.003% 1-phenyl-2-thiourea (PTU) starting from 8–22 hours post-fertilization to prevent pigmentation. All animal protocols were reviewed and approved by the Animal Care and Use Committee of the Center for Excellence in Brain Science and Intelligence Technology, Chinese Academy of Sciences (NA-046-2023).

### Genetic tools

To label E and I neurons, we generated DNA constructs and transgenic lines using either short promotor or BAC clone. For *vglut2a:GAL4FF* and *glyt2:GAL4FF*, we PCR amplified the *GAL4FF* fragment with a Kozak sequence and restriction enzyme digestion sites and inserted it into the *Tol2* vector with ∼2.2 kb *vglut2a* (a gift from Dr. Wei Hu, Institute of hydrobiology, CAS, Wuhan, China) and ∼6.8 kb *glyt2*^93^ promotor, respectively. The *cpce:GAL4FF* construct for generating transgenic fish was previously reported.^92^ For *vglut2b:GAL4FF* and *gad1b:GAL4-VP16*, we used BAC recombineering as described previously.^94^ First, the BAC clone was transformed into SW105 bacterial strain to enable the heat-inducible homologous recombination. Second, *iTol2* arms in opposing directions flanking an ampicillin resistance cassette were PCR amplified from the *iTol2* plasmid and introduced into the BAC backbone. Third, the reporter cassette was PCR amplified from the constructed *pIGCN21-Left recombination arm-Reporter/PolyA-FRT/Neo/FRT-Right recombination arm* and inserted into the BAC to replace the first exon or that with partial intron I of interested gene. Finally, the successful insertion of the reporter cassette was confirmed by PCR and sequencing. BAC clone sources, primer sequences, and insertion sites of the reporter cassette are listed in Table S3. The *TgBAC(vglut2a:LOXP-DsRed-LOXP-GFP)ion18h* line was generated using a previously reported construct.^95^

For the generation of *14×UAS-E1b:tdTomato-CAAX*, the tandem-dimer tomato (tdT) was PCR amplified with a 3’ membrane tag encoding the *CAAX* box of human Harvey Ras (CTGAACCCTCCTGATGAGAGTGGCCCCGGCTGCATGAGCTGCAAGTGTGTGCTCTCC) and cloned into the *miniTol2-14*×*UAS-E1b* backbone from plasmid *miniTol2-UAS:GCaMP5*.^96,97^ To label cerebellar cells, we generated the *Tg(5×UAS-hsp:lyn-tdTomato)* using zebrafish codon-optimized *lyn* as a membrane-bound signal.^98^ For the generation of *14×UAS-E1b:sypb-EGFP,14×UAS-E1b:tdKatushka2-CAAX*, we cloned the PCR amplified *sypb-EGFP*^99^ and *de novo* synthesized *tdKatushka2-CAAX* to the two sides of a bidirectional *14*×*UAS-E1b* in the *miniTol2* backbone.

### Sparse labeling of single neurons

Sparse single-neuron labeling was achieved through transient expression of the membrane-targeted reporter *14×UAS-E1b:tdTomato-CAAX*, delivered together with *Tol2* transposase mRNA into fertilized embryos carrying *Tg(elavl3:H2B-GCaMP6s)* for subsequent whole-brain registration. Reporter activation was driven either by co-injection of a GAL4/GAL4FF activator plasmid or by endogenous GAL4/GAL4FF transgene present in the embryos (see Key Resources Table). This transient transgenic approach produced stable fluorescent protein expression in only one or a few neurons that are random subsets of the overall GAL4/GAL4FF expression pattern. Labeling density can be adjusted by titrating the plasmid injection amount.

For *vglut2a*+ excitatory neurons, v*glut2a:GAL4FF*, *UAS* reporter, and *Tol2* mRNA were injected at either ∼(7 + 7 + 7) pg into one blastomere at the 16-cell stage or ∼(2 + 2 + 2) pg at 1-cell stage. For *vglut2b*+ excitatory neurons, *BACvglut2b:GAL4FF*, *UAS* reporter, and *Tol2* mRNA were injected at ∼(7 + 7 + 7) pg into one 16-cell blastomere or ∼(3 + 3 + 3) pg at the 1-cell stage. For *gad1b*+ inhibitory neurons, *BACgad1b:GAL4-VP16*, *UAS* reporter, and *Tol2* mRNA were injected at ∼(9 + 9 + 9) pg into one 16-cell blastomere or ∼(3 + 3 + 3) pg at the 1-cell stage. For *glyt2*+ inhibitory neurons, *glyt2:GAL4FF*, *UAS* reporter, and *Tol2* mRNA were injected at ∼(10 + 10 + 10) pg into one 16-cell blastomere or ∼(3 + 3 + 3) pg at the 1-cell stage. For *isl2b*+ RGCs, *UAS* reporter and *Tol2* mRNA were injected at ∼(1.5 + 1.5) pg into *Tg(isl2b:GAL4-VP16)* embryos at the 1-cell stage. For *atoh7*+ RGCs, the same components were injected at ∼(1.7 + 1.7) pg into *Tg(atoh7:GAL4-VP16)*. For *cbln12*+ excitatory and *cpce*+ inhibitory neurons, *Tg(cbln12:GAL4FF)* and *Tg(cpce:GAL4FF)* were crossed to *Tg(elavl3:H2B-GCaMP6s);Tg(5×UAS-hsp:lyn-tdTomato)*, respectively.

Larvae exhibiting tdTomato-positive neurons together with whole-brain nuclear GCaMP6s fluorescence were pre-selected. At 4–5 dpf, fish were embedded in 1.3% low-melting-point agarose and screened under an upright fluorescence setup with water-immersion optics. Only larvae containing clearly resolvable single-neuron labeling were advanced to high-resolution live imaging at 6 dpf.

### Imaging

For live imaging, zebrafish larvae at 6 dpf were anesthetized with 0.02% tricaine methanesulfonate or immobilized with pancuronium dibromide (PCD, 1 mM), and embedded in 2–2.5% low-melting point agarose on a custom-made imaging chamber. Imaging was performed on an Olympus FV3000 or a Nikon A1R upright confocal microscope with a 20×(1.0 N.A.) or a 25× (1.1 N.A.) water-immersion objective, respectively. For reference template construction, three overlapping image stacks along the anteroposterior axis were acquired at a voxel size of ∼ 0.62 × 0.62 × 1 μm (*xyz*), covering the brain and approximately one third of the spinal cord. These were stitched together using either native imaging software or the Grid/Collection stitching plugin in Fiji. For single-neuron reconstruction, larvae were scanned at a voxel size of ∼ 0.49 × 0.49 × 1 μm in a 1 × 2 or 1 × 3 tiled arrays depending on the projection range of the neuron. Laser intensity was progressively increased along the *z*-axis to compensate for fluorescence attenuation. The 405 nm, 488 nm, 561nm, and 594 nm diode-pumped solid state lasers were used to excite fluorophores of TagBFP, GCaMP6s/EGFP, tdTomato, and tdKatushka2, respectively.

### Template generation and image registration

To generate the original dual-channel template *elavl3:H2B-GCaMP6s;vglut2a:DsRed*, we used the open-source Advanced Normalization Tools (ANTs) software running on the CEBSIT’s Linux computing cluster. This software allows the creation of a multichannel template by using the shell script antsMultivariateTemplateConstruction2.sh.^100^ Confocal images were resampled from ∼0.62 × 0.62 × 1 μm (*xyz*) to 1 × 1 × 1 μm resolution before generating the template. In short, the template was created by shape-based averaging from 16 image scans of the double transgenic fish through an iterative process (see Figure S2A). In detail, an average image was first generated from the input fish scans by rigid alignment. Then, an iterative process began, which included deformable registration of each aligned scan to the average image based on the two expression patterns (1×1 weights). The resulting registered scans were then averaged for each expression pattern to form the new average image. This process continued until convergence was attained, as revealed by a growth plateau of the similarity scores between the average templates resulting from adjacent iterations using the normalized cross correlation (NCC)^101^ metric (see Figure S2B; see the section “Registration accuracy evaluation” in Methods).

Other reference templates in the common physical space (CPSv1) were generated similarly by first producing a two-channel, shape-based average, followed by transforming the target channel to the CPSv1 via bridging with the *elavl3:H2B-GCaMP6s* or *vglut2a:DsRed* channel using the antsApplyTransforms function in ANTs. The number of fish used, iteration cycles, and bridging patterns for other reference templates in CPSv1 are listed in Figure S2C. Additional templates, such as *isl2b* and *atoh7:GAL4* lines crossed to *Tg(4×nrUAS:GFP)* for labeling RGCs, were integrated into CPSv1 via the *elavl3:NES-jRGECO1a* bridge template.

Image registration was performed using the antsRegistration function in ANTs,^38^ with optimized parameters adapted from a previous study^44^ for improved accuracy, particularly between live and fixed fish scans (see the section “Registration accuracy evaluation” in Methods). Registration proceeded in two sequential steps using Slurm-based bash scripts: (1) rigid and affine (RA) registration (translation, rotation, anisotropic scale and shear), and (2) nonlinear warp registration. This two-step design allows RA transforms to be evaluated first, so poorly aligned samples can be excluded before the computationally intensive warp step, thereby greatly improving overall efficiency for large-scale batch-mode datasets. To improve registration accuracy, a fish-body mask, created using CMTK’s *levelset* and refined manually, was applied to both the reference and RA-registered images during warp registration to remove background fluorescence.

For most neuron-tracing datasets, the *elavl3:H2B-GCaMP6s* expression pattern in the CPSv1 was used as the registration template. This also applied to datasets expressing Sypb-EGFP-tdKatushka2-CAAX, in which the *elavl3:H2B-TagBFP* transgene served as the reference channel, because it exhibits the same nucleus-localized expression pattern as *elavl3:H2B-GCaMP6s*. For RGC datasets, a composite template (*Tg(elavl3:H2B-GCaMP6s)* and *Tg(isl2b/atoh7:GAL4);Tg(4×nrUAS:GFP)*) was used to enhance alignment accuracy in the tectal neuropil area.

The ANTs commands used are listed below:

1. Two-channel template generation: antsMultivariateTemplateConstruction2.sh -d 3 -b 1 -g 0.1 -i 10 -c 2 -j 16 -k 2 -w 1x1 -r 1 -n 0 -m CC -t SyN ${image-inputs}
2. Rigid and affine registration: antsRegistration --verbose 1--dimentionality 3 --float 1 --collapse-output-transforms 1 --output [${image-output}ra,${image-output}ra_warped.nrrd] --interpolation WelchWindowedSinc --use-histogram-matching 0 --initial-moving-transform [${template}, ${image-input},1] --transform Rigid[0.1] --metric MI[${template},${ image-input},1,32,Regular,0.25] --convergence [400 × 200 × 200 × 20,1e-8,10] --shrink-factors 12×8×4×2 --smoothing-sigmas 4×3×2×1vox -transform Affine[0.1] --metric MI[${template}, ${image-input},1,32,Regular,0.25] --convergence [400×200×200×20,1e-8,10] --shrink-factors 12×8×4×2 --smoothing-sigmas 4×3×2×1vox
3. Warp registration: antsRegistration -d 3 --float 1 --verbose 1 -o [${image-output}SyN,${image-output}SyN_warped.nrrd] -interpolation WelchWindowedSinc -use-histogram-matching 0 -r [${mask},${mask},1] -t SyN[0.2,6,0] -m CC[${template},${image-output}ra_warped.nrrd,1,2] -c [400 × 200 × 100 × 50 × 0,1e-7,10] -f 12×8×4×2×1 -s 5×3×2×1×0vox -x [${mask},${mask}]
4. Apply transformation matrix to the image in a second channel: antsApplyTransforms -d 3 --float -n WelchWindowedSinc -i ${image-input_second channel} -r ${template} -t ${image-output}SyN1Warp.nii.gz -t ${image-output}ra0GenericAffine.mat -o ${image-output_second channel}warped.nii.gz

### Registration accuracy evaluation

We assessed the registration precision using the MLD metric.^44^ Taking the *elavl3:H2B-GCAMP6s* expression pattern as an example, we identified 13 widely distributed landmarks across the zebrafish brain (see Figure S3A–S3C). Each landmark was verified for recognizability by three independent annotators and had a mean positional deviation ranging from 1.69 to 5.10 μm (mean = 3.10 ± 1.10 μm). We manually marked the landmarks on both the reference template and unregistered individual scans using ImageJ. For each scan, a secondary mask channel was created by embedding 3-pixel cubes centered at each landmark position. After registering individual scans to the reference template, we applied the resulting transformation matrix to the mask channel. The landmark distance (LD) was calculated as the Euclidean distance between the geometric center of corresponding cubes in the registered and reference stacks. The MLD for each registered scan was calculated as the average of its 13 LD values.

For evaluating inter-atlas registration accuracy, we applied six expression patterns: *elavl3:H2B-FPs* (*FPs* include *GCaMP6s* and *RFP*), *elavl3:GECIs* (*GECIs* include *GCaMP6s*, *NES-jRGECO1a*, *GCaMP5G*, and *CaMPARI*), *vglut2a:DsRed*, *gad1b:EGFP*, *glyt2:GFP*, and *vmat2:GFP*. For each pattern, we marked 6–9 landmarks (45 in total) in the corresponding reference templates in the Z-Brain, ZBB, mapzebrain, and our own ZExplorer atlas (see Figure S2H and S2I). Not all six expression patterns were available for each comparison. The co-registration precision was computed as the mean of LDs measured between matched expression patterns (see Figure S2K).

For evaluating the registration accuracy of single-neuron morphology atlas, we adopted two strategies. First, we applied a landmark-based approach by randomly selecting 88 *elavl3:H2B-GCAMP6s* fish scans and calculating MLDs (see Figure S3F). Second, we quantified 3D soma-to-soma distances of individually identifiable Mauthner cells using two independent datasets. For spinal dye backfill-labeled Mauthner cells, corresponding Mauthner cell pairs were manually identified in two fish. Their registration-channel images were registered, with each fish alternately treated as the reference or moving sample. Post-registration distances between matched somata were then calculated (see Figures S3G and S3I). For Mauthner cells reconstructed in the single-neuron morphology atlas, soma coordinates were directly extracted from SWC files, and pairwise soma-to-soma distances were computed for matched neurons from the same hemisphere (see Figures S3H and S3I).

The NCC was used as a rapid, quantitative metric for evaluating batch-level registration accuracy, calculated via the similarity function in the Computational Morphometry Toolkit (CMTK). During registration using ANTs, we measured NCC after performing the RA registration. Scans with NCC_RA_ >0.55 (empirically determined) proceeded to the next step of warp registration (see Figure S3J), while those below the threshold were inspected manually to determine their suitability for subsequent warp registration.

### Brain parcellation

To annotate the nervous system of larval zebrafish in 3D, we manually delineated the brain, partial spinal cord (segments 1–11), and peripheral ganglia into 68 anatomical regions (comprising 173 parcellations). Segmentations was performed using the 3D slicer (https://www.slicer.org), an open-source platform for image processing and 3D visualization, in 3 orthogonal planes (i.e., horizontal, transverse, and sagittal).

Region boundaries were primarily defined based on established neuroanatomical references^22^ and the reported Z-Brain ontology (https://zebrafishatlas.zib.de).^39^ Delineation was guided mainly by expression patterns of three key reference templates generated by ourselves (*elavl3:H2B-GCaMP6s*, *vglut2a:DsRed*, and *gad1b:EGFP*), as well as other transgenic line data incorporated in the CPSv1. Informative expression patterns were visualized in semi-transparent overlays, and each segmentation was illustrated in coordination with the surrounding ones. In addition, the medial octavolateralis nucleus (MON) was uniquely delineated based on afferent projections from octaval (gVIII) and lateral line (ALLG and PLLG) neurons in our single-neuron morphology atlas.

Initial segmentations were conducted in the horizontal (*xy*) plane, followed by iterative refinements across all three planes to ensure anatomical accuracy and smooth boundaries. The final parcellations were jointly smoothed using a custom multiple 3D median filter, resulting in a comprehensive, non-overlapping, and gap-free set of nervous system-wide regions. The complete ontology of annotated regions in the CPSv1 is listed in Table S2.

### Cell counting and registration

To construct a nervous system-wide neuronal cytoarchitecture, individual somata were manually annotated based on the centroid of fluorescent signals in 3D image stacks using the Cell Counter plugin in Fiji. We mapped five neuronal populations: *elavl3*+, *vglut2a*+, *vglut2b*+, *gad1b*+, and *glyt2*+, corresponding to the expression patterns of *elavl3:H2B-GCaMP6s*, *vglut2a:DsRed*, *vglut2b:GAL4FF*, *gad1b:EGFP/gad1b:GAL4-VP16*, and *glyt2:GFP*, respectively. For each neuronal population, four individual fish were quantified. Each fish scan was annotated by one researcher and independently reviewed by two others before final confirmation. The counts for *vglut2a*+ and *vglut2b*+ neurons were combined to represent the excitatory neuronal cytoarchitecture.

To register the cytoarchitecture data into the CPSv1, the expression pattern used for cell annotation or the alternative imaging channel that matches one of the two original reference templates was first registered to the CPSv1. The resulting transformation matrix was then applied to the digital cytoarchitecture data (a file containing the coordinates of annotated somata) using the antsApplyTransformsToPoints function in ANTs:

antsApplyTransformsToPoints -d 3 -i ${point-input} -o ${point-output}warped.csv -t [${image-output}ra0GenericAffine.mat,1]

antsApplyTransformsToPoints -d 3 -i ${point-output}warped.csv -o

${point-output}warped_SyN.csv -t ${image-output}SyN1InverseWarp.nii.gz

### Neuronal morphology reconstruction and registration

Individual neurons suitable for reconstruction were manually selected and semi-automatically traced using the Simple Neurite Tracer (SNT) plugin in Fiji.^102^ The resulting neuron skeletons, composed of interconnected nodes, were saved as individual SWC files.^103^ The soma radius for each neuron was determined by averaging the semi-major and semi-minor axes of the soma in its maximum-intensity-projection image, and this value was written into the SWC file. Each neuron was traced by a single annotator and reviewed in sequence by two experts of increasing expertise. On average, reconstruction of a single neuron requires ∼1.2 hours, with each fish providing ∼1.5 reconstructions (total of 13,429 fish for 20,011 neuron reconstructions). Neurons with deep ventral and/or caudal spinal cord projections extending beyond the imaging volume were marked as incomplete. Their inclusion in analyses was determined on case-by-case basis.

For neurons with presynaptic labeling (Sypb-EGFP), we manually annotated the node closest to the geometric center of each fluorescently discrete Sypb-EGFP punctum as a presynaptic site and assigned the type identifier as 7 in the SWC files. The type identifiers for soma, axon, and dendrite are 1, 2, and 3, respectively.

Prior to registration, each neuron reconstruction was preprocessed using smooth_neuron (σ = 1) and resample (step size = 1) functions in the R package nat. Presynaptic site coordinates were preserved during this step. The preprocessed neuron skeleton was then registered to the CPSv1 using the antsApplyTransformToPoints function in ANTs (see the section “Cell counting” in Methods).

### Neuronal polarity identification

To identify neuronal polarity (i.e., dendritic *vs*. axonal compartments), we trained a machine learning-based classifier using neuron reconstructions with dendrite-axon annotations obtained from morphology-presynaptic site (Sypb-EGFP) double-labeling data. Each neuron was represented as a tree graph, from which six morphological parameters were extracted to train the classifier: (1) terminal branch length, (2) branch point order, (3) path distance from soma to terminals, (4) Euclidean distance from soma to terminals, (5) normalized terminal coordinates relative to soma, and (6) maximum branch length along terminal paths. Because each node along a terminal path shares the same polarity identity as the terminal, we classified only the terminals and assigned all upstream nodes accordingly, resulting in some nodes being shared between axonal and dendritic branches.

We implemented this classification model based on the support vector machine (SVM) and used the reconstruction of 70 neurons to train the classifier, achieving ∼97% accuracy on a test dataset (12 morphotypes, n = 877; see Figures S4 and S5). This automatic classification method enables rapid identification of neuronal polarity for large-scale datasets. Polarity annotation is most reliable for projection neurons with clear structural polarization. Therefore, we left the polarity of most local neurons unassigned, as well as the arborizations of tectal neurons within the tectal neuropil, where dendritic and axonal processes from different fine subtypes are intermingled across multiple laminae. Final results were manually verified.

### Neuron projection analysis

We measured the projection strength of each neuron to target regions using axonal projection length (APL). This measure includes only axon compartments associated with terminals inside the region, excluding those merely passing through the region. To mitigate potential errors caused by registration or parcellation imprecision, we only considered a neuron to project into a region if its axonal length within that region exceeded 10 μm. The neuron cluster-to-region projection strength was calculated by summing the projection strength of all neurons in the cluster and normalizing by the number of cells in the cluster.

### Neuron classification

We performed classification analysis using mirrored data.^36,52^ Briefly, we horizontally flipped the *elavl3:H2B-GCaMP6s* reference template and registered the flipped stack to the original stack using ANTs to obtain a mirroring transformation matrix. Reconstructed and registered neurons were divided into left and right groups based on soma location relative to a mid-sagittal plane, fitted from scattered midline points across horizontal sections of the brain and spinal cord in the CPSv1. Neurons on either side were flipped to the opposite side and registered to the CPSv1 using the antsApplyTransformToPoints function in ANTs with the mirroring matrix. This procedure yielded a set of complete reconstructions for both hemispheres, and we used a set comprising the original left-hemisphere neurons together with the mirrored right-hemisphere neurons for subsequent morphological classification.

We developed an unsupervised learning-based neuron classification algorithm termed MBLAST by using neuron morphology. Requiring no prior information, this algorithm discovers clusters and detects the noise points on large databases efficiently. It consists of two functions: (1) detecting new clusters from neuronal populations; (2) assigning new neurons to existing clusters. The core of the algorithm is a shape-based distance metric inspired by the Hausdorff distance.^104^ We considered each neuron *P* as a trajectory consisting of tracing points with a size of *n_p_*. The directional distance from *P*_1_ to *P*_2_ was defined as:

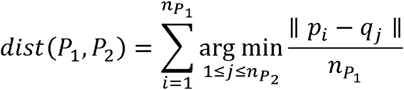

where *p_i_* ∊ *P*_1_, *q_j_* ∊ *P*_2_. The overall distance between two neurons *P*_1_ and *P*_2_ was defined as:

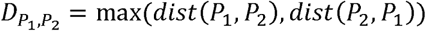

This metric is sufficiently sensitive to capture the similarity of neuron shapes and easy to compute. Besides, this distance score requires no parameter and thus is easily applied to construct a similarity matrix for any neuron reconstruction dataset. For neurons with similar morphology but soma positions shift along anteroposterior axis (e.g., in the hindbrain and spinal cord), coordinates were first normalized to the soma to prevent positional offsets from affecting the distance measure.

Morphological clustering was performed using DBSCAN (Density-Based Spatial Clustering of Applications with Noise),^105^ a density-based algorithm effective in discovering clusters of arbitrary shape. In order to find clusters with different densities, we implemented a multiscale DBSCAN approach, iteratively clustering neurons with different global density parameters. The resulting clusters were validated by visual inspection of individual neurons, which also served as an additional quality control for neuronal reconstruction accuracy, and were manually curated to generate the final set of basic neuron clusters.

To classify newly reconstructed neurons, we developed a probabilistic query method. For each existing cluster *C*, we defined its intra-cluster similarity distribution as *D(C)*. For an unclassified neuron *n*, we computed its distance to each neuron *n_c_* ∊ *C*, and evaluated the probability for *n*, belongs to *C* as: 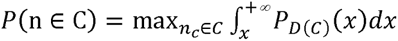, where *x* = *dist*(*n,n_c_*) Neuron *n* was assigned to the cluster with the highest probability above a custom threshold (30% in this study), and then manually reviewed for confirmation.

### Descending/ascending and ipsilateral/contralateral projection analysis of morphological Clusters

First, we calculated the neuron-to-region projection strength *APL_ij_* for each neuron *i*, in morphological cluster *c* projecting to region *j*. Second, we annotated each *APL_ij_* as either 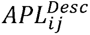 or 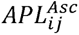 based on the relative anteroposterior position of the neuron soma and the target region centroid. The cluster-to-region descending and ascending projection strength were calculated as 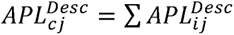 and 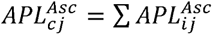, respectively. Then the cluster-to-region descending/ascending preference index was calculated as:

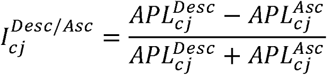

Similarly, each *APL_ij_* was also separated into ipsilateral 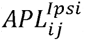 and contralateral 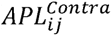 components, depending on whether the projection was to the same or opposite hemisphere as the soma. We calculated the total ipsilateral and contralateral projection strengths from cluster to region as 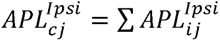 and, 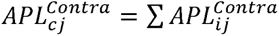, respectively. The cluster-to-region ipsilateral/contralateral preference index was calculated as:

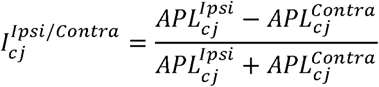

### Putative neuronal connectivity analysis

We utilized axon-dendrite spatial proximity to infer neuronal connections. The distance was measured between segments on axonal and dendritic branches. To ensure specificity, only axonal branches with terminal endings were included. Putative connections were defined when the Euclidean distance between axonal and dendritic branch segment centers fell below a specified threshold (e.g., 2 μm). The connection strength between individual neurons was quantified as the number of such segment pairs (≥1).

The mesoscale connection strength between neuronal clusters was calculated by summing the connection strength of all connected neuron pairs and normalizing by the number of neuron pairs. Two additional metrics were used to filter the connectivity between neuronal clusters: (1) connection ratio, the percentage of connected neurons in each cluster. The larger of the two percentages must be at least 20%; (2) divergence and convergence (D&C) indices, the number of connected neuron pairs divided by the number of participating neurons in input and output clusters, respectively. The D or C index must be larger than 1.

### Interregional connectivity analysis

Interregional connection strength is a concept used to measure the connectivity between any two regions, which can be defined in different ways.^36,106^ Here, we introduce a metric termed “complete axonal projection length (CAPL)”, which estimates the total macroscale connection strength between brain regions by incorporating endogenous neuronal population size, thereby approximating the weighted interregional connectivity. The computation of CAPL involves four main steps:

First, we generated the neuron-to-region projection strength matrix by calculating axonal projection length for each neuron-region pair:

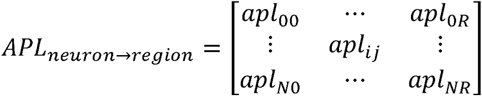

where *N* is the number of sampled neurons *R*, is the number of regions, and *apl_ij_* is the axonal projection length of neuron *i* inside region *j* and calculated as:

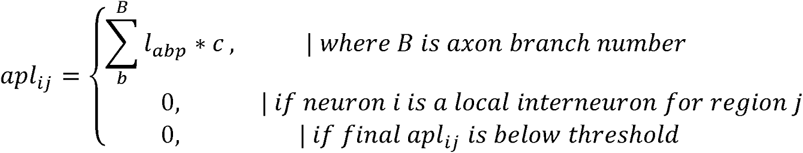

Here, *l_abp_* is the length of axon branches of neuron *i* inside region *j*, and *c* is a coefficient which is 0 when axon branch just passes through the region, or 1 otherwise.

Second, we obtained the region-to-region connection strength by summing the axonal projection lengths of all sampled neurons with soma located in the source region and projecting to the target region:

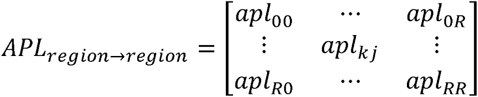

where 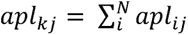, and *N* is the number of neurons inside source region *k*.

Third, we normalized the region-to-region connection strength by dividing the number of sampled neurons inside the source region:

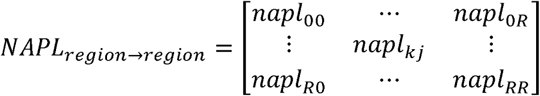

where 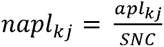, and *SNC* is the number of sampled neurons inside source region *k*.

Finally, we calculated the region-to-region CAPL by scaling the normalized connection strength with the total number of neurons in the source region (from cytoarchitecture data):

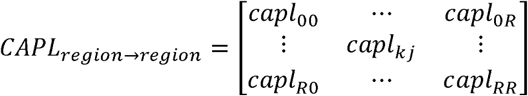

where *capl_k,j_* = *napl_k,j_* * *T* and *T* is the total number of neurons inside source region *k*.

To derive ipsilateral and contralateral CAPL, each neuron’s projection strength was partitioned based on its position relative to the aforementioned mid-sagittal plane. The ipsilateral and contralateral CAPL were then calculated separately using the same procedure as for total CAPL. The interregional ipsilateral/contralateral preference index was calculated as:

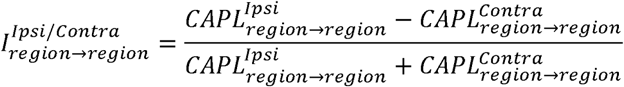

Region pairs were annotated as descending or ascending based on the relative anteroposterior positions of the source and target regions.

### Network hub analysis

Network hubs are considered as nodes in the network with high centrality for information transfer. In this weighted network, we defined the path distance between two nodes as the sum of the reciprocals of directed connection strength along the path. We scored node centrality based on nine metrics: (1) input strength, total connection strength from directly connected upstream nodes; (2) output strength, total connection strength to directly connected downstream nodes; (3) input degree, number of upstream nodes directly connected to the target node; (4) output degree, number of downstream nodes directly connected to the target node; (5) betweenness centrality, frequency with which a node lies on the shortest paths between all other node pairs; (6) closeness, ratio of the number of reachable downstream nodes to the average shortest path distance to those nodes; (7) PageRank, importance of a node based on the structure of its connections, an algorithm originally developed for ranking web pages;^107^ (8) clustering, fraction of bidirectional triangular motifs involving the target node; (9) vitality, change in the total shortest-path distance between all nodes when the target node is removed. Each metric was normalized within its group. The hub score of a node was defined as the sum of its normalized scores across all nine metrics. Nodes with above-mean scores for all metrics were defined as network hubs.

In the interregional hub network, edge width represents the weighted betweenness centrality of directed connections, indicating their influence on the network-wide information flow. Edges with higher values appear more frequently in the shortest paths and serve as key bridges between subnetworks.

### Statistical analysis

No randomization and blinding were used during data collection and analysis as there is a single experimental condition for all acquired data. The statistical analysis was performed using Python or GraphPad Prism. The central mark, bottom, and top of the boxplot indicate the median, first, and third quartiles, respectively. The height and error bars of the bar plots, as well as the descriptive statistics in the main text, represent mean ± SEM. Statistical significance is defined as *P* < 0.05. The specific statistical tests and *P*-value ranges are provided in the corresponding figure legends. Sample sizes (n) refer to the number of samples, fish, or cells, as specified in the relevant figure and/or figure legend.

---

**Figure S1.**
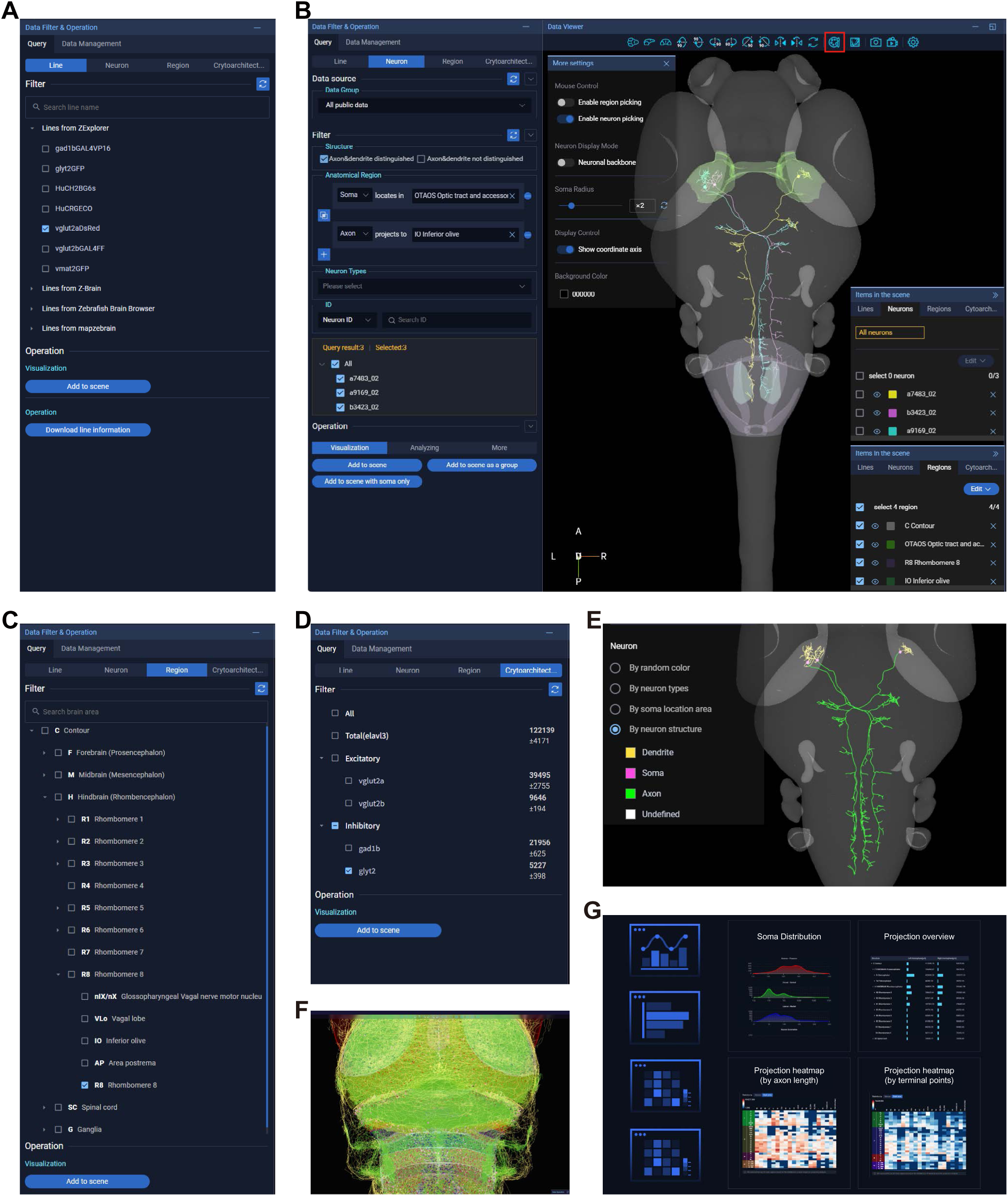

**Figure S2.**
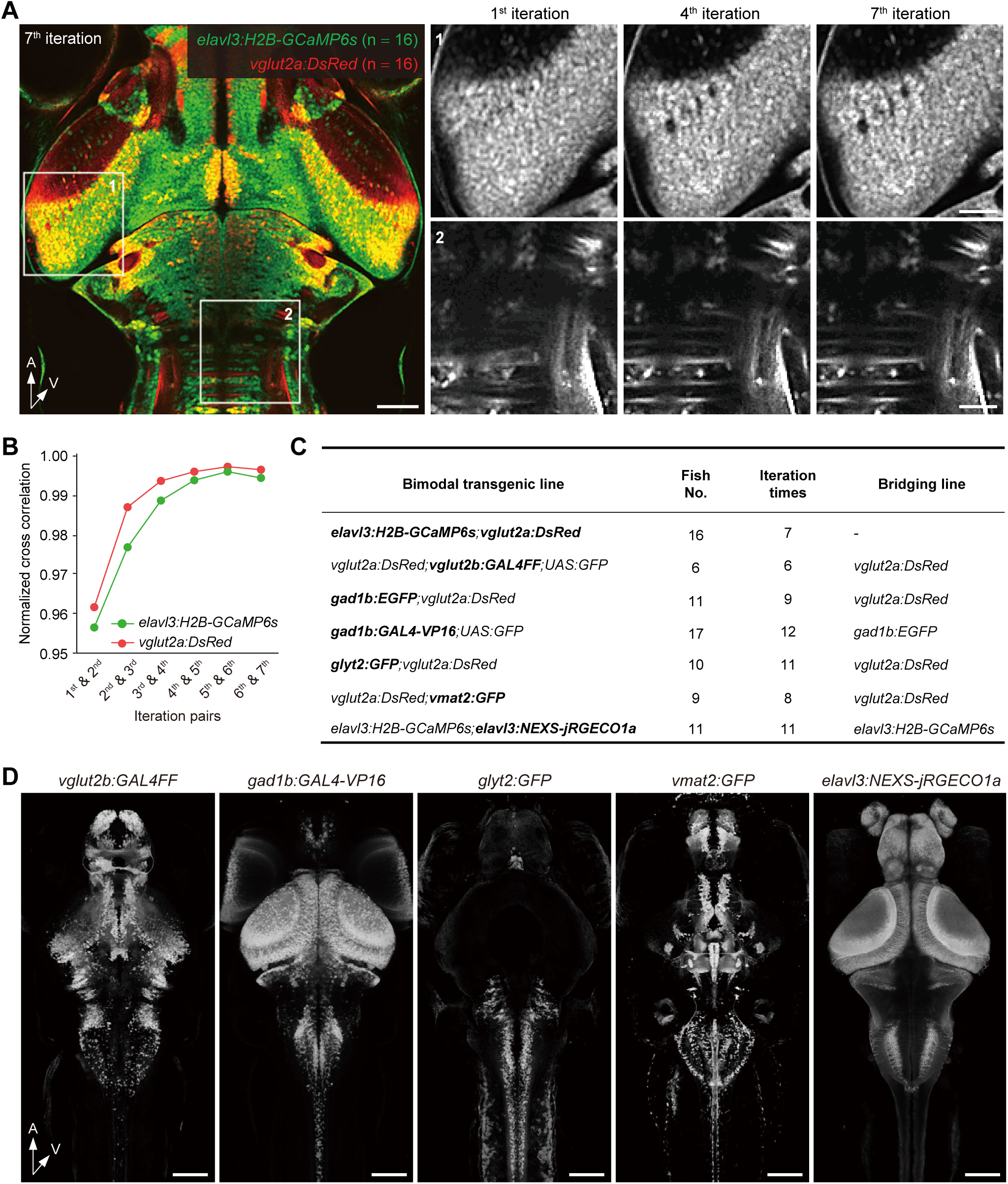
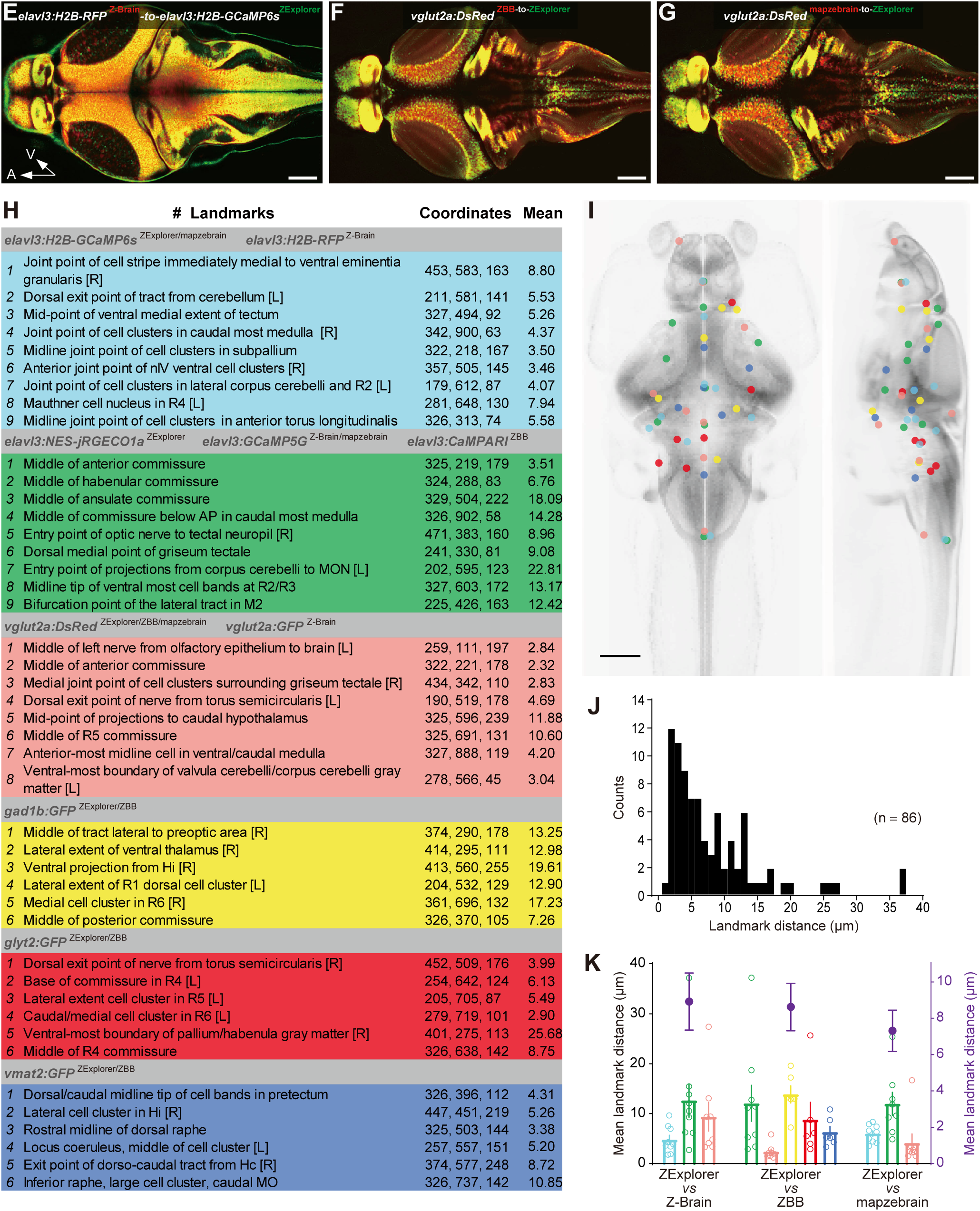

**Figure S3.**
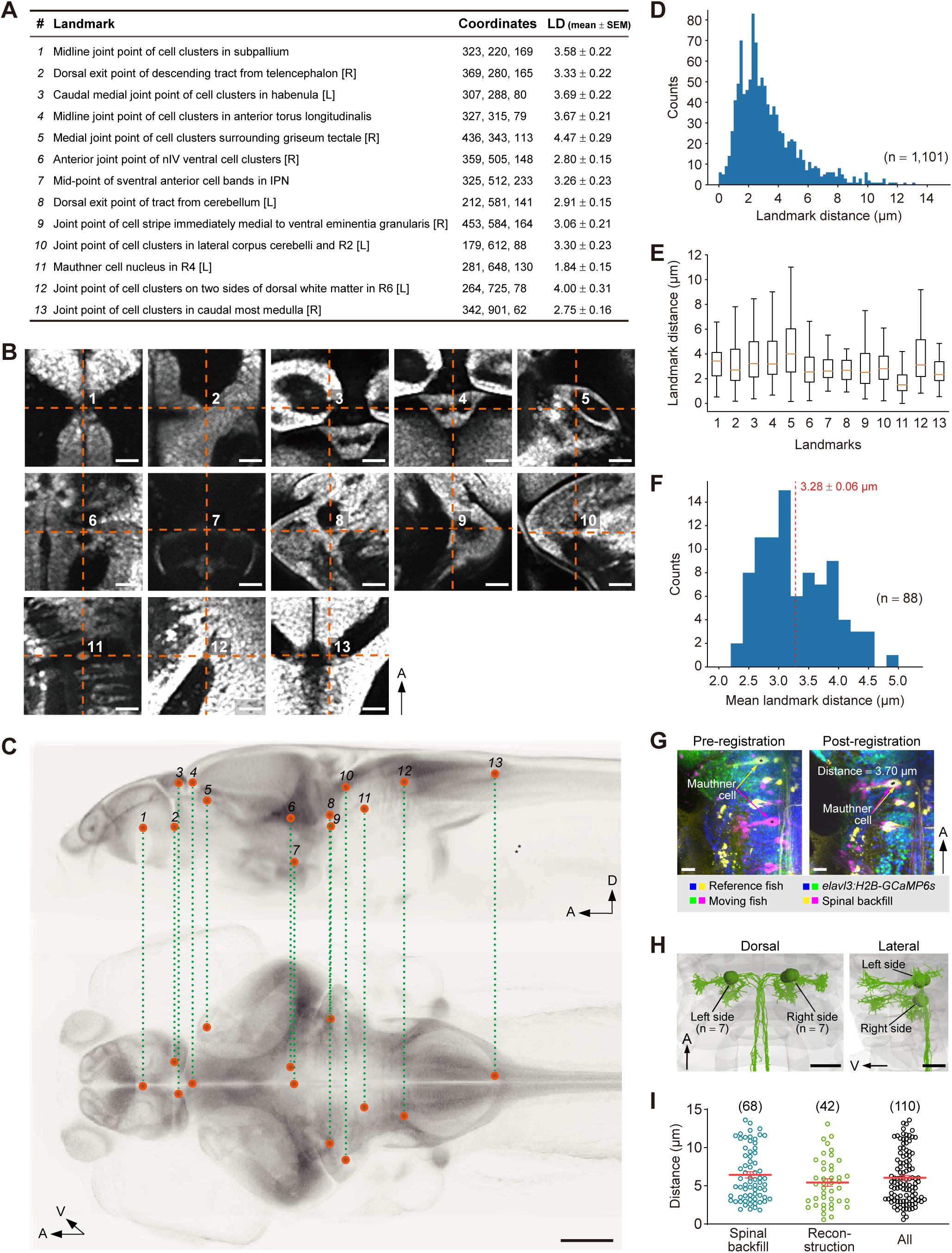
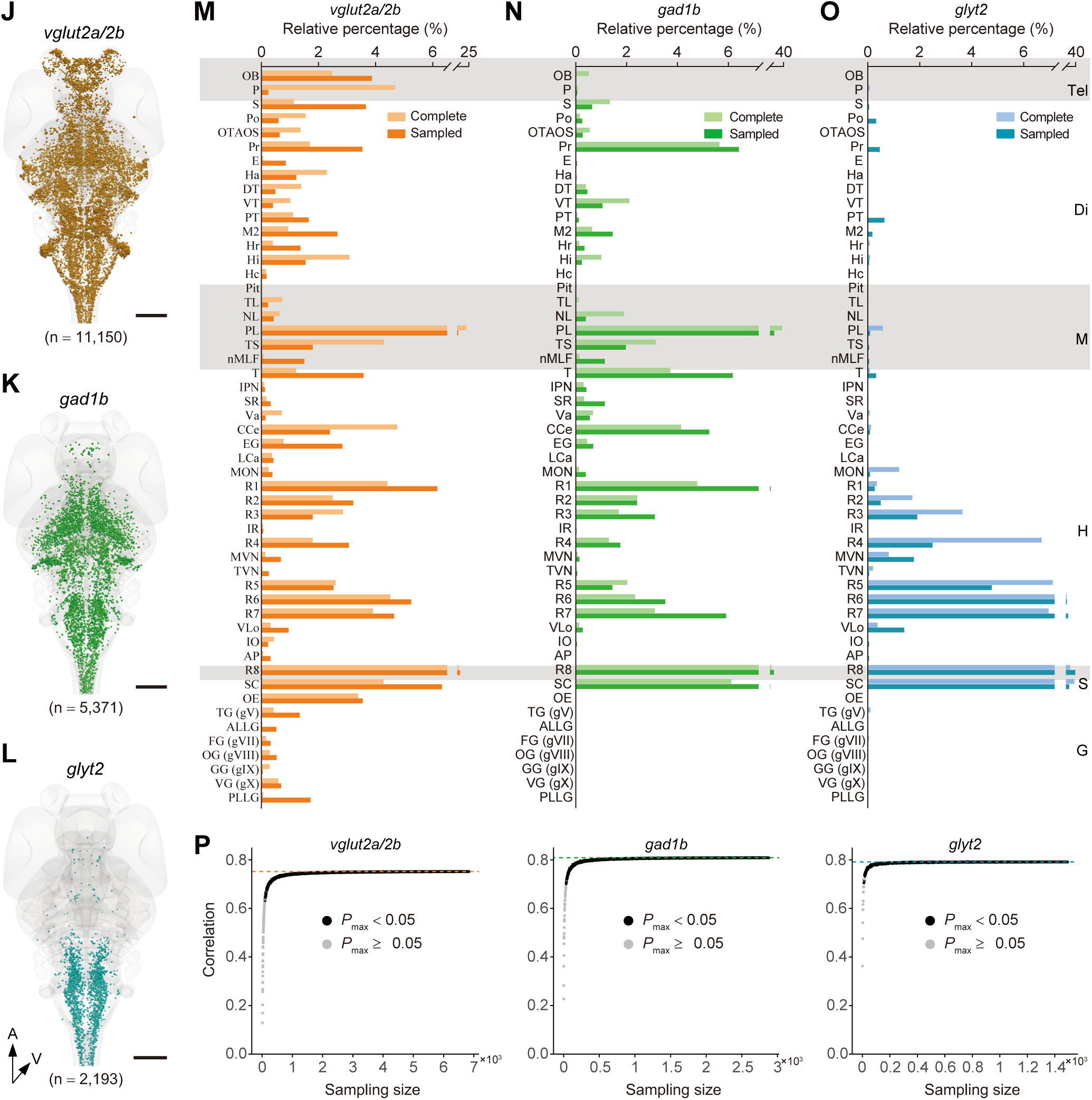

**Figure S4.**
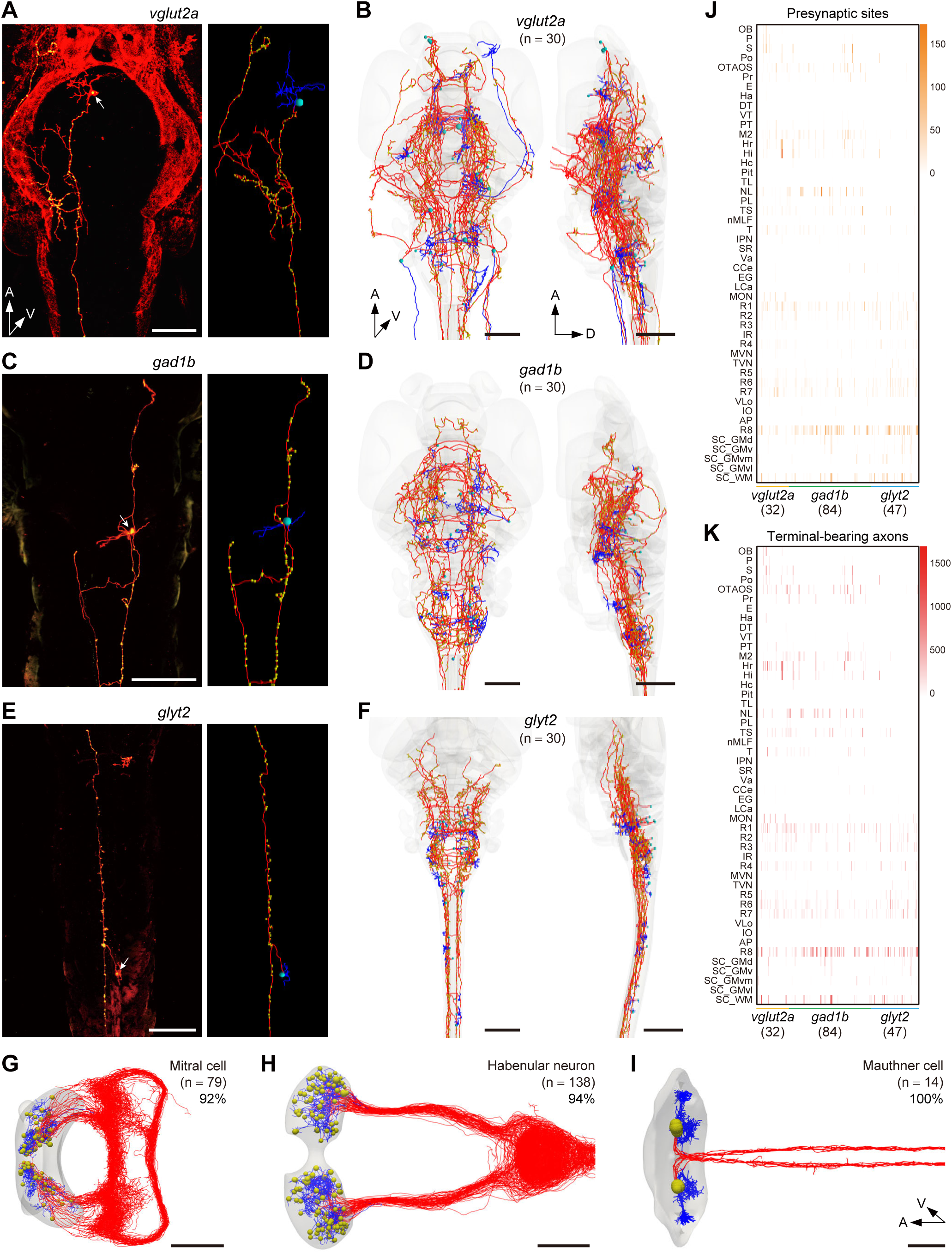

**Figure S5.**
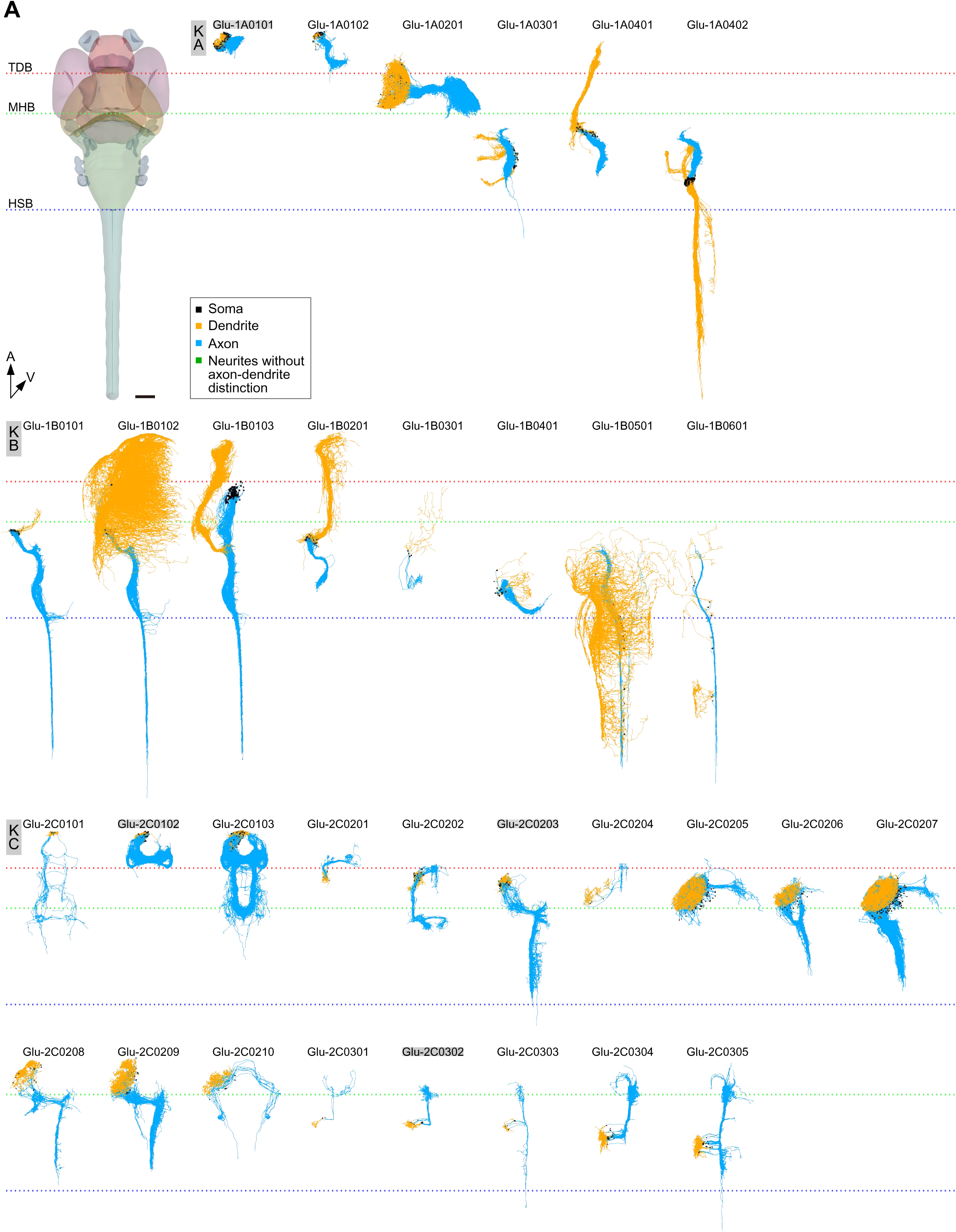
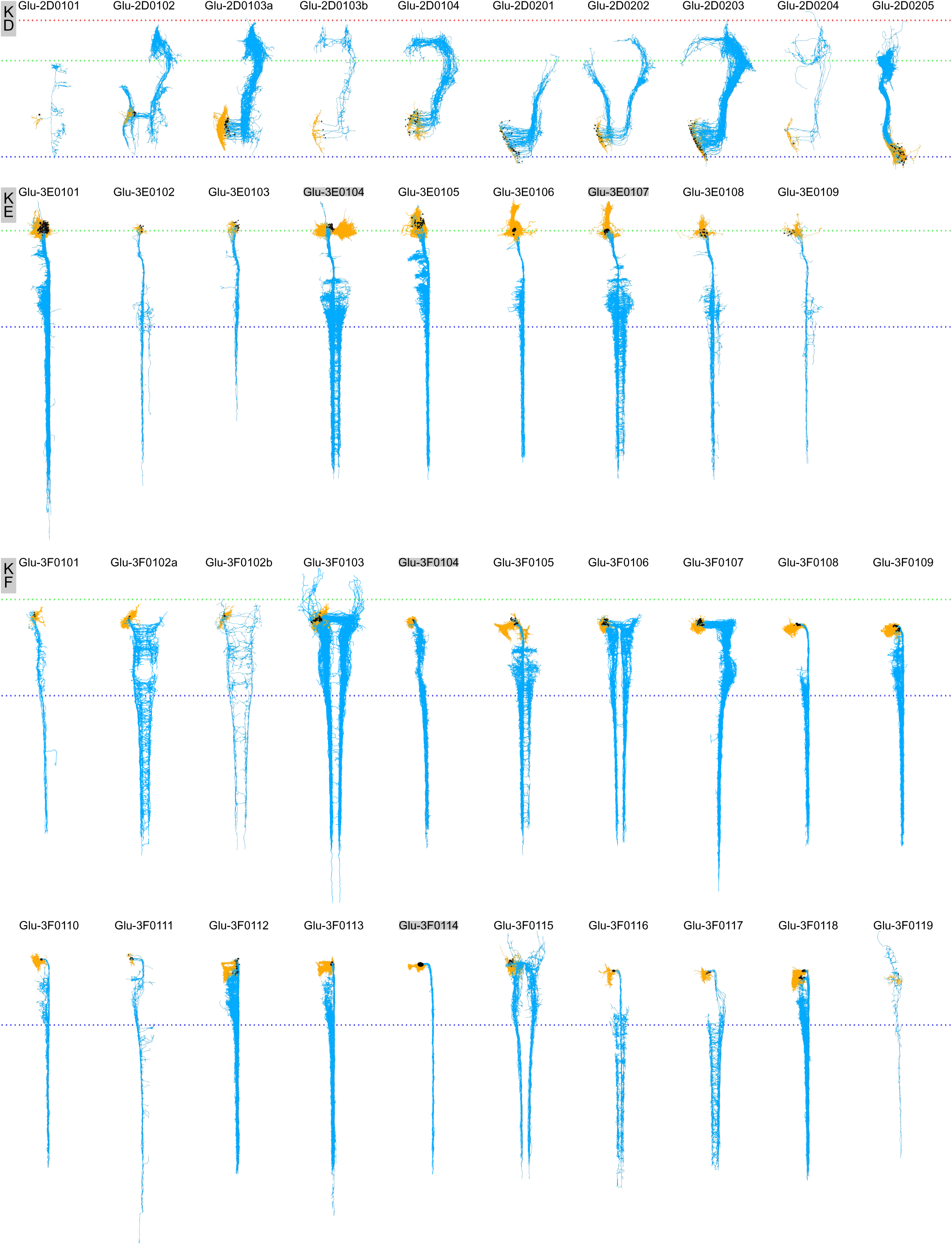
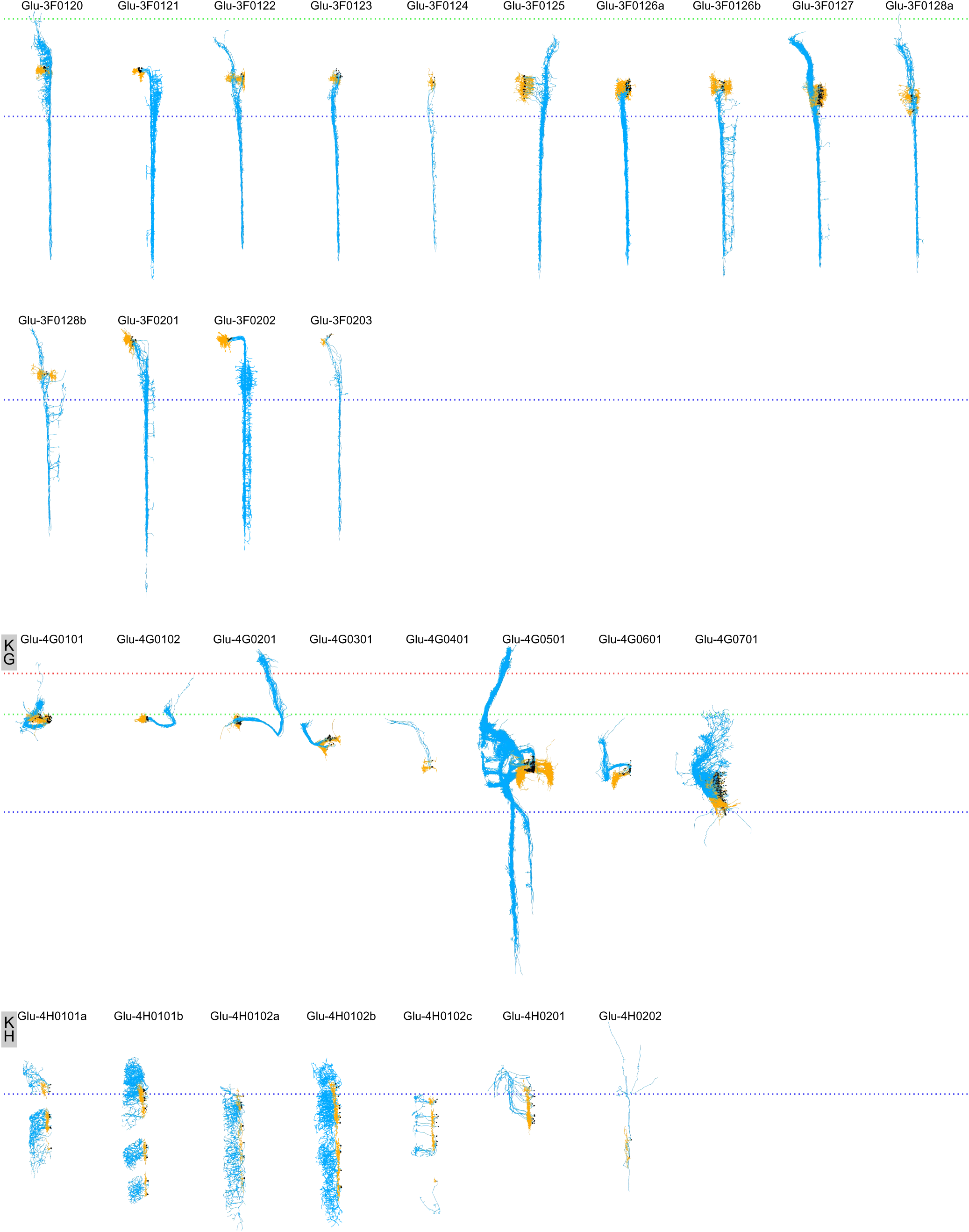
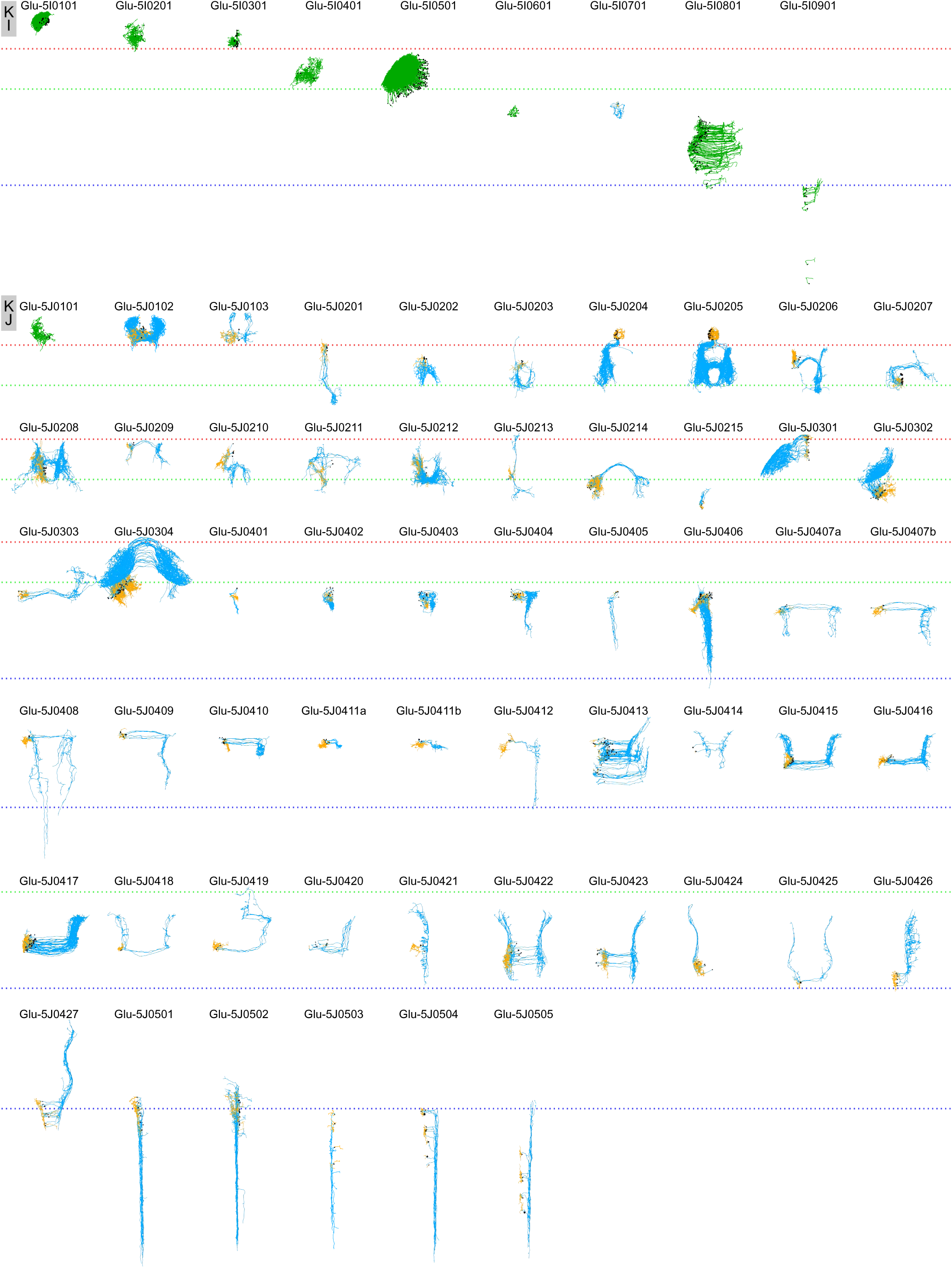
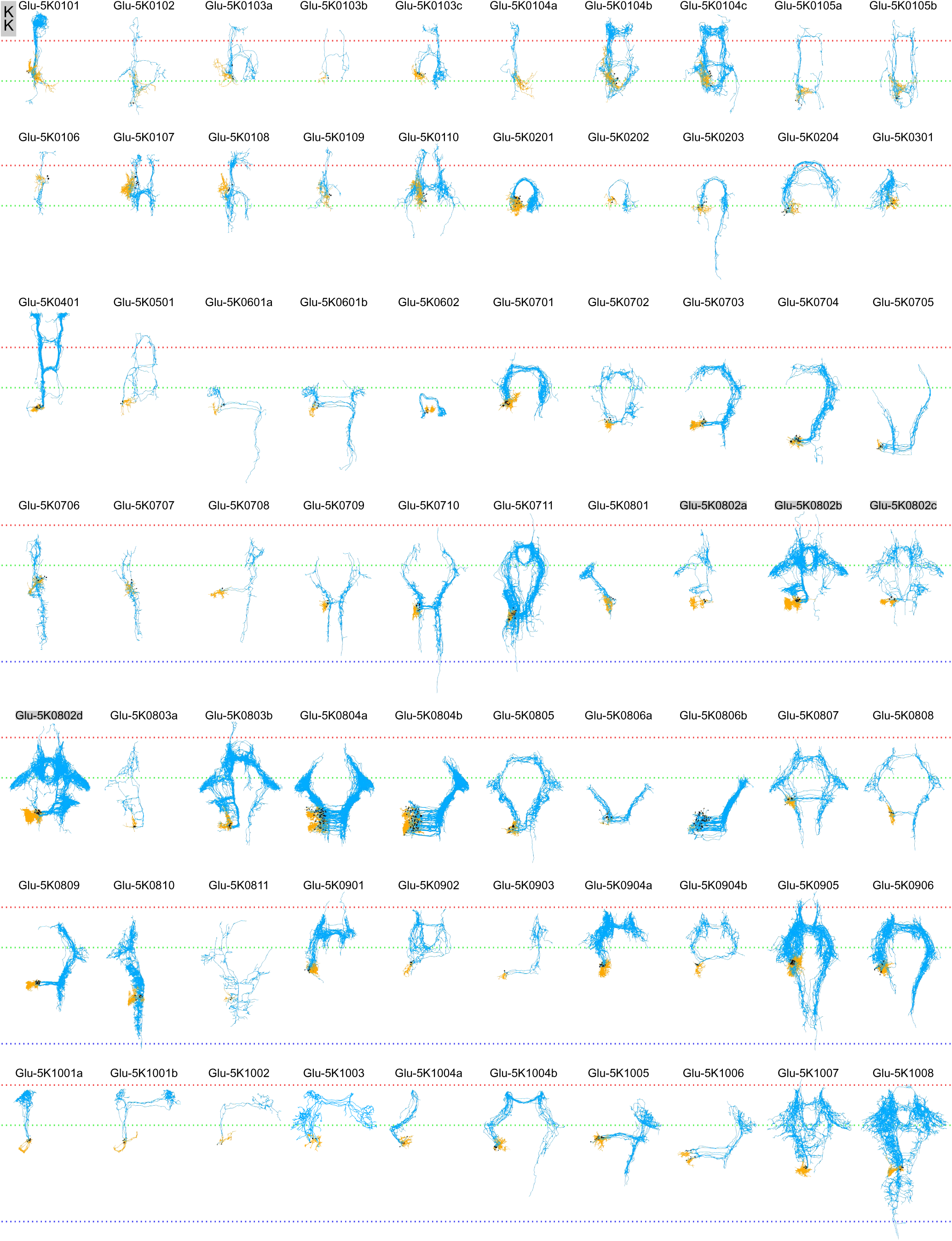
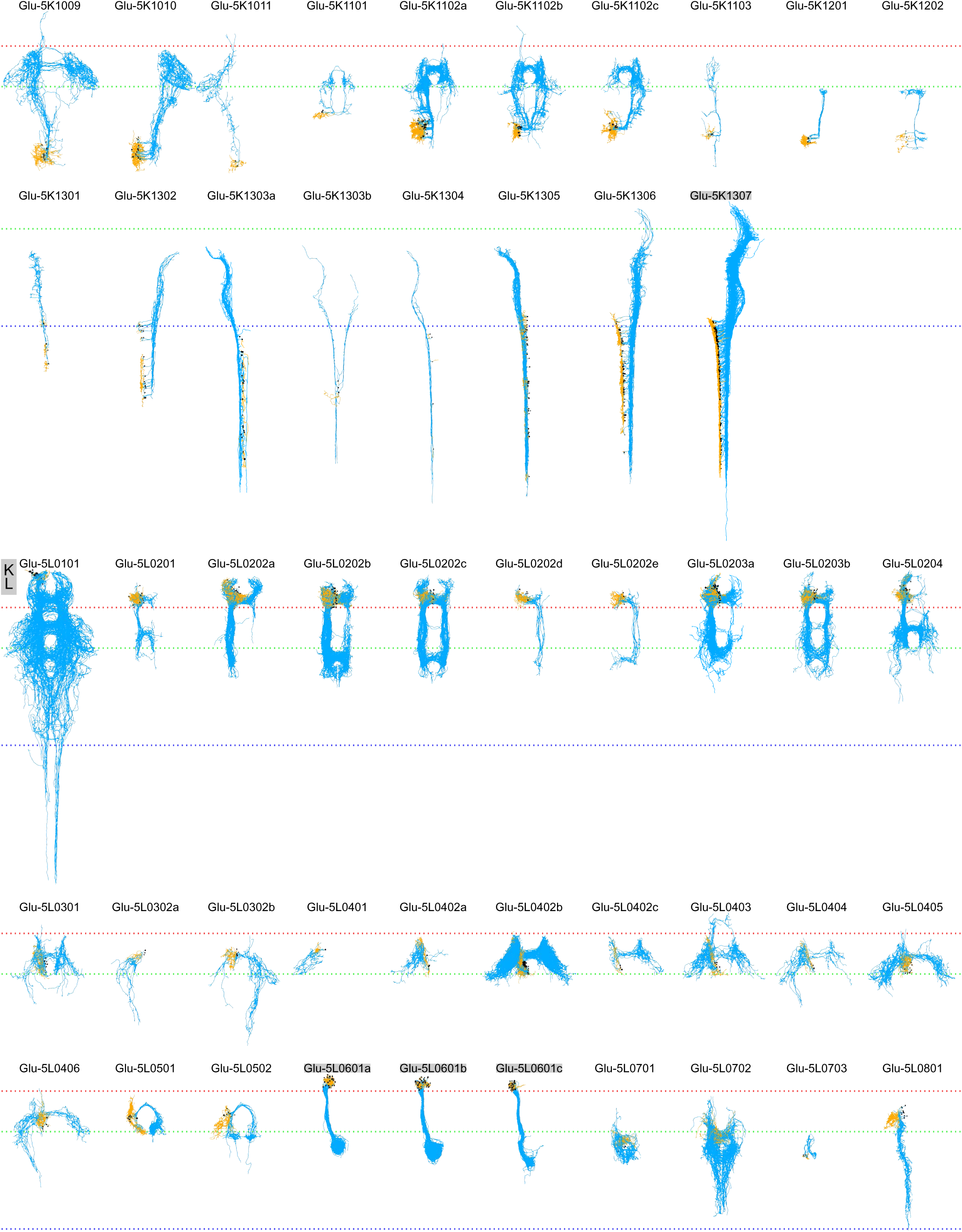
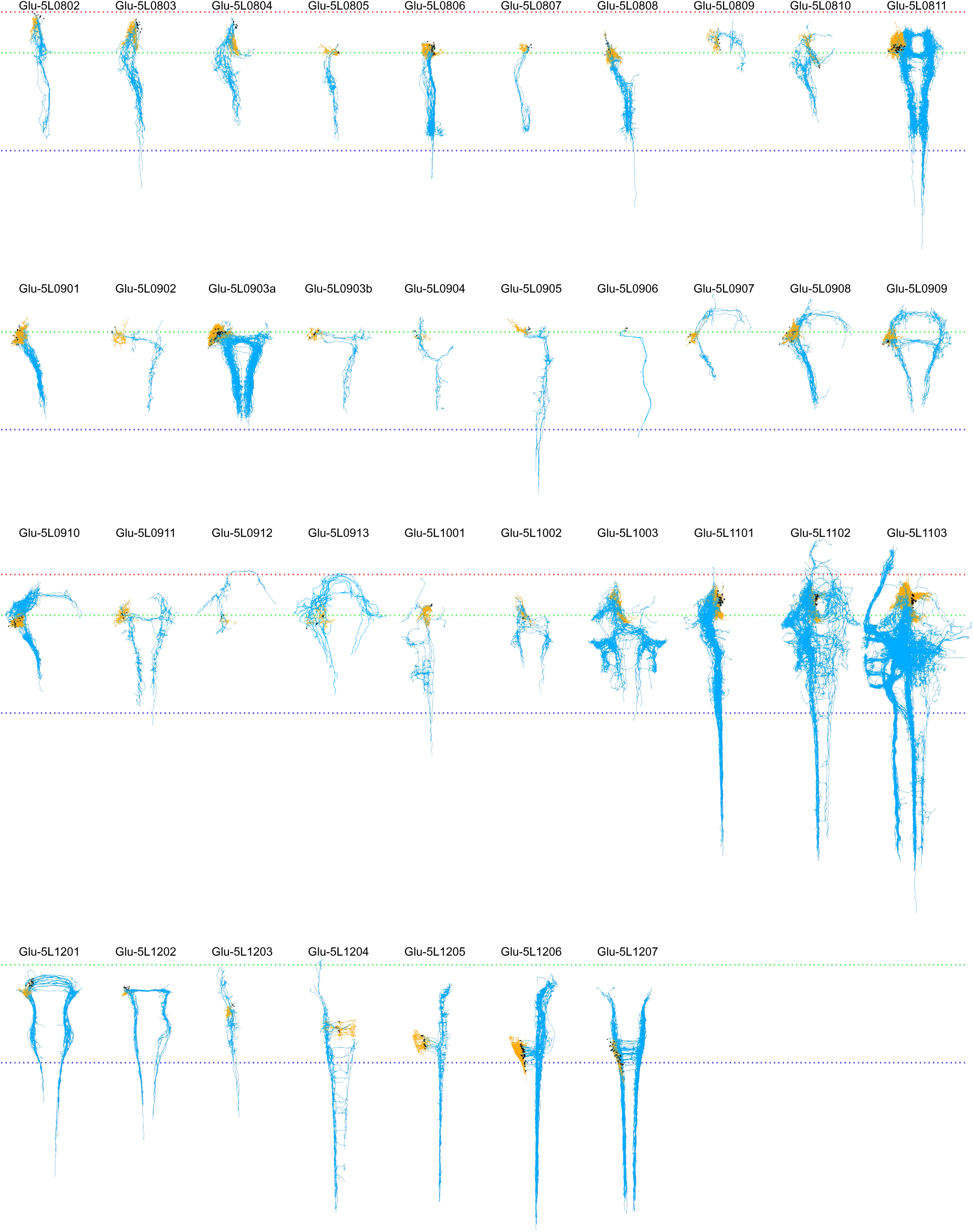
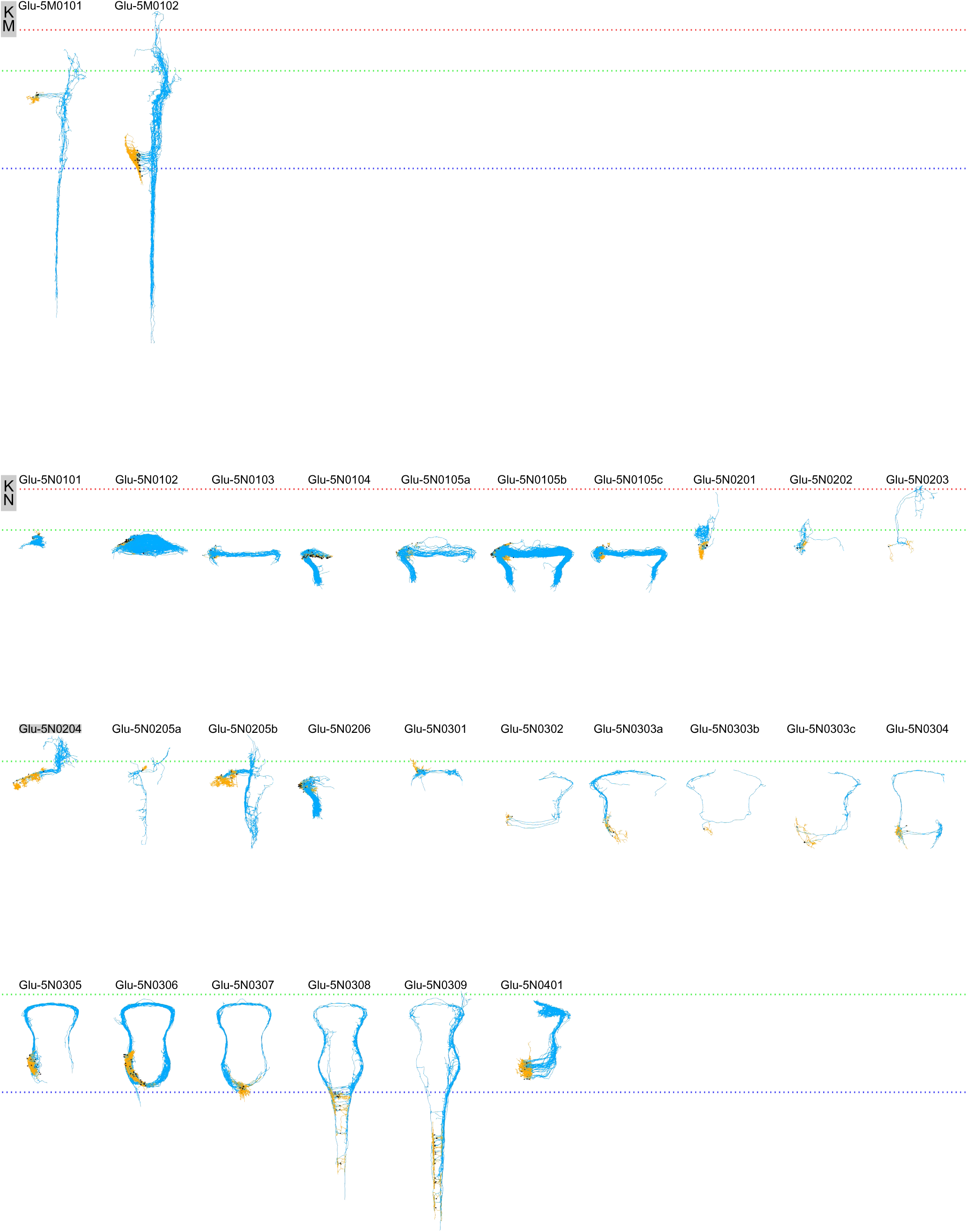
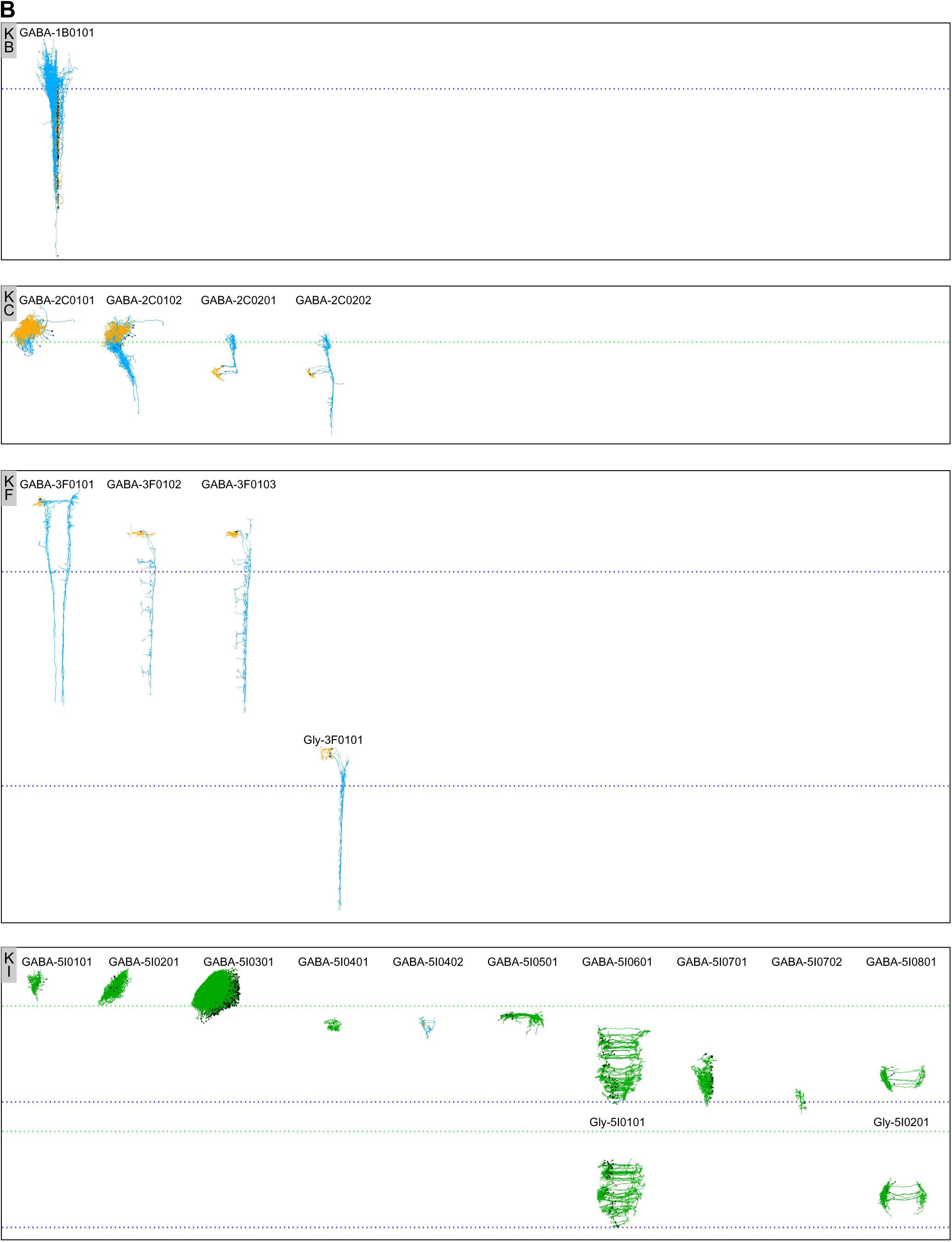
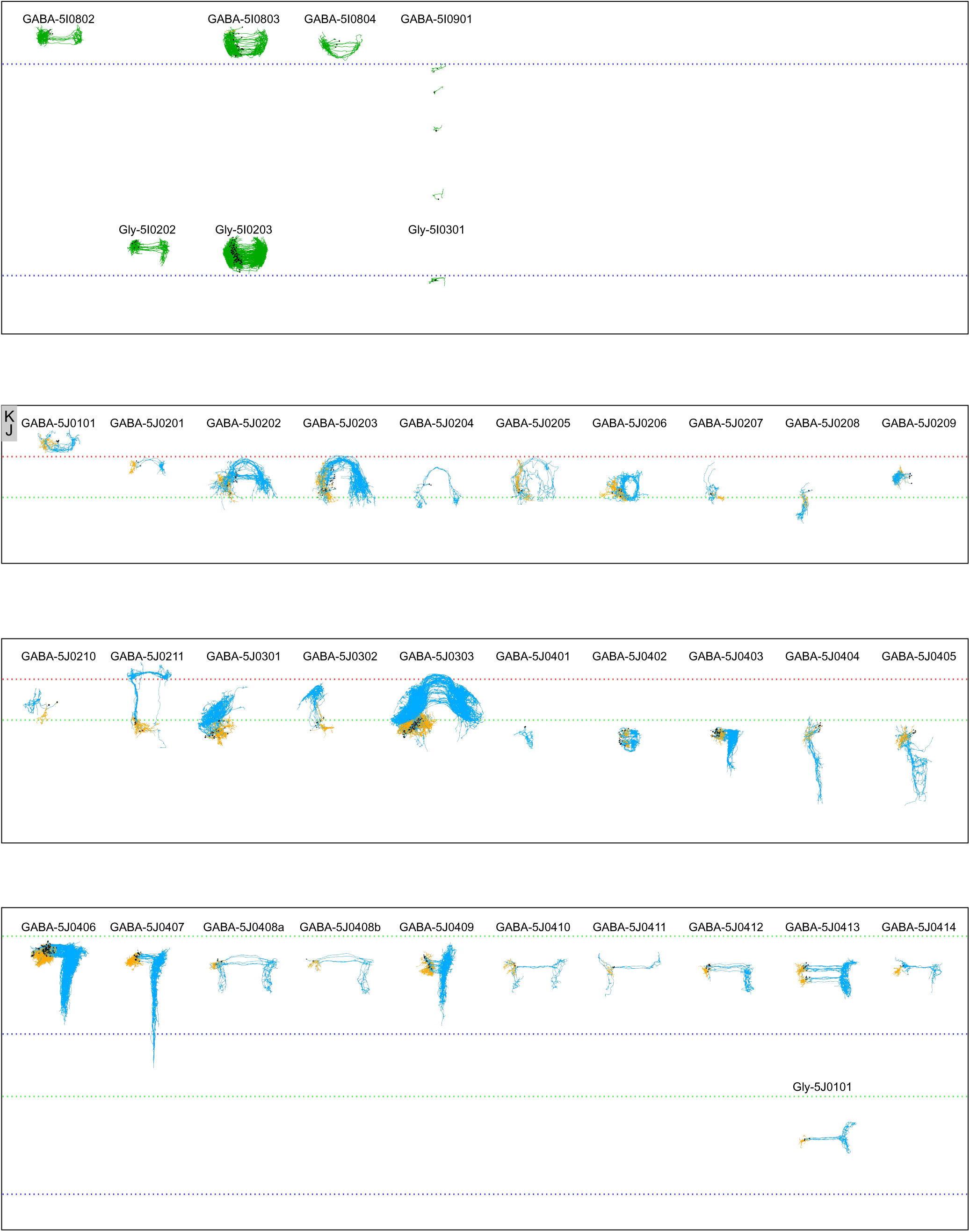
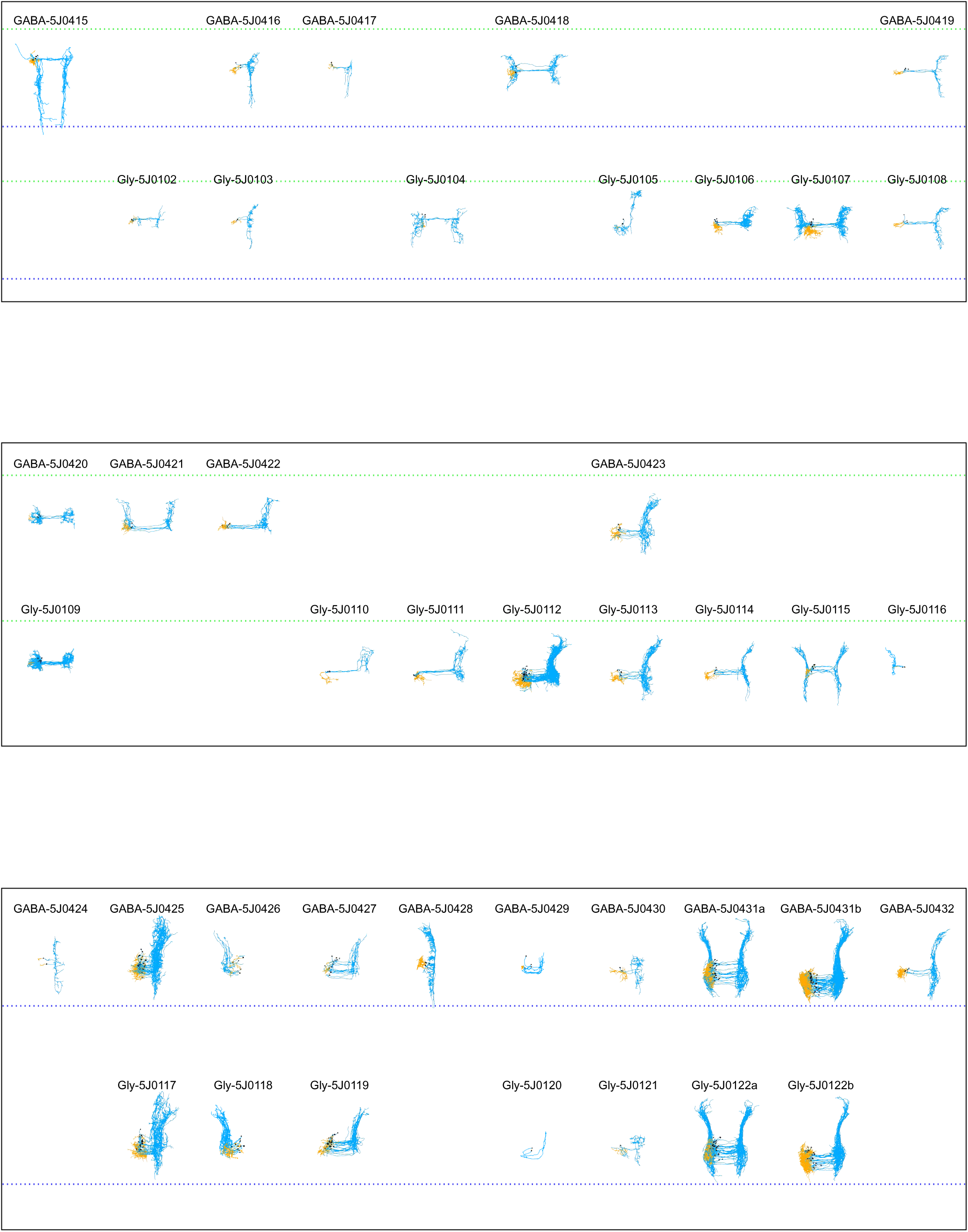
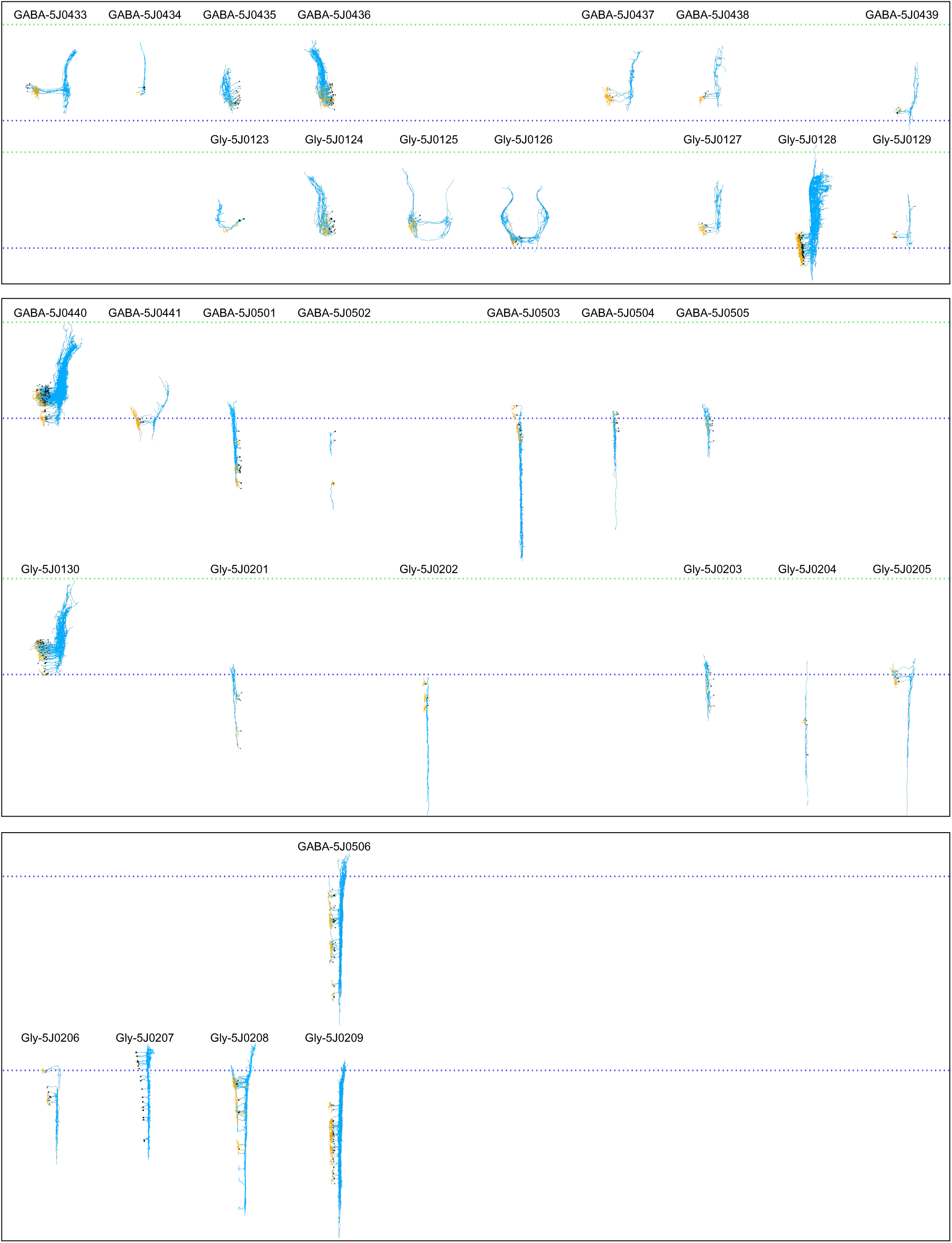
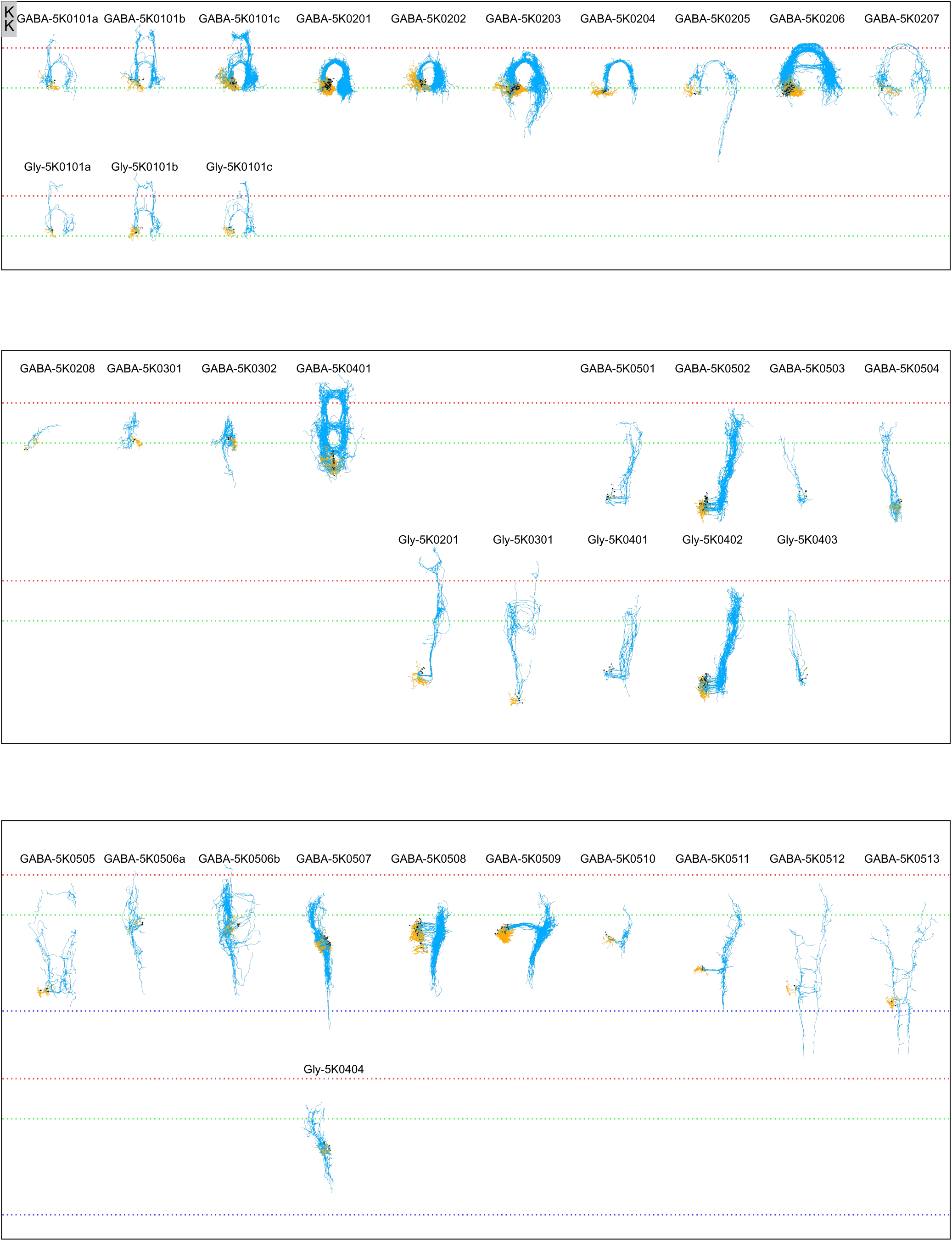
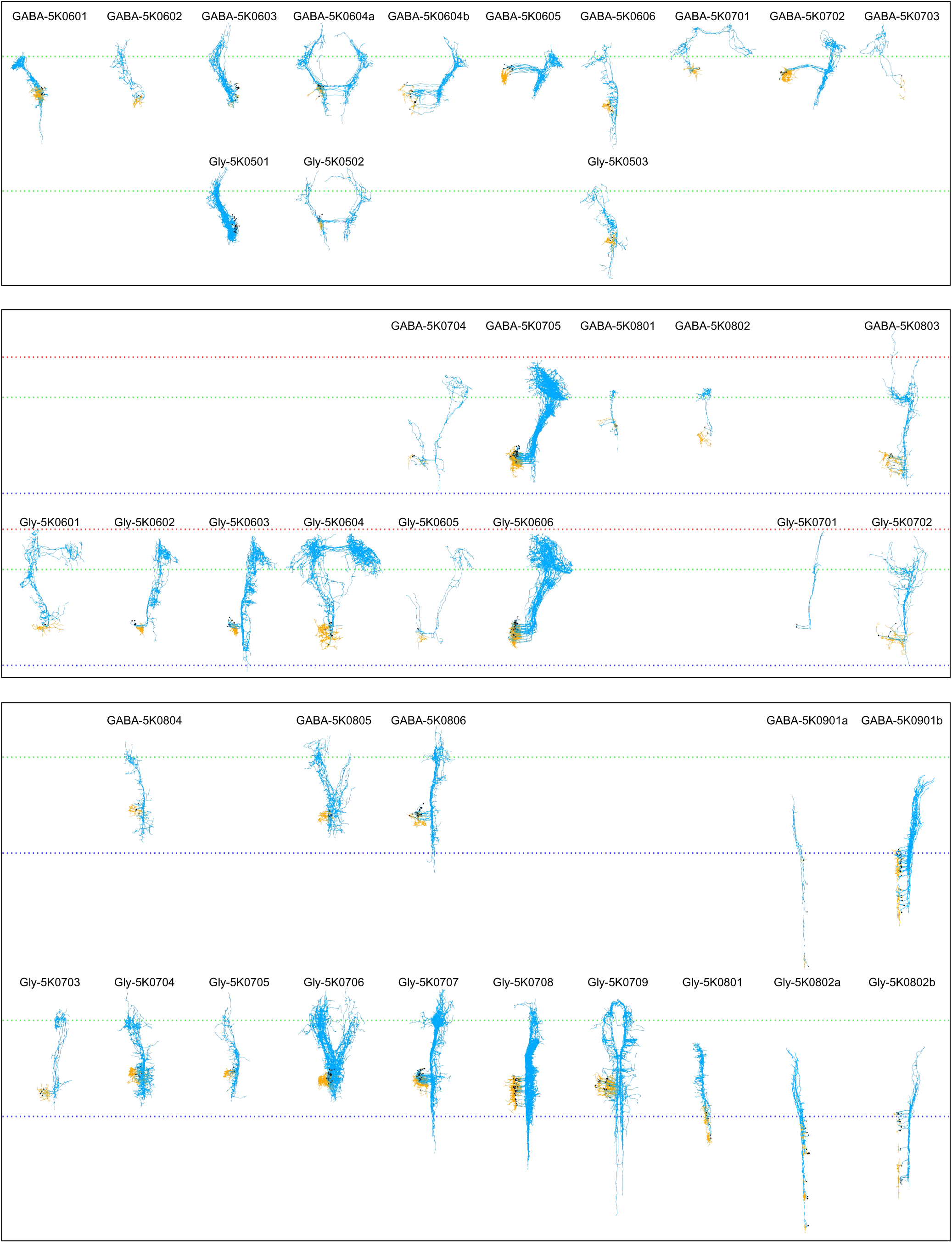
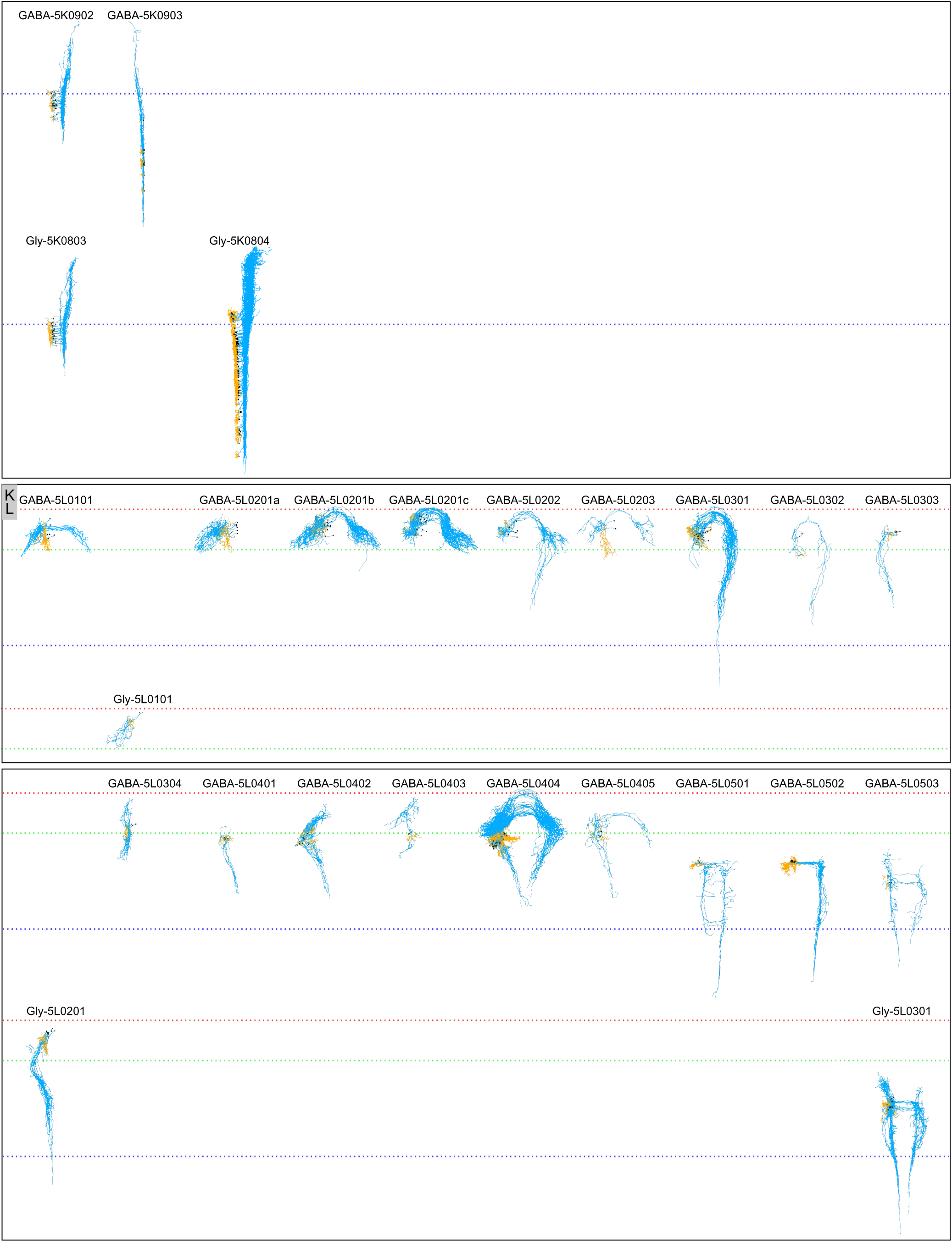
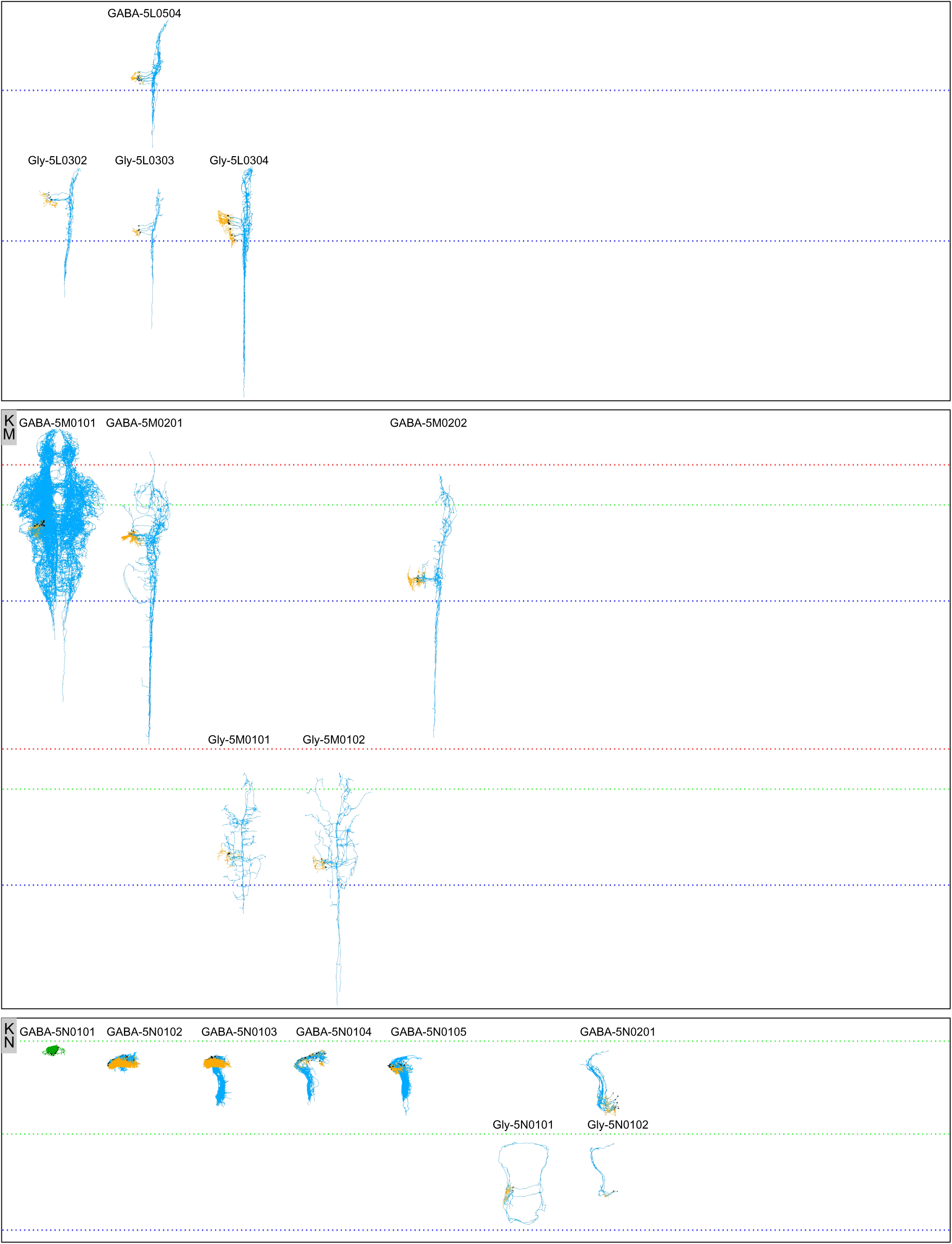

**Figure S6.**
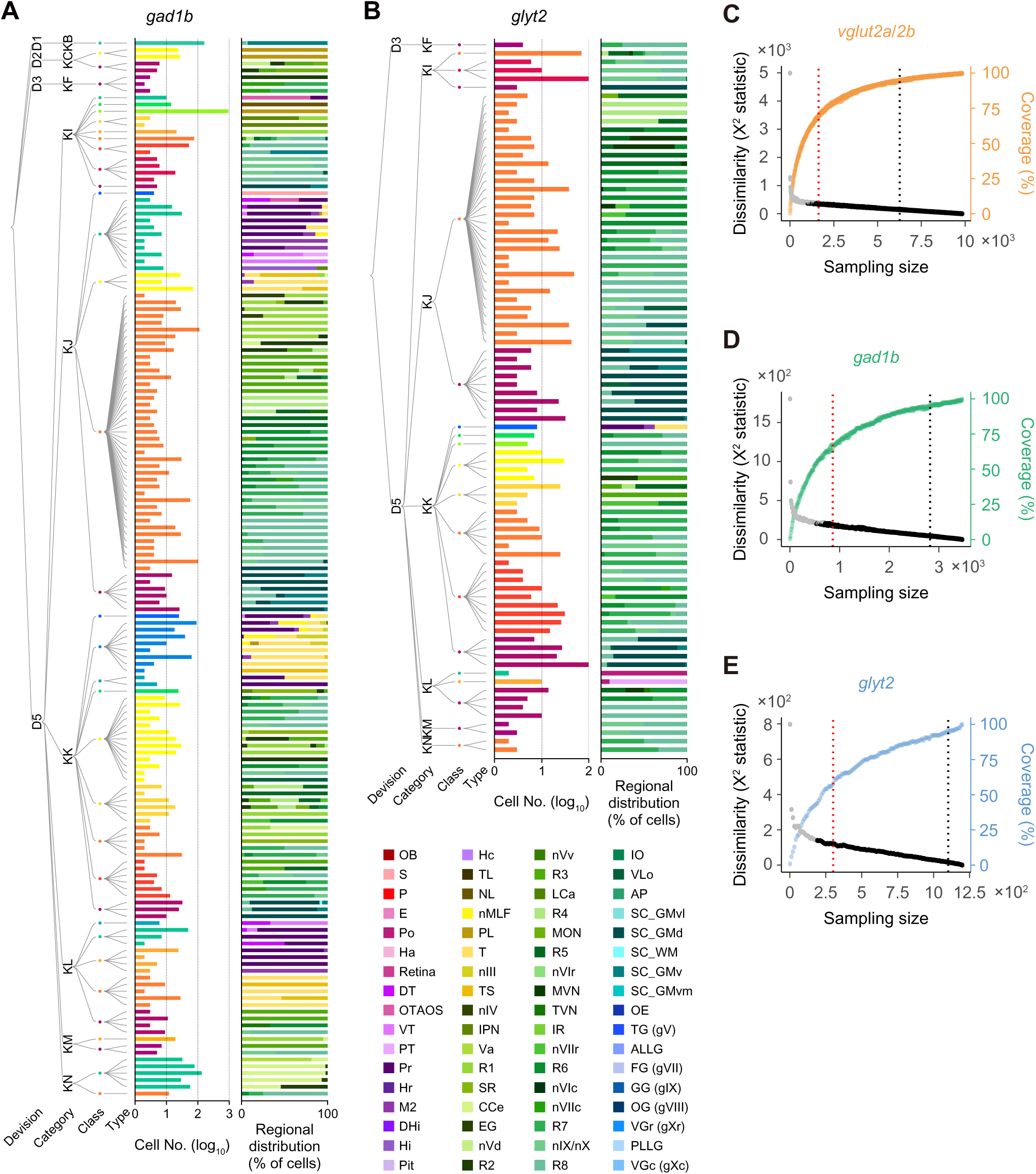

**Figure S7.**
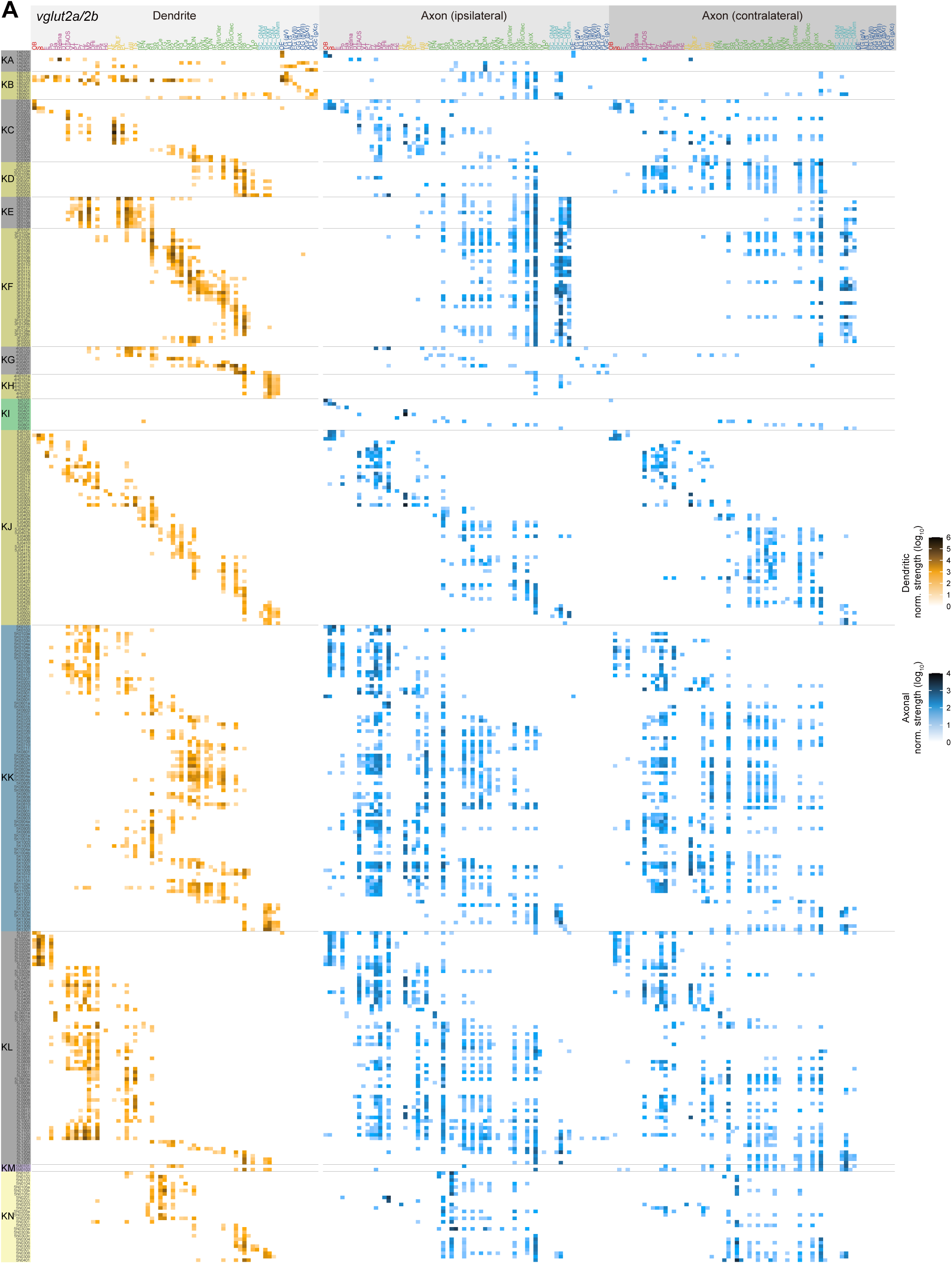
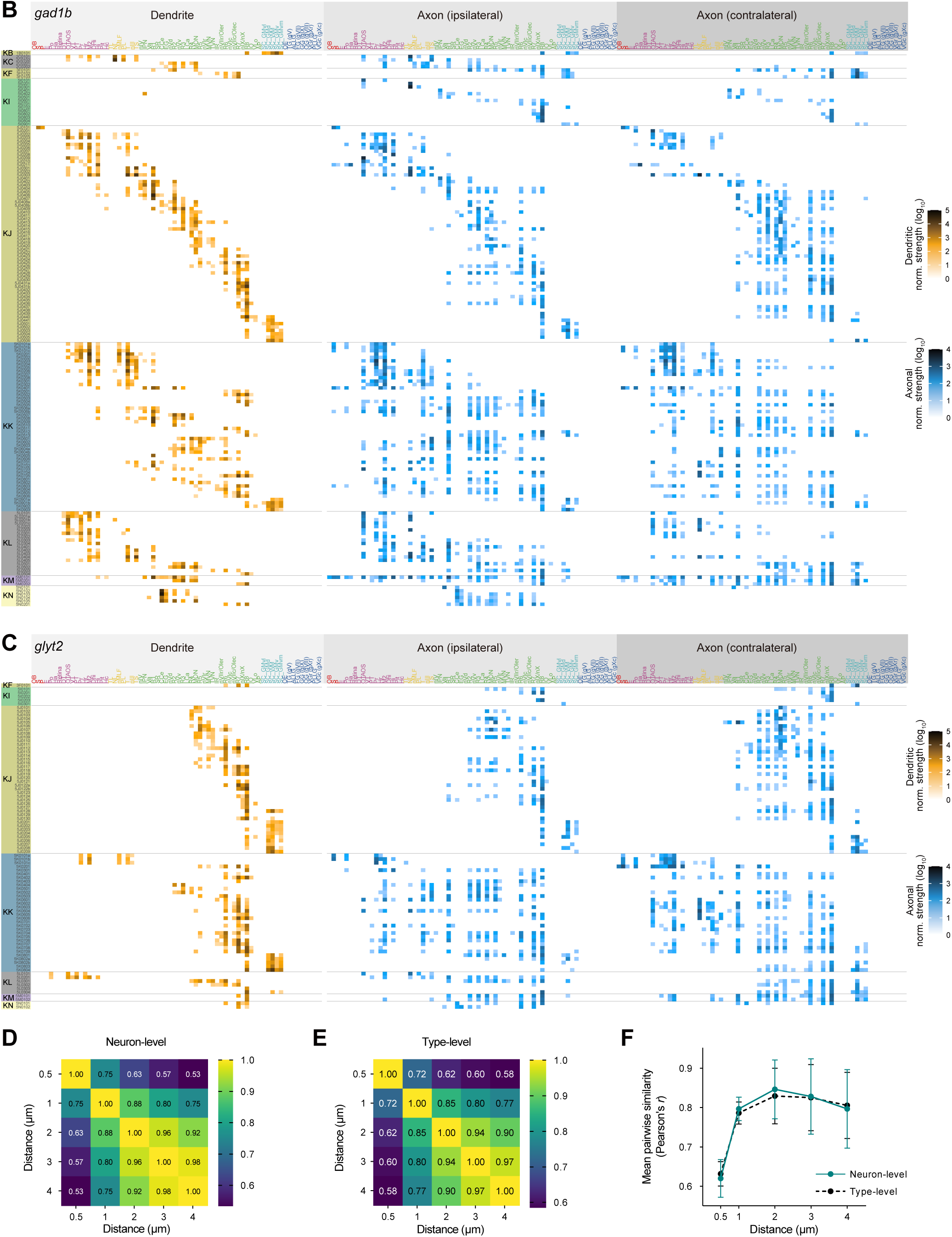
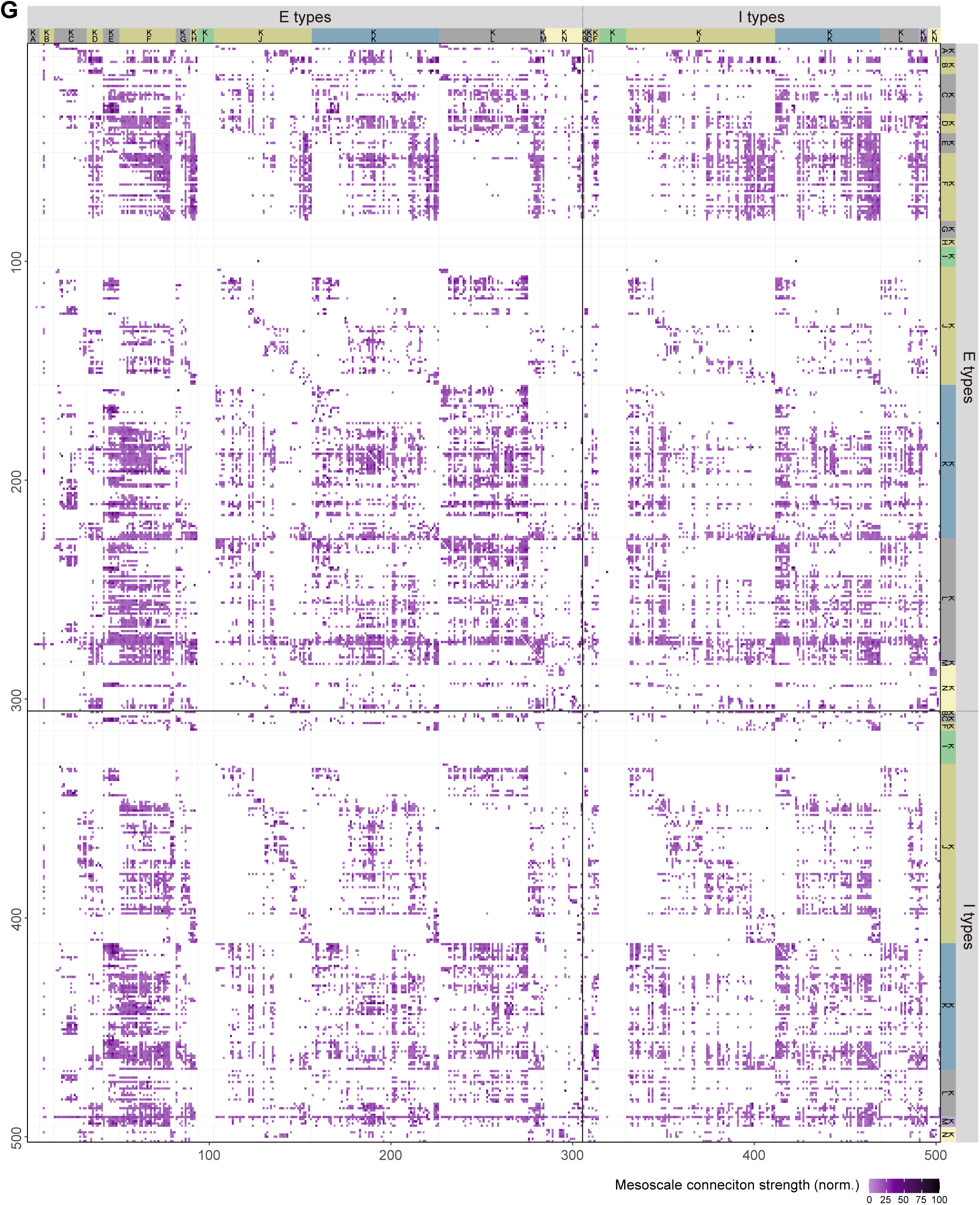

**Figure S8.**
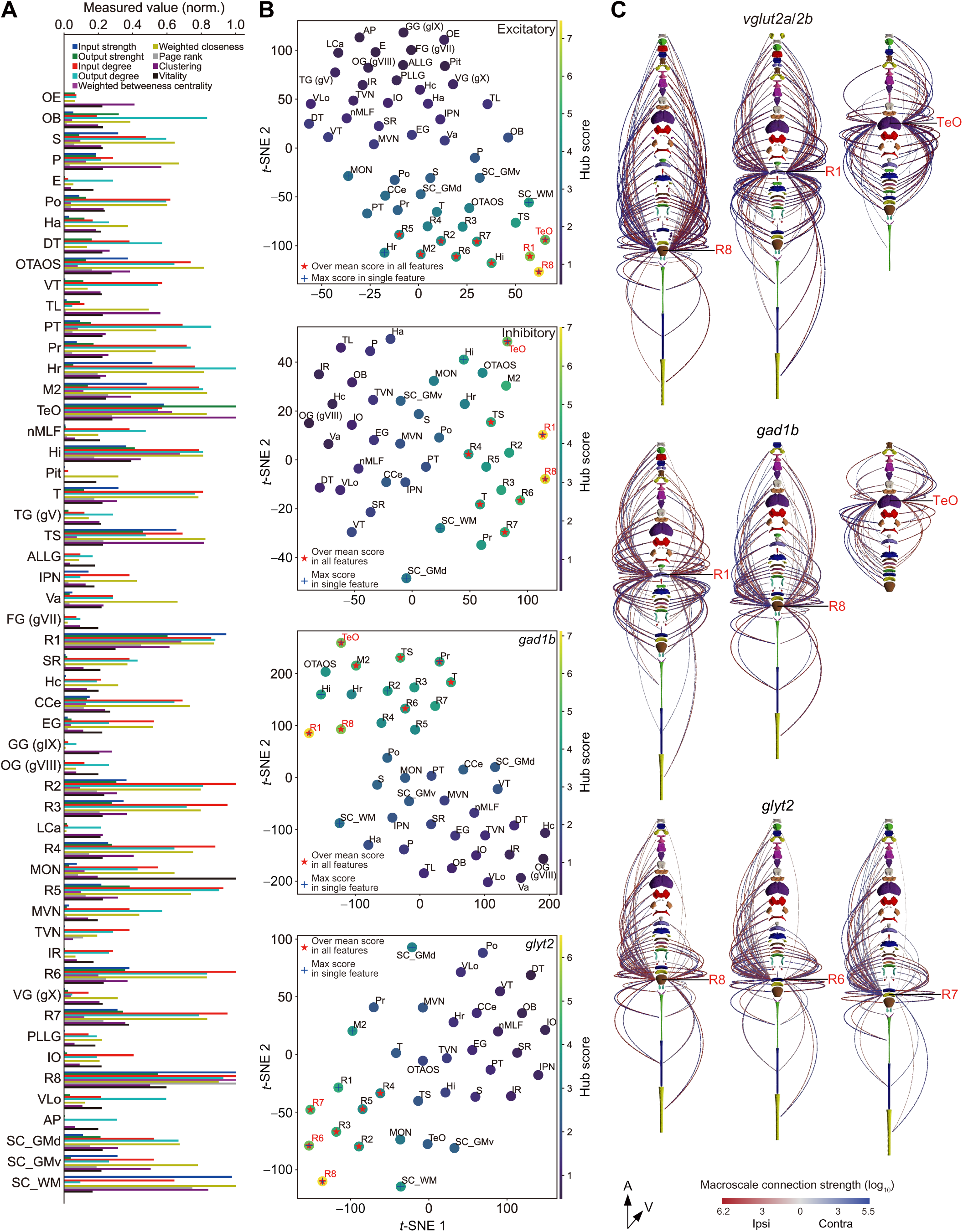
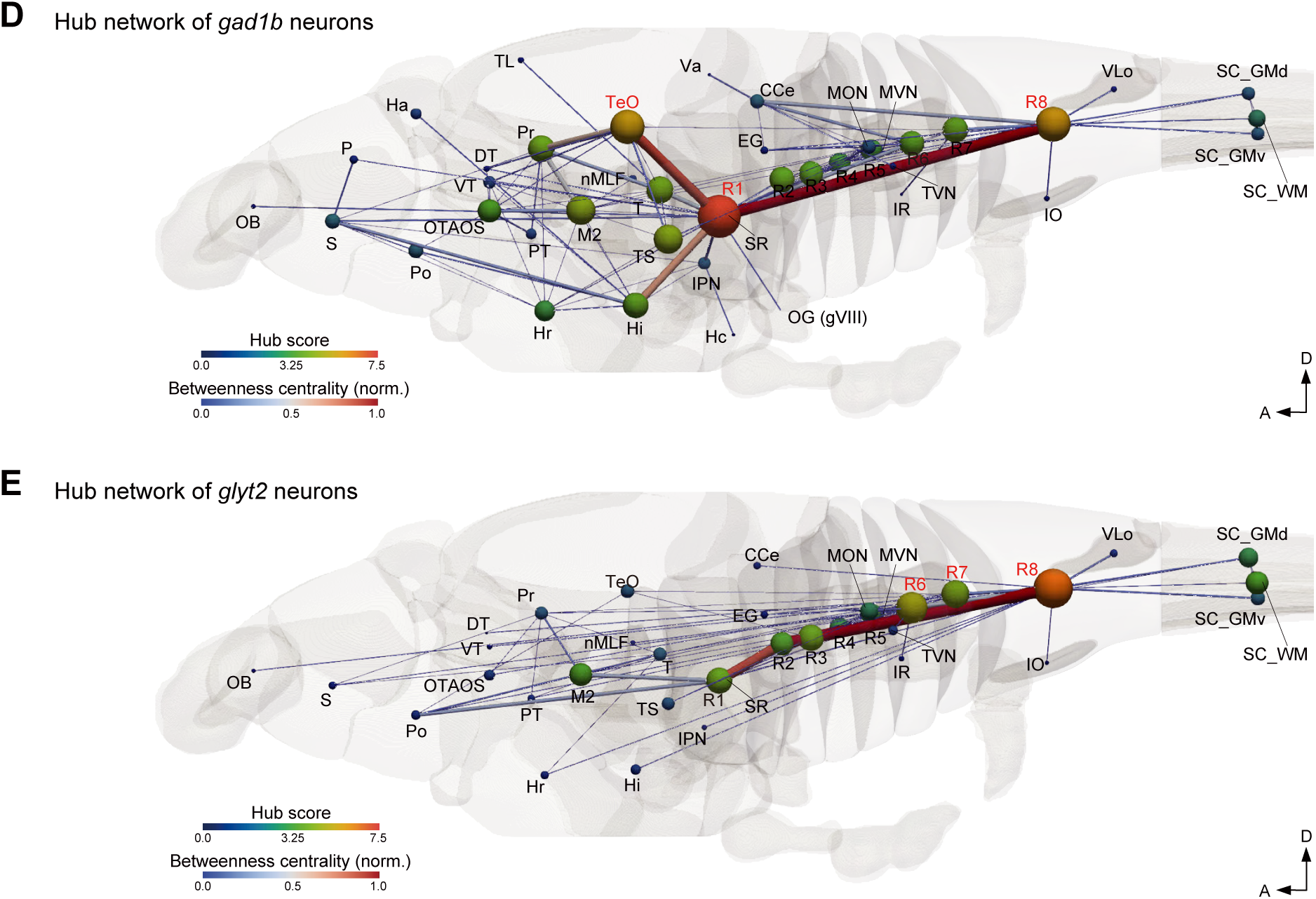

## Notes

https://zebrafish.cn/LM-Atlas/EI

### Competing Interest Statement

The authors have declared no competing interest.

### Summary of Updates

Title and main text revised for clarity; Figure 2, 4, and 6~8 revised; Supplemental data updated; author affiliations updated.

## REFERENCES

1. Santiago, R. y C. Estructura de los centros nerviosos de las aves. Rev. Trim. Histol. Norm. Patol. 1:1–10.

2. Swanson, L.W., and Lichtman, J.W. (2016). From Cajal to Connectome and beyond. Annu. Rev. Neurosci. 39, 197–216. 10.1146/annurev-neuro-071714-033954.

3. Suárez, L.E., Markello, R.D., Betzel, R.F., and Misic, B. (2020). Linking Structure and Function in Macroscale Brain Networks. Trends Cogn. Sci. 24, 302–315. 10.1016/j.tics.2020.01.008.

4. J G White 1, E Southgate, J N Thomson, S.B. (1986). The structure of the nervous system of the nematode Caenorhabditis elegans. Philos. Trans. R. Soc. London. B, Biol. Sci. 314, 1–340. 10.1098/rstb.1986.0056.

5. Cook, S.J., Jarrell, T.A., Brittin, C.A., Wang, Y., Bloniarz, A.E., Yakovlev, M.A., Nguyen, K.C.Q., Tang, L.T.H., Bayer, E.A., Duerr, J.S., et al. (2019). Whole-animal connectomes of both Caenorhabditis elegans sexes. Nature 571, 63–71. 10.1038/s41586-019-1352-7.

6. Witvliet, D., Mulcahy, B., Mitchell, J.K., Meirovitch, Y., Berger, D.R., Wu, Y., Liu, Y., Koh, W.X., Parvathala, R., Holmyard, D., et al. (2021). Connectomes across development reveal principles of brain maturation. Nature 596, 257–261. 10.1038/s41586-021-03778-8.

7. Chiang, A.S., Lin, C.Y., Chuang, C.C., Chang, H.M., Hsieh, C.H., Yeh, C.W., Shih, C.T., Wu, J.J., Wang, G.T., Chen, Y.C., et al. (2011). Three-dimensional reconstruction of brain-wide wiring networks in drosophila at single-cell resolution. Curr. Biol. 21, 1–11. 10.1016/j.cub.2010.11.056.

8. Zheng, Z., Lauritzen, J.S., Perlman, E., Saalfeld, S., Fetter, R.D., Bock, D.D., Zheng, Z., Lauritzen, J.S., Perlman, E., Robinson, C.G., et al. (2018). Resource A Complete Electron Microscopy Volume of the Brain of Adult Drosophila melanogaster A Complete Electron Microscopy Volume of the Brain of Adult Drosophila melanogaster. Cell 174, 730–743.e22. 10.1016/j.cell.2018.06.019.

9. Scheffer, L.K., Xu, C.S., Januszewski, M., Lu, Z., Takemura, S., Hayworth, K.J., Huang, G.B., Shinomiya, K., Maitlin-Shepard, J., Berg, S., et al. (2020). A connectome and analysis of the adult Drosophila central brain. Elife 9, 1–74. 10.7554/eLife.57443.

10. Winding, M., Pedigo, B.D., Barnes, C.L., Patsolic, H.G., Park, Y., Kazimiers, T., Fushiki, A., Andrade, I. V., Khandelwal, A., Valdes-Aleman, J., et al. (2023). The connectome of an insect brain. Science 379, eadd9330. 10.1126/science.add9330.

11. Dorkenwald, S., Matsliah, A., Sterling, A.R., Schlegel, P., Yu, S.-C., McKellar, C.E., Lin, A., Costa, M., Eichler, K., Yin, Y., et al. (2024). Neuronal wiring diagram of an adult brain. Nature 634, 124–138. 10.1038/s41586-024-07558-y.

12. Winnubst, J., Bas, E., Ferreira, T.A., Wu, Z., Economo, M.N., Edson, P., Arthur, B.J., Bruns, C., Rokicki, K., Schauder, D., et al. (2019). Reconstruction of 1,000 Projection Neurons Reveals New Cell Types and Organization of Long-Range Connectivity in the Mouse Brain. Cell 179, 268–281.e13. 10.1016/j.cell.2019.07.042.

13. Peng, H., Xie, P., Liu, L., Kuang, X., Wang, Y., Qu, L., Gong, H., Jiang, S., Li, A., Ruan, Z., et al. (2021). Morphological diversity of single neurons in molecularly defined cell types. Nature 598, 174–181. 10.1038/s41586-021-03941-1.

14. Gao, L., Wang, H., Chen, Y., Xu, C., Wang, X., and Yan, J. (2026). Integrative analysis of single-neuron projectomes links connectome, transcriptome, and function in the mouse cortex. Neuron, 1–19. 10.1016/j.neuron.2025.10.019.

15. Gao, L., Liu, S., Gou, L., Hu, Y., Liu, Y., Deng, L., Ma, D., Wang, H., Yang, Q., Chen, Z., et al. (2022). Single-neuron projectome of mouse prefrontal cortex. Nat. Neurosci. 25, 515–529. 10.1038/s41593-022-01041-5.

16. Qiu, S., Hu, Y., Huang, Y., Gao, T., Wang, X., Wang, D., Ren, B., Shi, X., Chen, Y., Wang, X., et al. (2024). Whole-brain spatial organization of hippocampal single-neuron projectomes. Science. 383. 10.1126/science.adj9198.

17. Zeng, H. (2018). Mesoscale connectomics. Curr. Opin. Neurobiol. 50, 154–162. 10.1016/j.conb.2018.03.003.

18. Yao, S., Wang, Q., Hirokawa, K.E., Ouellette, B., Ahmed, R., Bomben, J., Brouner, K., Casal, L., Caldejon, S., Cho, A., et al. (2023). A whole-brain monosynaptic input connectome to neuron classes in mouse visual cortex. Nat. Neurosci. 26, 350–364. 10.1038/s41593-022-01219-x.

19. Kasthuri, N., Hayworth, K.J., Berger, D.R., Schalek, R.L., Conchello, J.A., Knowles-Barley, S., Lee, D., Vázquez-Reina, A., Kaynig, V., Jones, T.R., et al. (2015). Saturated Reconstruction of a Volume of Neocortex. Cell 162, 648–661. 10.1016/j.cell.2015.06.054.

20. Motta, A., Berning, M., Boergens, K.M., Staffler, B., Beining, M., Loomba, S., Hennig, P., Wissler, H., and Helmstaedter, M. (2019). Dense connectomic reconstruction in layer 4 of the somatosensory cortex. Science. 366. 10.1126/science.aay3134.

21. Ngai, J. (2022). BRAIN 2.0: Transforming neuroscience. Cell 185, 4–8. 10.1016/j.cell.2021.11.037.

22. Mueller, D.T., and Wullimann, M. (2016). Atlas of Early Zebrafish Brain Development (Elsevier) 10.1016/C2013-0-00639-1.

23. Tabor, K.M., Marquart, G.D., Hurt, C., Smith, T.S., Geoca, A.K., Bhandiwad, A.A., Subedi, A., Sinclair, J.L., Rose, H.M., Polys, N.F., et al. (2019). Brain-wide cellular resolution imaging of cre transgenic zebrafish lines for functional circuit-mapping. Elife 8, 1–21. 10.7554/eLife.42687.

24. Hiraki-Kajiyama, T., Miyasaka, N., Ando, R., Wakisaka, N., Itoga, H., Onami, S., and Yoshihara, Y. (2024). An atlas and database of neuropeptide gene expression in the adult zebrafish forebrain. J. Comp. Neurol. 532, 1–46. 10.1002/cne.25619.

25. Cui, W.W., Low, S.E., Hirata, H., Saint-Amant, L., Geisler, R., Hume, R.I., and Kuwada, J.Y. (2005). The zebrafish shocked gene encodes a glycine transporter and is essential for the function of early neural circuits in the CNS. J. Neurosci. 25, 6610–6620. 10.1523/JNEUROSCI.5009-04.2005.

26. Bai, L., Zhang, Z., Ye, L., Cong, L., Zhao, Y., Zhang, T., Shi, Z., and Wang, K. (2022). Volumetric Imaging of Neural Activity by Light Field Microscopy. Neurosci. Bull. 38, 1559–1568. 10.1007/s12264-022-00923-9.

27. Wang, X., Allen, W.E., Wright, M.A., Sylwestrak, E.L., Samusik, N., Vesuna, S., Evans, K., Liu, C., Ramakrishnan, C., Liu, J., et al. (2018). Three-dimensional intact-tissue sequencing of single-cell transcriptional states. Science. 361. 10.1126/science.aat5691.

28. Alon, S., Goodwin, D.R., Sinha, A., Wassie, A.T., Chen, F., Daugharthy, E.R., Bando, Y., Kajita, A., Xue, A.G., Marrett, K., et al. (2021). Expansion sequencing: Spatially precise in situ transcriptomics in intact biological systems. Science. 371. 10.1126/science.aax2656.

29. Hildebrand, D.G.C., Cicconet, M., Torres, R.M., Choi, W., Quan, T.M., Moon, J., Wetzel, A.W., Scott Champion, A., Graham, B.J., Randlett, O., et al. (2017). Whole-brain serial-section electron microscopy in larval zebrafish. Nature 545, 345–349. 10.1038/nature22356.

30. Gautam, R., Hsu, N.C., Tsay, S.C., Lau, K.M., Holben, B., Bell, S., Smirnov, A., Li, C., Hansell, R., Ji, Q., et al. (2024). Predicting modular functions and neural coding of behavior from a synaptic wiring diagram. Nat. Neurosci. 8. 10.1038/s41593-024-01784-3.

31. Svara, F., Förster, D., Kubo, F., Januszewski, M., dal Maschio, M., Schubert, P.J., Kornfeld, J., Wanner, A.A., Laurell, E., Denk, W., et al. (2022). Automated synapse-level reconstruction of neural circuits in the larval zebrafish brain. Nat. Methods 19, 1357–1366. 10.1038/s41592-022-01621-0.

32. Petkova, M.D., Januszewski, M., Blakely, T., Herrera, K.J., Schuhknecht, G.F.P., Tiller, R., Choi, J., Schalek, R.L., Boulanger-Weil, J., Peleg, A., et al. (2025). A connectomic resource for neural cataloguing and circuit dissection of the larval zebrafish brain. bioRxiv, 2025.06.10.658982.

33. Li, F., Liu, J., Shi, C., Yuan, J., Lv, Y., Liu, J., and Zhang, L. (2025). Multi-level broad-yet-sparse input organization of LC-NE neurons revealed by multiplexed whole-brain EM reconstruction. bioRxiv, 2025.06.12.659365.

34. Vishwanathan, A., Daie, K., Ramirez, A.D., Lichtman, J.W., Aksay, E.R.F., and Seung, H.S. (2017). Electron Microscopic Reconstruction of Functionally Identified Cells in a Neural Integrator. Curr. Biol. 27, 2137–2147.e3. 10.1016/j.cub.2017.06.028.

35. Wanner, A.A., and Vishwanathan, A. (2018). Methods for Mapping Neuronal Activity to Synaptic Connectivity: Lessons From Larval Zebrafish. Front. Neural Circuits 12, 1–12. 10.3389/fncir.2018.00089.

36. Kunst, M., Laurell, E., Mokayes, N., Kramer, A., Kubo, F., Fernandes, A.M., Förster, D., Dal Maschio, M., and Baier, H. (2019). A Cellular-Resolution Atlas of the Larval Zebrafish Brain. Neuron 103, 21–38.e5. 10.1016/j.neuron.2019.04.034.

37. Wang, Y.-F., Chen, M.-Q., Wang, Y.-F., Chen, M.-Q., Ma, H.-B., Zhong, Y.-W., Jin, C.-X., Zhang, T.-L., Wang, K., Yue, Z.-F., et al. (2025). iBrAVE: a unified framework for 3D interactive and integrative analysis of brain atlas data across modalities and scales. bioRxiv. 10.1101/2025.10.04.680445.

38. Avants, B.B., Tustison, N.J., Song, G., Cook, P.A., Klein, A., and Gee, J.C. (2011). A reproducible evaluation of ANTs similarity metric performance in brain image registration. Neuroimage 54, 2033–2044. 10.1016/j.neuroimage.2010.09.025.

39. Randlett, O., Wee, C.L., Naumann, E.A., Nnaemeka, O., Schoppik, D., Fitzgerald, J.E., Portugues, R., Lacoste, A.M.B., Riegler, C., Engert, F., et al. (2015). Whole-brain activity mapping onto a zebrafish brain atlas. Nat. Methods 12, 1039–1046. 10.1038/nmeth.3581.

40. Marquart, G.D., Tabor, K.M., Brown, M., Strykowski, J.L., Varshney, G.K., LaFave, M.C., Mueller, T., Burgess, S.M., Higashijima, S., and Burgess, H. a. (2015). A 3D Searchable Database of Transgenic Zebrafish Gal4 and Cre Lines for Functional Neuroanatomy Studies. Front. Neural Circuits 9, 78. 10.3389/fncir.2015.00078.

41. Shainer, I., Kuehn, E., Laurell, E., Al Kassar, M., Mokayes, N., Sherman, S., Larsch, J., Kunst, M., and Baier, H. (2023). A single-cell resolution gene expression atlas of the larval zebrafish brain. Sci. Adv. 9, 1–10. 10.1126/sciadv.ade9909.

42. Němec, P., and Osten, P. (2020). The evolution of brain structure captured in stereotyped cell count and cell type distributions. Curr. Opin. Neurobiol. 60, 176–183. 10.1016/j.conb.2019.12.005.

43. Asakawa, K., and Kawakami, K. (2008). Targeted gene expression by the Gal4□UAS system in zebrafish. Dev. Growth Differ. 50, 391–399. 10.1111/j.1440-169X.2008.01044.x.

44. Marquart, G.D., Tabor, K.M., Horstick, E.J., Brown, M., Geoca, A.K., Polys, N.F., Nogare, D.D., and Burgess, H.A. (2017). High-precision registration between zebrafish brain atlases using symmetric diffeomorphic normalization. Gigascience 6, 1–15. 10.1093/gigascience/gix056.

45. Orger, M.B., Kampff, A.R., Severi, K.E., Bollmann, J.H., and Engert, F. (2008). Control of visually guided behavior by distinct populations of spinal projection neurons. Nat. Neurosci. 11, 327–333. 10.1038/nn2048.

46. Barker, A.J., and Baier, H. (2015). Sensorimotor decision making in the Zebrafish tectum. Curr. Biol. 25, 2804–2814. 10.1016/j.cub.2015.09.055.

47. Bianco, I.H., and Engert, F. (2015). Visuomotor transformations underlying hunting behavior in zebrafish. Curr. Biol. 25, 831–846. 10.1016/j.cub.2015.01.042.

48. Chen, X., Mu, Y., Hu, Y., Kuan, A.T., Nikitchenko, M., Randlett, O., Chen, A.B., Gavornik, J.P., Sompolinsky, H., Engert, F., et al. (2018). Brain-wide Organization of Neuronal Activity and Convergent Sensorimotor Transformations in Larval Zebrafish. Neuron 100, 876–890.e5. 10.1016/j.neuron.2018.09.042.

49. Privat, M., Romano, S.A., Pietri, T., Jouary, A., Boulanger-Weill, J., Elbaz, N., Duchemin, A., Soares, D., and Sumbre, G. (2019). Sensorimotor Transformations in the Zebrafish Auditory System. Curr. Biol. 29, 4010–4023.e4. 10.1016/j.cub.2019.10.020.

50. Lee, M.M., Arrenberg, A.B., and Aksay, E.R.F. (2015). A structural and genotypic scaffold underlying temporal integration. J. Neurosci. 35, 7903–7920. 10.1523/JNEUROSCI.3045-14.2015.

51. Koyama, M., Kinkhabwala, A., Satou, C., Higashijima, S.I., and Fetcho, J. (2011). Mapping a sensory-motor network onto a structural and functional ground plan in the hindbrain. Proc. Natl. Acad. Sci. U. S. A. 108, 1170–1175. 10.1073/pnas.1012189108.

52. Costa, M., Manton, J.D., Ostrovsky, A.D., Prohaska, S., and Jefferis, G.S.X.E. (2016). NBLAST: Rapid, Sensitive Comparison of Neuronal Structure and Construction of Neuron Family Databases. Neuron 91, 293–311. 10.1016/j.neuron.2016.06.012.

53. Zeileis, A., Leisch, F., Homik, K., and Kleiber, C. (2002). strucchange: An R Package for Testing for Structural Change. J. Stat. Softw. 7, 1–38.

54. Zeileis, A., Kleiber, C., Walter, K., and Hornik, K. (2003). Testing and dating of structural changes in practice. Comput. Stat. Data Anal. 44, 109–123. 10.1016/S0167-9473(03)00030-6.

55. Straka, H., Simmers, J., and Chagnaud, B.P. (2018). A New Perspective on Predictive Motor Signaling. Curr. Biol. 28, R232–R243. 10.1016/j.cub.2018.01.033.

56. Wang, W.C., and McLean, D.L. (2014). Selective Responses to Tonic Descending Commands by Temporal Summation in a Spinal Motor Pool. Neuron 83, 708–721. 10.1016/j.neuron.2014.06.021.

57. Schoppik, D., Bianco, I.H., Prober, D.A., Douglass, A.D., Robson, D.N., Li, J.M.B., Greenwood, J.S.F., Soucy, E., Engert, F., and Schier, A.F. (2017). Gaze-stabilizing central vestibular neurons project asymmetrically to extraocular motoneuron pools. J. Neurosci. 37, 11353–11365. 10.1523/JNEUROSCI.1711-17.2017.

58. Hellwig, B. (2000). A quantitative analysis of the local connectivity between pyramidal neurons in layers 2/3 of the rat visual cortex. Biol. Cybern. 82, 111–121. 10.1007/PL00007964.

59. Shepherd, G.M.G., Stepanyants, A., Bureau, I., Chklovskii, D., and Svoboda, K. (2005). Geometric and functional organization of cortical circuits. Nat. Neurosci. 8, 782–790. 10.1038/nn1447.

60. Cook, S.J., Kalinski, C.A., and Hobert, O. (2023). Neuronal contact predicts connectivity in the C. elegans brain. Curr. Biol. 33, 2022.11.13.516316. 10.1016/j.cub.2023.04.071.

61. Gao, L., Liu, S., Wang, Y., Wu, Q., Gou, L., and Yan, J. (2023). Single-neuron analysis of dendrites and axons reveals the network organization in mouse prefrontal cortex. Nat. Neurosci. 26, 1111–1126. 10.1038/s41593-023-01339-y.

62. Xiong, F., Liu, L., and Peng, H. (2025). Reconstruction of a connectome of single neurons in mouse brains by cross-validating multi-scale multi-modality data. Nat. Methods 22, 2670–2683. 10.1038/s41592-025-02784-2.

63. Helmbrecht, T.O., dal Maschio, M., Donovan, J.C., Koutsouli, S., and Baier, H. (2018). Topography of a Visuomotor Transformation. Neuron 100, 1429–1445.e4. 10.1016/j.neuron.2018.10.021.

64. Zhang, B. bing, Yao, Y. yuan, Zhang, H. fei, Kawakami, K., and Du, J. lin (2017). Left Habenula Mediates Light-Preference Behavior in Zebrafish via an Asymmetrical Visual Pathway. Neuron 93, 914–928.e4. 10.1016/j.neuron.2017.01.011.

65. Cheng, R.K., Krishnan, S., Lin, Q., Kibat, C., and Jesuthasan, S. (2017). Characterization of a thalamic nucleus mediating habenula responses to changes in ambient illumination. BMC Biol. 15, 1–21. 10.1186/s12915-017-0431-1.

66. Lau, B.Y.B., Mathur, P., Gould, G.G., and Guo, S. (2011). Identification of a brain center whose activity discriminates a choice behavior in zebrafish. Proc. Natl. Acad. Sci. U. S. A. 108, 2581–2586. 10.1073/pnas.1018275108.

67. Bullmore, E., and Sporns, O. (2009). Complex brain networks: Graph theoretical analysis of structural and functional systems. Nat. Rev. Neurosci. 10, 186–198. 10.1038/nrn2575.

68. Luo, L. (2021). Architectures of neuronal circuits. Science. 373. 10.1126/science.abg7285.

69. Milo, R., Shen-Orr, S., Itzkovitz, S., Kashtan, N., Chklovskii, D., and Alon, U. (2002). Network Motifs: Simple Building Blocks of Complex Networks. Science. 298, 824–827. 10.1126/science.298.5594.824.

70. Maaten, L. van der, and Hinton, G. (2008). Visualizing Data using t-SNE. J. Mach. Learn. Res. 9, 2579–2605.

71. Loring, M.D., Thomson, E.E., and Naumann, E.A. (2020). Whole-brain interactions underlying zebrafish behavior. Curr. Opin. Neurobiol. 65, 88–99. 10.1016/j.conb.2020.09.011.

72. Wang, Q., Ding, S.L., Li, Y., Royall, J., Feng, D., Lesnar, P., Graddis, N., Naeemi, M., Facer, B., Ho, A., et al. (2020). The Allen Mouse Brain Common Coordinate Framework: A 3D Reference Atlas. Cell 181, 936–953.e20. 10.1016/j.cell.2020.04.007.

73. Spencer, N.J., and Hu, H. (2020). Enteric nervous system: sensory transduction, neural circuits and gastrointestinal motility. Nat. Rev. Gastroenterol. Hepatol. 17, 338–351. 10.1038/s41575-020-0271-2.

74. Filippi, A., Mueller, T., and Driever, W. (2014). vglut2 and gad expression reveal distinct patterns of dual GABAergic versus glutamatergic cotransmitter phenotypes of dopaminergic and noradrenergic neurons in the zebrafish brain. J. Comp. Neurol. 522, 2019–2037. 10.1002/cne.23524.

75. Higashijima, S.I., Mandel, G., and Fetcho, J.R. (2004). Distribution of prospective glutamatergic, glycinergic, and gabaergic neurons in embryonic and larval zebrafish. J. Comp. Neurol. 480, 1–18. 10.1002/cne.20278.

76. Kidwell, C.U., Su, C.Y., Hibi, M., and Moens, C.B. (2018). Multiple zebrafish atoh1 genes specify a diversity of neuronal types in the zebrafish cerebellum. Dev. Biol. 438, 44–56. 10.1016/j.ydbio.2018.03.004.

77. Raj, B., Wagner, D.E., McKenna, A., Pandey, S., Klein, A.M., Shendure, J., Gagnon, J.A., and Schier, A.F. (2018). Simultaneous single-cell profiling of lineages and cell types in the vertebrate brain. Nat. Biotechnol. 36, 442–450. 10.1038/nbt.4103.

78. Svensson, E., Apergis-Schoute, J., Burnstock, G., Nusbaum, M.P., Parker, D., and Schiöth, H.B. (2019). General principles of neuronal co-transmission: Insights from multiple model systems. Front. Neural Circuits 12, 1–28. 10.3389/fncir.2018.00117.

79. Tay, T.L., Ronneberger, O., Ryu, S., Nitschke, R., and Driever, W. (2011). Comprehensive catecholaminergic projectome analysis reveals single-neuron integration of zebrafish ascending and descending dopaminergic systems. Nat. Commun. 2, 112–171. 10.1038/ncomms1171.

80. Chen, A., Chiu, C.N., Mosser, E.A., Kahn, S., Spence, R., and Prober, D.A. (2016). QRFP and its receptors regulate locomotor activity and sleep in zebrafish. J. Neurosci. 36, 1823–1840. 10.1523/JNEUROSCI.2579-15.2016.

81. Herget, U., Gutierrez-Triana, J.A., Thula, O.S., Knerr, B., and Ryu, S. (2017). Single-cell reconstruction of oxytocinergic neurons reveals separate hypophysiotropic and encephalotropic subtypes in larval zebrafish. eNeuro 4, 1–16. 10.1523/ENEURO.0278-16.2016.

82. Lovett-Barron, M., Chen, R., Bradbury, S., Andalman, A.S., Wagle, M., Guo, S., and Deisseroth, K. (2020). Multiple convergent hypothalamus–brainstem circuits drive defensive behavior. Nat. Neurosci. 23, 959–967. 10.1038/s41593-020-0655-1.

83. Yokogawa, T., Hannan, M.C., and Burgess, H.A. (2012). The dorsal raphe modulates sensory responsiveness during arousal in zebrafish. J. Neurosci. 32, 15205–15215. 10.1523/JNEUROSCI.1019-12.2012.

84. Huang, K.W., Ochandarena, N.E., Philson, A.C., Hyun, M., Birnbaum, J.E., Cicconet, M., and Sabatini, B.L. (2019). Molecular and anatomical organization of the dorsal raphe nucleus. Elife 8, 1–34. 10.7554/eLife.46464.

85. Güntürkün, O., Ströckens, F., and Ocklenburg, S. (2020). Brain lateralization: A comparative perspective. Physiol. Rev. 100, 1019–1063. 10.1152/physrev.00006.2019.

86. Miletto Petrazzini, M.E., Sovrano, V.A., Vallortigara, G., and Messina, A. (2020). Brain and Behavioral Asymmetry: A Lesson From Fish. Front. Neuroanat. 14, 1–22. 10.3389/fnana.2020.00011.

87. Sun, W., Ripp, I., Borrmann, A., Moll, M., and Fairhurst, M. (2025). Touch-driven advantages in reaction time but not in performance in a cross-sensory comparison of reinforcement learning. Heliyon 11, e41330. 10.1016/j.heliyon.2024.e41330.

88. Gouwens, N.W., Sorensen, S.A., Baftizadeh, F., Budzillo, A., Lee, B.R., Jarsky, T., Alfiler, L., Baker, K., Barkan, E., Berry, K., et al. (2020). Integrated Morphoelectric and Transcriptomic Classification of Cortical GABAergic Cells. Cell 183, 935–953.e19. 10.1016/j.cell.2020.09.057.

89. Dohaku, R., Yamaguchi, M., Yamamoto, N., Shimizu, T., Osakada, F., and Hibi, M. (2019). Tracing of afferent connections in the zebrafish cerebellum using recombinant rabies virus. Front. Neural Circuits 13, 1–15. 10.3389/fncir.2019.00030.

90. Kler, S., Ma, M., Narayan, S., Ahrens, M.B., and Albert, Y. (2021). Cre-Dependent Anterograde Transsynaptic Labeling and Functional Imaging in Zebrafish Using VSV With Reduced Cytotoxicity. Front. Neuroanat. 15, 1–10. 10.3389/fnana.2021.758350.

91. Satou, C., Neve, R.L., Oyibo, H.K., Zmarz, P., Huang, K.-H., Bouldoires, E.A., Mori, T., Higashijima, S.I., Keller, G.B., and Friedrich, R.W. (2022). A viral toolbox for conditional and transneuronal gene expression in zebrafish. Elife 11, 1–24. 10.7554/ELIFE.77153.

92. Chen, T.-L., Deng, Q.-S., Lin, K.-Z., Zheng, X.-D., Wang, X., Zhong, Y.-W., Ning, X.-Y., Li, Y., Xu, F.-Q., Du, J.-L., et al. (2024). An applicable and efficient retrograde monosynaptic circuit mapping tool for larval zebrafish, 10.7554/eLife.100880.1 10.7554/eLife.100880.1.

93. McLean, D.L., Fan, J., Higashijima, S.I., Hale, M.E., and Fetcho, J.R. (2007). A topographic map of recruitment in spinal cord. Nature 446, 71–75. 10.1038/nature05588.

94. Suster, M.L., Abe, G., Schouw, A., and Kawakami, K. (2011). Transposon-mediated BAC transgenesis in zebrafish. Nat. Protoc. 6, 1998–2021. 10.1038/nprot.2011.416.

95. Satou, C., Kimura, Y., and Higashijima, S. ichi (2012). Generation of multiple classes of V0 neurons in Zebrafish spinal cord: Progenitor heterogeneity and temporal control of neuronal diversity. J. Neurosci. 32, 1771–1783. 10.1523/JNEUROSCI.5500-11.2012.

96. Yao, Y., Li, X., Zhang, B., Yin, C., Liu, Y., Chen, W., Zeng, S., and Du, J. (2016). Visual Cue-Discriminative Dopaminergic Control of Visuomotor Transformation and Behavior Selection. Neuron 89, 598–612. 10.1016/j.neuron.2015.12.036.

97. Balciunas, D., Wangensteen, K.J., Wilber, A., Bell, J., Geurts, A., Sivasubbu, S., Wang, X., Hackett, P.B., Largaespada, D.A., McIvor, R.S., et al. (2006). Harnessing a high cargo-capacity transposon for genetic applications in vertebrates. PLoS Genet 2, e169.

98. Köster, R.W., and Fraser, S.E. (2001). Tracing transgene expression in living zebrafish embryos. Dev. Biol. 233, 329–346. 10.1006/dbio.2001.0242.

99. Meyer, M.P., and Smith, S.J. (2006). Evidence from In Vivo Imaging That Synaptogenesis Guides the Growth and Branching of Axonal Arbors by Two Distinct Mechanisms. J. Neurosci. 26, 3604–3614. 10.1523/JNEUROSCI.0223-06.2006.

100. Avants, B.B., Yushkevich, P., Pluta, J., Minkoff, D., Korczykowski, M., Detre, J., and Gee, J.C. (2010). The optimal template effect in hippocampus studies of diseased populations. Neuroimage 49, 2457–2466. 10.1016/j.neuroimage.2009.09.062.

101. Rohlfing, T. (2012). Image Similarity and Tissue Overlaps as Surrogates for Image Registration Accuracy: Widely Used but Unreliable. IEEE Trans. Med. Imaging 31, 153–163. 10.1109/TMI.2011.2163944.

102. Arshadi, C., Günther, U., Eddison, M., Harrington, K.I.S., and Ferreira, T.A. (2021). SNT: a unifying toolbox for quantification of neuronal anatomy. Nat. Methods 18, 374–377. 10.1038/s41592-021-01105-7.

103. Cannon, R.C., Turner, D.A., Pyapali, G.K., and Wheal, H. V. (1998). An on-line archive of reconstructed hippocampal neurons. J. Neurosci. Methods 84, 49–54. 10.1016/S0165-0270(98)00091-0.

104. Huttenlocher, D.P., Klanderman, G.A., and Rucklidge, W.J. (1993). Comparing images using the Hausdorff distance. IEEE Trans. Pattern Anal. Mach. Intell. 15, 850–863. 10.1109/34.232073.

105. Sander, J., Ester, M., Kriegel, H.P., and Xu, X. (1998). Density-based clustering in spatial databases: The algorithm GDBSCAN and its applications. Data Min. Knowl. Discov. 2, 169–194. 10.1023/A:1009745219419.

106. Oh, S.W., Harris, J.A., Ng, L., Winslow, B., Cain, N., Mihalas, S., Wang, Q., Lau, C., Kuan, L., Henry, A.M., et al. (2014). A mesoscale connectome of the mouse brain. Nature 508, 207–214. 10.1038/nature13186.

107. Page, L., Brin, S., Motwani, R., and Winograd, T. (1999). The PageRank Citation Ranking: Bringing Order to the Web. Tech. Report. Stanford InfoLab.

