## Supplementary information for "Nervous system-wide single-cell morphology atlas of excitatory and inhibitory neurons in larval zebrafish"

Contents：

Figures S1-S8

Tables S1-S3

Videos S1-S5

**SUPPLEMENTAL FIGURE LEGENDS**

**Figure S1. A web portal for exploring all datasets in the CPSv1, related to Figure 1**

(A) Interactive panel for selecting aligned line image datasets from ZExplorer, Z-Brain, ZBB, and mapzebrain to display in the 3D viewer.

(B) Interactive panel for selecting interested neuron tracings (left) and corresponding 3D view showing both tracings and anatomical regions (right). Users can query neurons based on soma and axon position using “overlap/exclude/AND” operations, and adjust color or visibility for individual or grouped tracings using the “item in the scene” panel.

(C) Interactive panel for selecting anatomical regions in the CPSv1 to display in the 3D viewer.

(D) Interactive panel for selecting cytoarchitecture data of specific cell populations for visualization.

(E) Neuron tracings visualized with different color schemes for cellular compartments (soma, dendrite, axon; menu toggled by clicking the red-boxed icon in (B).

(F) Example of an integrated 3D view for the combination of >10,000 neuron tracings, brain regions, and cytoarchitectures of *vglut2a*, *vglut2b*, *gad1b*, and *glyt2* neurons.

(G) Example analysis outputs from the online analysis functions computed based on selected neuronal tracings, including soma distribution along the *x*, *y*, and *z* axes (top left), total projection length across regions (top right), interregional projection heatmap (bottom left and right). Icons for accessing analysis modules are shown on the left.

See also <https://zebrafish.cn/LM-Atlas/EI>.

**Figure S2. Template creation and inter-atlas registration accuracy evaluation, related to Figure 1**

(A) Iterative template refinement. Two cropped areas (1,2) from a horizontal section (left) show increased anatomical clarity in the average images (right) with iteration. Area 1: *elavl3:H2B-GCaMP6s* pattern around region boundaries; Area 2: *vglut2a:DsRed* pattern in neuronal fiber-enriched regions.

(B) Improved similarity measured by normalized cross correlation (NCC) between successive iterations, showing convergence to a stable average template.

(C) Summary of lines, fish numbers, iteration counts, and bridging lines for generating the original dual-channel template (*elavl3:H2B-GCaMP6s;vglut2a:DsRed*) and six additional reference templates (in bold).

(D) Volume rendering (dorsal view) of five additional reference templates in the CPSv1. Partial spinal cord shown.

(E–G) Horizontal sections showing merged images of our bridge template (ZExplorer, green) and the registered counterpart from another atlas (red). The notation above each image indicates the name of the bridge template and its source atlas (as superscripts). The “to” denotes the registration direction between two atlases. Specifically, Z-Brain (E) was registered via the *elavl3:H2B-RFP* template to our ZExplorer *elavl3:H2B-GCaMP6*s template, ZBB (F) and mapzebrain (G) were registered via the *vglut2a:DsRed* template to our corresponding *vglut2a:DsRed* template.

(H) Point-based landmarks identified in the template counterparts across ZExplorer and three other atlases, with their coordinates reported in the CPSv1 reference frame and mean registration precision indicated. A total of six expression patterns were used, comprising 11 pairs of comparisons, with each pattern indicated by a distinct background color. The name of each template counterpart for each expression pattern is shown, with its source atlas indicated as a superscript.

(I) Landmark positions for each expression pattern (H) overlaid on the horizontal (left) and sagittal (right) maximum projections of *elavl3:H2B-GCaMP6s*. Coloring scheme follows that in (H).

(J) Distribution of landmark distances (LDs) across 86 inter-atlas landmark pairs (26 from 3 expression patterns for Z-Brain *vs*. ZExplorer; 35 from 5 expression patterns for ZBB *vs*. ZExplorer; 25 from 3 expression patterns for mapzebrain *vs*. ZExplorer).

(K) Summary of LDs for pairwise expression pattern comparisons (left *y* axis; colored bars and circles; coloring scheme follows that in (H)) and for atlas pairs (right *y* axis; purple dots with error bars) used to evaluate inter-atlas registration precision. ZExplorer *vs*. Z-brain and mapzebrain: *elavl3:H2B-FPs*, *elavl3:GECIs*, and *vglut2a:DsRed*; ZExplorer *vs*. ZBB: *elavl3:GECIs*, *vglut2a:DsRed*, *gad1b:GFP*, *glyt2:GFP*, and *vmat2:GFP*.

Scale bars: 50 μm (A, left), 20 μm (A, right), 100 μm (D–G, I). A, anterior; D, dorsal; V, ventral.

**Figure S3. Evaluation of registration precision and neuron sampling randomness for the single E and I neuron atlas, related to Figure 3**

(A) Thirteen point-based landmarks identified in the *elavl3:H2B-GCaMP6s* reference template for evaluating registration precision. Coordinates refer to positions on sagittal (*x*), transverse (*y*), and horizontal (*z*) planes in the CPSv1. LD (mean ± SEM) represents average registration precision for each landmark from 88 randomly selected individual fish.

(B) Horizontal sections showing landmark positions (cross center) in the *elavl3:H2B-GCaMP6s* expression pattern.

(C) Spatial distribution of the 13 landmarks overlaid on sagittal (top) and horizontal (bottom) maximum projections of the *elavl3:H2B-GCaMP6s* reference template.

(D) Distribution of 1,101 LDs from 88 individual fish, with less than 13 landmarks detected in some fish.

(E) Boxplot of LDs for each of the 13 landmarks, with 78–88 fish per landmark. The central mark, bottom, and top of the boxplot indicate the median, first, and third quartiles, respectively.

(F) Distribution of MLDs across 88 fish. Red dashed line indicates the overall mean value.

(G) Registration of Mauthner cells in two fish backfilled by spinal dye injection (reference fish, pseudocolored in yellow; moving fish, pseudocolored in magenta; see Methods). Pan-neuronal labeling by *elavl3:H2B-GCaMP6s* (reference fish, pseudocolored in blue; moving fish, pseudocolored in green) was used to guide registration. Asterisks mark individual left Mauthner cells used for quantifying registration precision.

(H) Reconstructed Mauthner cells (n = 14) in the single-cell morphology atlas, used for registration precision evaluation. Left, dorsal view; Right, lateral view.

(I) Summary of distances between corresponding single Mauthner cells across registered fish pairs (see Methods). Mauthner cells were labeled by spinal backfill (N = 9 larvae; examples shown in G), or reconstructed in this study shown in (H). Numbers in parentheses represent the number of cell pairs.

(J–L) Volume rendering (dorsal view) of somata from reconstructed *vglut2a/2b* (J), *gad1b* (K), and *glyt2* (L) neurons in the CPSv1. Sphere size reflects soma diameter measured. Partial spinal cord shown.

(M–O) Bar plots comparing the regional distributions of complete (light color) and sampled (dark color) neurons across 52 anatomical regions (with subregions in the spinal cord and vagal ganglia merged) for each E and I neuron populations.

(P) Scatterplot showing the average Pearson correlation between the regional distributions of complete and randomly sampled neurons (1,000 times) from E and I neuron populations, with sampling size increasing from 1 cell to ~50% of the total by one-cell increment. Dotted line indicates the Pearson’s *r* at the maximum sampling size.

Scale bars: 20 μm (B and G), 50 μm (H), 100 μm (C, J–L). A, anterior; D, dorsal; V, ventral.

**Figure S4. Dendrite-axon identification and validation of projection strength metrics, related to Figures 4**

(A, C, and E) Volume rendering (dorsal view) of image stacks (left) with a single *vglut2a* (A), *gad1b* (C), or *glyt2* (E) neuron labeled by *Sypb-EGFP-tdKatushka2-CAAX* and corresponding reconstructions (right) showing soma (cyan), dendrites (blue), axons (red), and presynaptic puncta (yellow).

(B, D, and F) Dorsal (left) and lateral (right) 3D views of single-neuron reconstructions for *vglut2a* (B), *gad1b* (D), and *glyt2* (F) neurons labeled by *Sypb-EGFP-tdKatushka2-CAAX*, with sub-structures color-coded as in (A). Partial neurons are shown for clarity.

(G–I) Machine learning-based dendrite-axon classification for three known cell types: mitral cells (G), habenular neurons (H), and Mauthner cells (I), with the classification accuracy indicated by the percentage. The background is the brain regions where the soma locates. Soma, yellow; dendrite, blue; axon, red.

(J and K) Neuron-to-region projection matrices generated using two metrics: presynaptic site count (J) and terminal-bearing axon length (K). These two metrics display high similarity (Pearson’s *r* = 0.78, *P* < 0.001), analyzed by *corrcoef* function.

Scale bars: 100 μm (A–F), 50 μm (G–I). A, anterior; D, dorsal; V, ventral. Numbers in parentheses indicate the number of neurons examined.

**Figure S5. Overview of E and I morphotypes, related to Figures 5-7**

(A and B) Volume rendering (dorsal view) of 348 excitatory (A), 164 GABAergic, and 88 glycinergic inhibitory (B) morphological subtypes displayed in ontological order. In (B), GABAergic and glycinergic neurons are displayed in alternating rows, with vertically aligned positions indicating corresponding inhibitory morphotypes used to generate the inter-cell-type putative connectome shown in Figures 7 and S7H. Each subtype ID (up to 11 characters) encodes its neurotransmitter identity, division (1–5), category (A–N), class (two-digit numbers), type (two-digit numbers), and subtype (a lowercase letter). Morphotypes used for dendrite-axon classification accuracy assessment (see Results) are indicated with a gray background in the ID.

Most morphotypes in the “Local” category (KI) lack dendrite and axon annotations. Scale bar: 100 μm. KA–KN denote 14 categories. A, anterior; V, ventral. HSB, hindbrain-spinal cord boundary; MHB, midbrain-hindbrain boundary; TDB, telencephalon-diencephalon boundary.

**Figure S6. Morphological classification of I neurons and cell type coverage assessment, related to Figure 5**

(A) Dendrogram of GABAergic (*gad1b*) inhibitory neurons comprising 4 divisions, 9 categories, 36 classes, and 155 types, with bar plots showing cell number and regional distribution.

(B) Dendrogram of glycinergic (*glyt2*) inhibitory neurons comprising 2 divisions, 7 categories, 19 classes, and 84 types, with bar plots showing cell number and regional distribution.

Color schemes for brain regions are shown in the bottom right corner.

(C–E) Scatterplot showing the cell type distribution dissimilarity (left *y*-axis; gray dots, *P* < 0.05; black dots, *P* > 0.05; maximum *Χ*^2^ statistic) between randomly sampled and full classified populations, and the cell type coverage (right *y*-axis, color-coded dots, minimum value), based on 1000 randomly selected cell groups. Sampling sizes with a 10-cell step range from 1 to the total classified cells for *vglut2a/2b* (C, 9,774 neurons), *gad1b* (D, 3,411 neurons), and *glyt2* (E, 1,116 neurons) populations. Dashed red line, sampling size at the breakpoint; dashed black line, sampling size at which 95% cell type coverage is reached.

**Figure S7. Morphotype-to-region and putative inter-morphotype connectivity matrices, related to Figures 6 and 7**

(A–C) Dendritic recipient and axonal projection matrices of E and I morphological subtypes. Quantitative recipient (left), ipsilateral projection (middle), and contralateral projection (right) strengths of each morphological subtype in *vglut2a/2b* (A, 348 subtypes), *gad1b* (B, 164 subtypes), and *glyt2* (C, 88 subtypes) neurons across 68 anatomical regions. Regions are ordered by primary subdivisions (color-coded; see Figure 2) and then by anteroposterior centroid. Morphotypes are displayed in ontological order, and KA–KN indicate 14 categories. Matrices show log_10_-transformed strengths normalized by the number of cells. For subtypes without dendrite-axon annotations (see Figure S5), total neurites were analyzed and are presented in the axon projection section.

(D and E) Pairwise similarity (Pearson’s *r*) of neuron-level (D) and type-level (E) putative connectivity matrices across dendrite-axon distance thresholds of 0.5, 1, 2, 3, and 4 µm (computed between axonal and dendritic branch segment centers; see Methods), with all correlations significant (all *P* < 0.001).

(F) Mean pairwise similarity (Pearson’s *r*) of each threshold to all others, derived from the similarity matrices in (D) and (E), peaking at 2 µm at both neuron and type levels.

(G) Putative inter-cell-type mesoscale connectivity matrix. Directed and weighted connection strengths between 475 types (n_E_ = 295, n_I_ = 180; excluding those without dendrite-axon annotations), displayed by E and I groups and in ontological order. Matrix values represent the pairwise mesoscale connection strength from row to column types, normalized by the number of connected neuron pairs. Connections were filtered by requiring neuronal participation ≥20% in at least one member of each connected morphotype pair, and both divergence and convergence indices >1, resulting in 32,803 directed connections

**Figure S8. Analysis of hubs and information flow in the interregional network, related to Figure 8**

(A) Nine network metrics, including input strength, output strength, input degree, output degree, weighted betweenness centrality, weighted closeness, page rank, clustering, and vitality. Only E network-relevant data are shown.

(B) t-SNE plots of regions, based on nine network metrics. Top, E (*vglut2a*/*2b*); upper middle, I (*gad1b* and *glyt2*); lower middle, *gad1b*; bottom, *glyt2*. Perplexity = 20. Regions are color-coded by hub score. Red star indicates hub regions with above-mean values across all metrics, and blue crosses mark regions with the highest score in any single metric (with overlaps across some metrics). The top three hub regions (highest overall hub scores) are labeled in red.

(C) Regional input-output diagrams of the top three hub regions for *vglut2a/2b* (top; R8, R1, and TeO), *gad1b* (middle; R1, R8, and TeO), and *glyt2* (bottom; R8, R6, and R7) neuron networks. Parabolas represent directed ipsilateral (red) and contralateral (blue) connections between regions, which are arranged linearly based on their anteroposterior centroid coordinates. Right parabolas indicate descending connections, while left parabolas indicate ascending connections. The parabola’s height and color tone both represent macroscale connection strength.

(D and E) Inhibitory hub networks constructed from *gad1b* (D) or *glyt2* (E) neurons. Each node (sphere) represents one of 53 regions (retina excluded), with its color and size indicating its hub score. The color and width (normalized betweenness centrality) of the edge both reflect its importance in the information flow among regions. The names of the top three hub regions are in red.

A, anterior; D, dorsal; V, ventral.

**SUPPLEMENTAL VIDEOS**

**Video S1. Common physical space (CPSv1): Composite horizontal tomography of 7 transgenic reference templates and radial disassembly of 173 delineated parcellations in the CPSv1, related to Figure 1.**

**Video S2. Pipeline for building digital cytoarchitecture atlases of E and I neurons in 6-dpf larval zebrafish, related to Figure 2.**

**Video S3. Single-cell morphology atlases of E and I neurons: Pipeline for building digital single-cell morphology atlases and single-cell morphology atlases of E and I neurons in 6-dpf larval zebrafish, related to Figure 3.**

**Video S4. Morphotype-based visualization of the visuomotor pathways: The visuomotor pathway from RGCs to motor neurons and the visuomotor pathway underlying light-preference behaviors, related to Figure 7.**

**Video S5. 3D rotation of the nervous system-wide hub networks shown in Figures 8J, S8D, and S8E constructed based on E, I, *gad1b*+, or *glyt2*+ single-neuron morphology atlas, respectively, related to Figure 8.**

**SUPPLEMENTAL TABLES**

**Table S1**. **Information of incorporated reference templates from Z-Brain (24), ZBB (245), and mapzebrain (428), related to Figure 1.**

See spreadsheet Table S1.

**Table S2. Anatomical parcellations** **in the zebrafish CPSv1, related to F****igure 1.**

See spreadsheet Table S2.

**Table S3. Information of BAC constructs, related to Figure 1.**

| BAC  construct | BAC clone name and source | iTol2 primers | Recombination arm primers | Replaced site |
| --- | --- | --- | --- | --- |
| *vglut2b:GAL4FF* | CH211**-**5O13 (BACPAC) | pTARBAC_iTol2_F (5′–3′):  GCGTAAGCGGGGCACATTTCATTACCTCTTTCTCCGCACCCGACATAGATCCCTGCTCGAGCCGGGCCCAAGTG  pTARBAC_iTol2_R (5′–3′):  CGCGGGGCATGACTATTGGCGCGCCGGATCGATCCTTAATTAAGTCTACTAATTATGATCCTCTAGATCAGATCT | *vglut2b*-F (5′–3′): CATCGTCTTCTTGCCATT  *vglut2b*-R (5′–3′): GATTTTCTCGTCATGAAT | Exon I |
| *gad1b:GAL4-VP16* | DKEY-251E16 (BioScience) | pIndigoBAC_iTol2_F (5′–3′):  TTCTCTGTTTTTGTCCGTGGAATGAACAATGGAAGTCCGAGCTCATCGCTCCCTGCTCGAGCCGGGCCCAAGTG  pIndigoBAC_iTol2_R (5′–3′):  CCCGCCAACACCCGCTGACGCGAACCCCTTGCGGCCGCATATTATGATCCTCTAGATCAGATCT | *gad1b*-F (5′–3′): CATGAGCGGCGCAATGAT  *gad1b*-R (5′–3′): CGCCTGTGTAAAAGCGCA | Exon I and 739 bp of intron I |
